# Land use and climate warming interact to reshape global avian biodiversity through ecological filtering and homogenisation

**DOI:** 10.64898/2026.10.08.757631

**Authors:** Chenchen Ding, Tim Newbold, Adrienne Etard, Richard Cornford, Martin Jung, Kyle J. Brumm, Zhigang Jiang, Piero Visconti, Zhiheng Wang

**Author notes:** Corresponding authors: Chenchen Ding, Piero Visconti and Zhiheng Wang.

## Abstract

Land-use and climate change are dominant drivers of terrestrial biodiversity change, yet are usually analysed separately although they co-occur and may interact. We integrated global bird assemblage records from PREDICTS with temperature anomalies and species’ ecological characteristics to test how land use modifies biodiversity responses to climate warming across three dimensions: local abundance and richness, compositional dissimilarity among assemblages, and species occurrence across ecological groups. Land use modified warming associations for both abundance and richness. Relative to primary vegetation, warming associations were more positive in low-intensity agriculture for both responses, but more negative for richness in high-intensity agriculture and urban land. Predicted richness remained below the reference, defined as primary vegetation at temperature anomaly 0, across all modified land uses, with the largest declines in high-intensity agriculture (−17.9%) and urban land (−21.8%). Within land-use classes, greater warming exposure was generally associated with higher compositional similarity, consistent with warming-associated biotic homogenisation. Occurrence models revealed ecological filtering: broad-niched generalists were favoured in warmer modified habitats, whereas range-restricted habitat specialists were filtered out under intensive land use and warming. Conservation assessments should therefore integrate land-use– warming interactions with changes in assemblage composition and ecological strategies, particularly in agricultural and urban landscapes where warming further disadvantaged range-restricted specialists.

---

Biodiversity is being reorganised by multiple anthropogenic pressures that increasingly operate in concert rather than in isolation^1,2^. Land-use change and climate change are among the strongest drivers of terrestrial biodiversity change, contributing to species loss and shifts in species distributions, phenology and community structure^3–9^. Yet the two drivers are still often studied separately or treated as additive pressures^10–12^, even though each can modify responses to the other^13–17^.

Two mechanisms make non-additive responses possible. First, land-use conversion and intensification reshape habitat resources and connectivity, filtering species unable to persist in human-modified conditions and constraining climate-driven range shifts^14,18^. Second, the conversion also alters microclimate, for example through reduced canopy cover and moisture, amplifying exposure to heat under warming^19,20^. Joint effects are therefore expected to be synergistic (amplifying biodiversity losses) or antagonistic (partially offsetting impacts) rather than additive^15,16,21^.

Evidence for such interactions is accumulating^22–25^, but their direction and magnitude remain hard to generalise. For example, high-intensity agriculture combined with strong warming was associated with substantially lower insect abundance and richness worldwide^26^. For birds, assemblage change in Costa Rica was greatest in intensive agriculture under hotter, drier conditions^27^, and the two drivers have jointly driven homogenisation across British taxa^28^, population declines in North America^29^, and both concordant and opposing effects on species occupancy in California^21^. Ignoring the interactive effects of climate change and land-use change risks misattributing biodiversity change, biasing projections and misdirecting conservation efforts^22,30^.

Three main gaps constrain ecological understanding of global change. First, only a few analyses have explicitly tested how the two drivers interact at a global scale^26,31,32^, with most evidence coming from particular regions and environmental contexts. Second, biodiversity change is often diagnosed from species richness or abundance alone, which misses shifts in identity and dominance when losses of vulnerable species are offset by gains of disturbance-tolerant species^8,33–35^. Third, species responses are structured by ecological characteristics: restricted ranges, narrow thermal niches and limited dispersal confer vulnerability, whereas widespread, broad-niched and mobile species more often persist^36–38^. These characteristics are not independent but covary along axes of morphology, life history and dispersal^39,40^. Species may therefore respond to land use and warming through combinations of characteristics rather than any single one, but whether such combinations identify consistent winners and losers under combined land-use and climate pressures has rarely been tested. Birds provide a suitable system for addressing these gaps: they are globally distributed, respond sensitively to both drivers^38,41^, have rich assemblage and trait data^40,42^, and provide essential ecosystem services^43,44^.

Here we tested whether land use and climate change interact to reshape avian biodiversity globally (Fig. 1). We combined bird assemblage records from PREDICTS^42,45,46^ with harmonised land-use classes, standardised temperature anomalies from CRU TS^47^ (Extended Data Fig. 1) and ecological characteristics^40^. We evaluated joint effects of land-use and climate change across three dimensions of ecological organisation: local abundance and species richness; pairwise dissimilarity in assemblage composition; and species-level occurrence among groups defined by ecological characteristics.

**Fig. 1.**
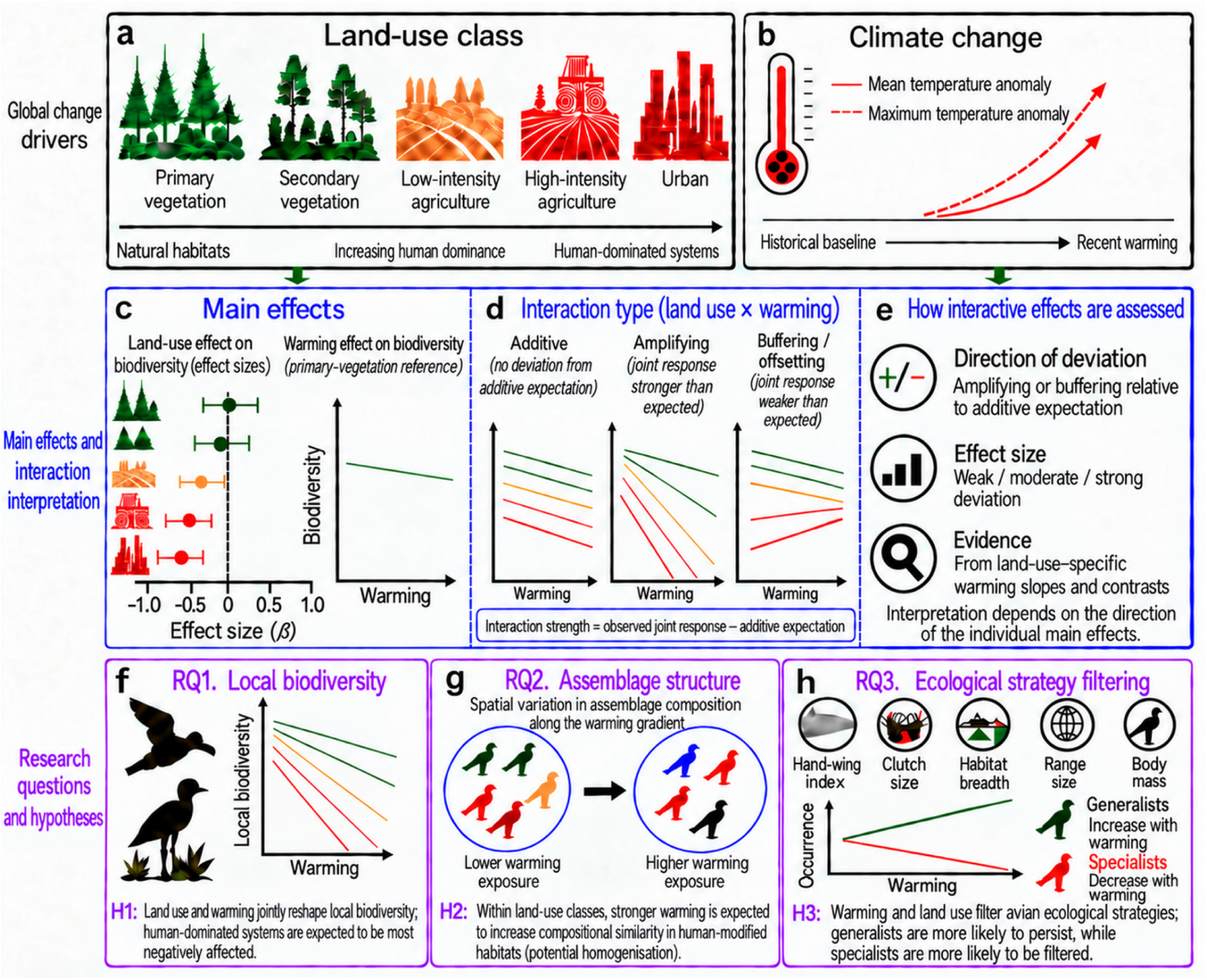
Conceptual framework for testing how land-use class modifies avian biodiversity responses to climate change. The framework links two global-change drivers, land-use change and climate change, to three dimensions of bird diversity change. a, Five categorical land-use classes: primary vegetation, secondary vegetation, low-intensity agriculture, high-intensity agriculture and urban land. b, Climate change is represented by standardised mean- and maximum-temperature anomalies, defined as recent temperature departures from the 1901–1930 local baseline expressed in units of baseline temperature variability. c–e, Effects are interpreted from a land-use-by-warming interaction model. Land-use coefficients represent contrasts with primary vegetation at a temperature anomaly of zero, the warming coefficient represents the warming slope in primary vegetation, and interaction coefficients quantify deviations from that slope in modified land-use classes. On the response scale, interaction strength is defined as the observed joint response minus the additive expectation; responses are interpreted as additive, amplifying or buffering according to the direction and magnitude of this departure and its statistical uncertainty. f–h, The framework addresses three research questions (RQ1–RQ3) and associated hypotheses (H1–H3). f, RQ1 tests whether warming associations with local total abundance and species richness differ among land-use classes, with more negative responses expected in human-modified land uses than in primary vegetation. g, RQ2 tests whether greater warming exposure is associated with changes in pairwise compositional dissimilarity among sites within the same land-use class. Warming-associated homogenisation is inferred where total dissimilarity declines with warming exposure. h, RQ3 tests whether species-level occurrence responses differ among groups defined by shared ecological characteristics. Groups characterised by broader thermal niches, wider geographic ranges and greater dispersal or reproductive capacity are expected to be less negatively affected, whereas range-restricted, less mobile and more specialised groups are expected to be more strongly filtered.

We addressed three questions: First, do the effects of climate change on bird abundance and richness differ among land-use classes? Because land conversion and intensification reduce habitat resources and microclimatic buffering against heat^14,19,20^, we hypothesised that human-modified habitats would show lower bird diversity than primary vegetation, with the strongest warming-associated declines in high-intensity agriculture and urban land. Second, is greater warming exposure associated with higher compositional similarity within land-use classes, and is this pattern driven mainly by species replacement or nestedness? Because land-use and climate change may favour overlapping sets of widespread, disturbance-tolerant species, we expected greater warming exposure to be associated with higher compositional similarity, particularly in modified land uses^9,27,48^, while leaving open whether homogenisation would arise mainly from species replacement or from nested species loss^35,49^. Third, do species-level occurrence responses to land use and warming differ among groups of species sharing similar ecological characteristics? Given that broad niches and high mobility help species persist under environmental change^36–38^, we expected groups characterised by broader habitat and thermal niches, wider geographic ranges and greater dispersal or reproductive capacity to be less negatively affected, or potentially favoured, in some warmer human-modified landscapes, whereas range-restricted, less mobile and more specialised groups would be more strongly filtered under the combined pressures.

We measured warming exposure as standardised temperature anomalies: the difference between mean or warm-season maximum temperature in the five years before sampling and in 1901–1930, divided by the baseline standard deviation^26^. Using mixed-effects models, we related local abundance and richness, compositional dissimilarity between sites of the same study and land-use class, and species occurrence within ecological groups to land use, warming exposure and their interactions, assessing interactions as departures from additive expectations (Methods).

## Results

### Land-use conversion decoupled bird abundance from richness

We first compared land-use classes with mixed-effects models of total abundance (5,817 sites in 121 studies) and species richness (6,457 sites in 138 studies) that accounted for the nested sampling design. Bird total abundance and species richness in human-modified landscapes were generally lower than in primary vegetation, although urban abundance contrasted sharply with this pattern (likelihood-ratio tests: both P < 0.001; Fig. 2b,c). Total abundance was 14.0% lower in secondary vegetation, 12.0% lower in low-intensity agriculture and 16.5% lower in high-intensity agriculture than in primary vegetation, whereas in urban environments it was 32.3% higher, albeit with a wide confidence interval that probably reflects the smaller number of urban sites (Fig. 2b). Species richness was lower in all human-modified habitats, by 9.9% in secondary vegetation, 17.6% in low-intensity agriculture, 18.8% in high-intensity agriculture and 14.8% in urban sites (Fig. 2c).

**Fig. 2.**
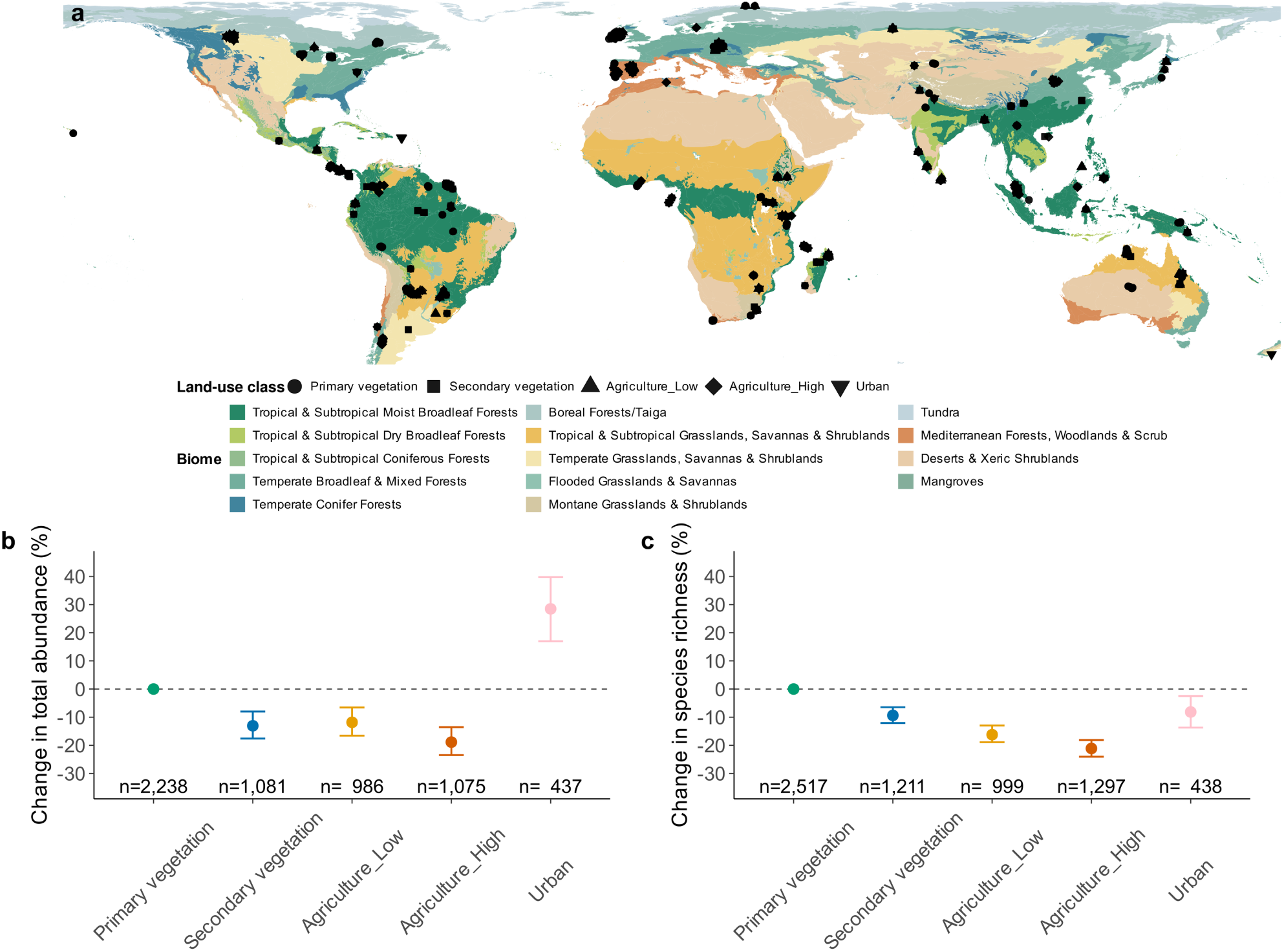
Global distribution of bird sampling sites and local biodiversity responses to land use. a, Geographic distribution of PREDICTS bird-sampling sites used in this study, overlaid on WWF terrestrial biomes^86^. Biomes are distinguished by colour, and point shapes indicate land-use class: primary vegetation, secondary vegetation, low-intensity agriculture, high-intensity agriculture and urban land. b,c, Model-estimated change in local total bird abundance (b) and species richness (c) for each human-modified land-use class relative to primary vegetation (reference). Points show fixed-effect contrasts and error bars indicate 95% confidence intervals; sample sizes (n) below each land-use class are the number of sampling sites contributing to the estimate. Estimates whose intervals exclude zero differ significantly from primary vegetation at the 5% level.

### Warming responses varied among land-use classes

In models including the standardised mean-temperature anomaly and its interaction with land use, warming associations with local bird diversity differed among land-use classes (likelihood-ratio tests: total abundance, χ²₄= 20.23, P = 4.5 × 10⁻⁴; species richness, χ²₄= 60.22, P = 2.6 × 10⁻¹²; Fig. 3a,b; Supplementary Table S1). In primary vegetation, warming associations were weak and not statistically supported for either total abundance (β = 0.049 ± 0.056, z = 0.87, P = 0.39) or species richness (β = 0.074 ± 0.050, z = 1.49, P = 0.14). For species richness, the warming slope was more positive in low-intensity agriculture than in primary vegetation (slope difference β = 0.068 ± 0.017, z = 4.03, P < 0.001) but more negative in high-intensity agriculture (β = −0.063 ± 0.016, z = −3.99, P < 0.001) and urban land (β = −0.160 ± 0.063, z = −2.53, P = 0.011); secondary vegetation showed no detectable difference (β = −0.008 ± 0.021, z = −0.37, P = 0.71; Fig. 3b). For total abundance, only low-intensity agriculture had a significantly more positive warming slope than primary vegetation (β = 0.128 ± 0.035, z = 3.69, P < 0.001; Fig. 3a). Slope differences for secondary vegetation and urban land were also positive but not statistically supported (β = 0.061 ± 0.047, z = 1.30, P = 0.19; β = 0.092 ± 0.114, z = 0.81, P = 0.42), and that for high-intensity agriculture was close to zero (β = −0.007 ± 0.035, z = −0.20, P = 0.84).

**Fig. 3.**
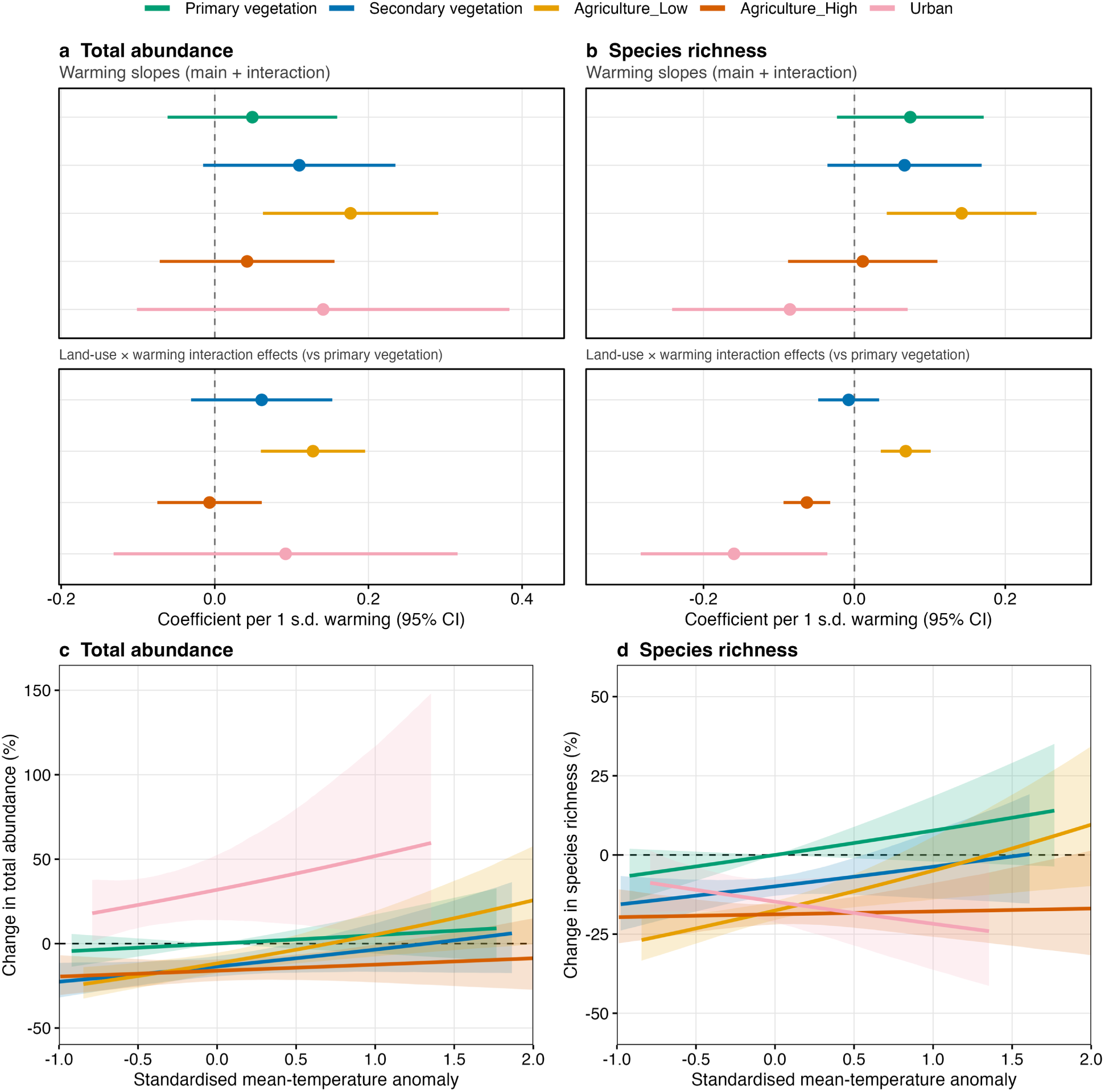
Land-use-specific associations between climate warming and local bird diversity. **a,b,** Warming slopes for total abundance (a) and species richness (b), expressed as the change in log abundance or log expected richness per one-unit increase in the standardised mean-temperature anomaly. Upper sections show the slope in each land-use class; lower sections show the difference between each modified class and primary vegetation (the land-use-by-warming interaction), with positive values indicating a more positive association with warming. Points are estimates and horizontal lines are 95% confidence intervals; vertical dashed lines indicate zero. **c,d,** Model-predicted total abundance (**c**) and species richness (**d**) expressed as percentage differences from primary vegetation at a standardised mean-temperature anomaly of 0. Lines show fixed-effect predictions, and ribbons show 95% confidence intervals derived by parametric simulation of fixed-effect coefficient uncertainty. Horizontal dashed lines indicate no difference from the reference. The total-abundance and species-richness models included 5,817 and 6,457 site-level observations, respectively.

At a standardised temperature anomaly of 1, equivalent to one local baseline standard deviation of warming, predicted total abundance relative to primary vegetation at an anomaly of 0 was 5.2% higher in primary vegetation, 3.7% lower in secondary vegetation, 5.4% higher in low-intensity agriculture, 12.8% lower in high-intensity agriculture and 52.2% higher in urban land (Fig. 3c,d). Predicted species richness was 7.8% higher in primary vegetation but remained lower in all modified classes, by 3.7% in secondary vegetation, 5.0% in low-intensity agriculture, 17.9% in high-intensity agriculture and 21.8% in urban land. Decomposing these joint responses against the additive expectation (Methods) showed that warming counteracted land-use-associated losses in low-intensity agriculture, an antagonistic departure for both richness (5.6%) and abundance (12.3%). By contrast, warming compounded richness losses in high-intensity agriculture (−7.5%) and urban land (−15.6%), a synergistic response, whereas the remaining departures had intervals spanning zero.

Models using the standardised maximum-temperature anomaly (based on warm-season daily maxima) revealed slightly different interaction effects between climate and land-use change (Extended Data Fig. 2). For richness, the positive interaction for low-intensity agriculture was retained (β = 0.074 ± 0.015, P < 0.001), whereas the negative interaction for high-intensity agriculture was weaker and not statistically supported (β = −0.029 ± 0.017, P = 0.094). For total abundance, positive interactions were supported for low-intensity agriculture (β = 0.104 ± 0.032, P = 0.0013) and urban land (β = 0.408 ± 0.127, P = 0.0013).

### Warming-associated changes in assemblage composition

Ordered-beta mixed models related dissimilarity between sites of the same study and land-use class (144,728 site pairs) to their mean warming exposure, accounting for geographic distance, climatic separation and sites shared among pairs. Greater warming exposure was generally associated with lower pairwise dissimilarity within land-use classes (Fig. 4), implying biotic homogenisation through reduced compositional distinctiveness. This association was largely consistent across land-use classes, with little evidence that its slope differed among them, although individual coefficients suggested a steeper increase in replacement-related dissimilarity in secondary vegetation (P = 0.013 –0.034; Extended Data Fig. 3d). Decomposition showed that, within land-use classes, loss-related components (nestedness-resultant and abundance-gradient dissimilarity) tended to decline with increasing warming exposure across non-urban classes. Replacement-related components (turnover and balanced variation in abundance) instead remained generally stable or increased, especially in secondary vegetation. Warming was associated with smaller loss-related differences between assemblages rather than with less species replacement. In primary vegetation, nestedness-resultant dissimilarity, which arises when the poorer site of a pair holds a subset of the species at the richer site, declined with warming (Jaccard β = −0.343, 95% CI −0.601 to −0.085; Sørensen β = −0.321, −0.564 to −0.077), whereas turnover did not, and total dissimilarity showed a weaker decline whose interval overlapped zero (Bray–Curtis β = −0.158, −0.338 to 0.022; Supplementary Table S2). In urban land, the nestedness-resultant slope was positive but highly uncertain. These patterns suggest that convergence arose from declining loss-related differences rather than from species replacement.

**Fig. 4.**
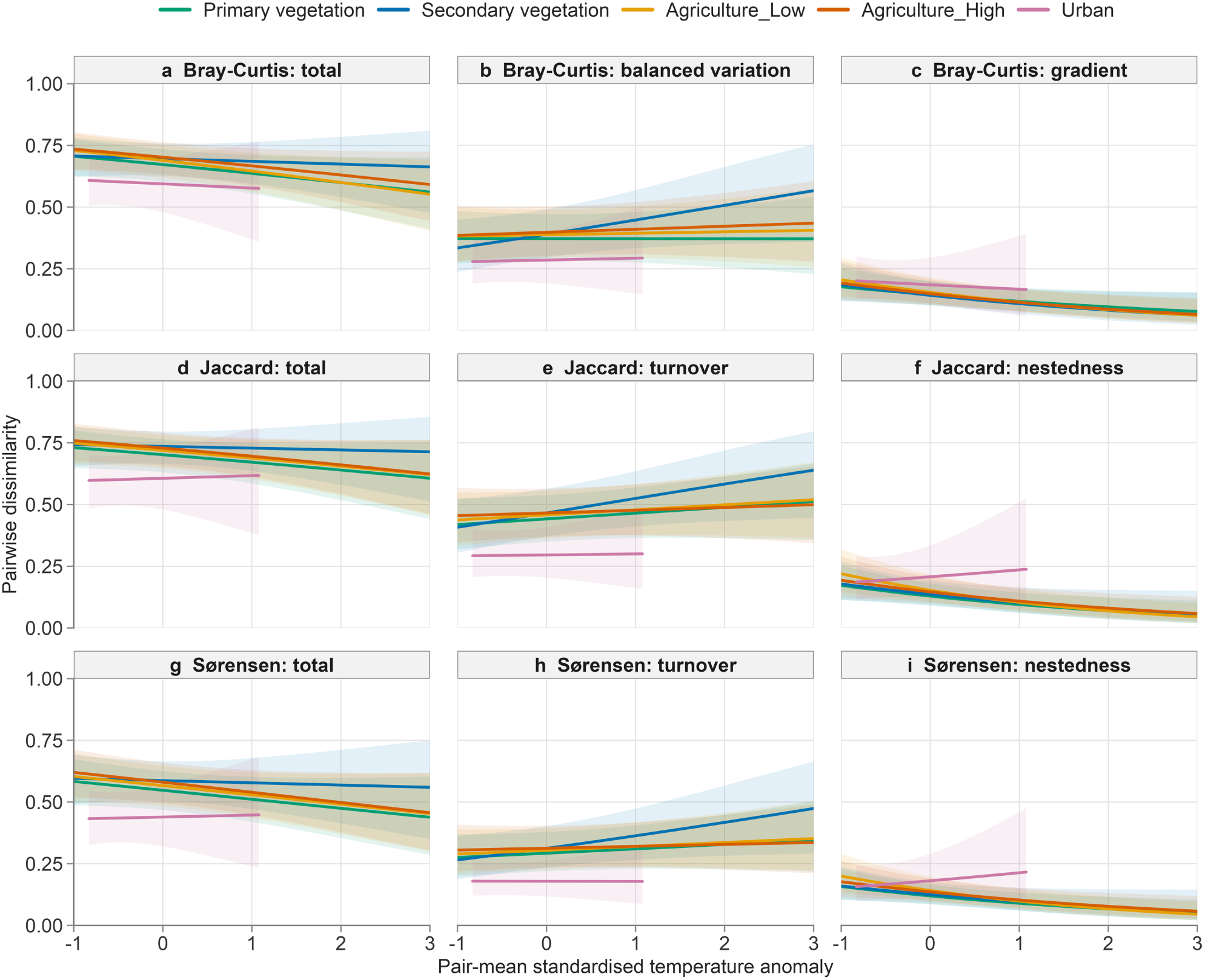
Warming-associated variation in assemblage dissimilarity within land-use classes. Fixed-effect predictions from full-data ordered-beta mixed-effects models showing Bray–Curtis, Jaccard and Sørensen total dissimilarity and their decomposition components along the pair-mean standardised temperature-anomaly gradient. Colours denote land-use class; lines show predicted dissimilarity on the response scale, and shaded bands indicate 95% confidence intervals. For each land-use class, predictions are shown only over the central 95% of its observed pair-mean anomaly distribution. Panels show Bray–Curtis total dissimilarity (a), balanced variation in abundance (b) and abundance-gradient dissimilarity (c); Jaccard total dissimilarity (d), turnover (e) and nestedness-resultant dissimilarity (f); and Sørensen total dissimilarity (g), turnover (h) and nestedness-resultant dissimilarity (i). All models were fitted to 144,728 within-study, within-land-use site pairs from 88 studies and 4,809 sites.

These results were robust to non-independence among site pairs and to the influence of individual studies. In 1,000 subsamples in which no site appeared in more than one pair, median warming slopes had the same sign as the full-data estimates for 38 of 45 metric-by-land-use combinations, and the seven exceptions had intervals including zero in both analyses (Supplementary Fig. S1; Supplementary Table S3a). Leave-one-study-out refits showed no disproportionate influence of any single study, although uncertainty was greatest for urban land (Supplementary Fig. S2; Supplementary Table S3c).

### Ecological winners and losers were land-use dependent

Clustering of eight ecological characteristics in principal-component space identified three stable ecological groups (Methods). Among k-means, partitioning-around-medoids and Ward solutions for K = 2–5, the k-means K = 3 solution best satisfied our pre-specified stability, balance and separation criteria (Extended Data Fig. 4). EG1 comprised larger-bodied, slow-lived species with longer generation lengths (n = 612); EG2 comprised species with wider habitat and realised thermal niches and larger clutch sizes (n = 954); and EG3 comprised species with smaller geographic ranges, lower hand-wing indices and narrower diets (n = 1,380).

The three-way interaction among land use, warming exposure and ecological group was supported in the binomial occurrence model of 432,560 species–site records (likelihood-ratio test: χ²₈= 165.83, P < 0.001). We visualised it as group- and land-use-specific occurrence trajectories along the standardised mean-temperature anomaly (Fig. 5), and decomposed the predictions into the land-use effect at anomaly 0 (Fig. 6a), the warming effect from anomaly 0 to 1 within each land-use class and ecological group (Fig. 6b) and the departure from the additive expectation (Fig. 6c), which we used to classify each contrast (Fig. 6d).

**Fig. 5.**
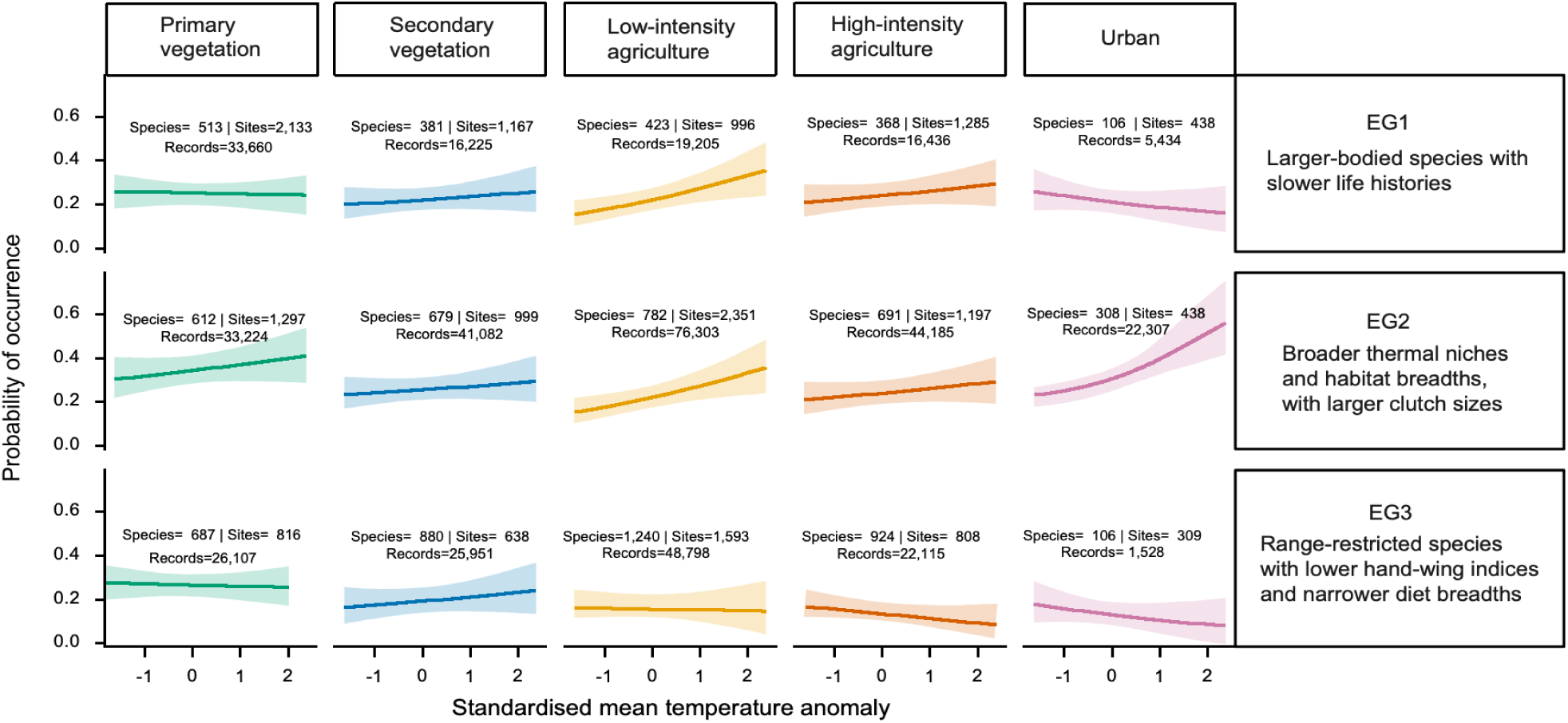
Land-use-specific associations between historical warming and species occurrence across avian ecological groups. Model-predicted occurrence probabilities along the standardised mean-temperature-anomaly gradient for EG1 (top row), EG2 (middle row) and EG3 (bottom row) across primary vegetation, secondary vegetation, low-intensity agriculture, high-intensity agriculture and urban land. EG1 comprised predominantly larger-bodied species with slower life histories; EG2 comprised species with broader realised thermal niches and habitat breadths and larger clutch sizes; and EG3 comprised range-restricted species with lower hand-wing indices and narrower diet breadths. Lines are median predicted values from the binomial generalised linear mixed model fitted to 432,560 species–site records from 6,357 sites, 137 studies and 2,946 species; shaded areas are 95% confidence intervals. Predictions are restricted to the central 95% of standardised temperature anomaly values (2.5th to 97.5th percentiles) observed for each ecological-group-by-land-use combination. Numbers within panels indicate the numbers of species, sites and species–site records contributing to each combination. The overall land-use-by-warming-by-ecological-group interaction was statistically supported (likelihood-ratio test, χ²₈= 165.83, P < 0.001).

**Fig. 6.**
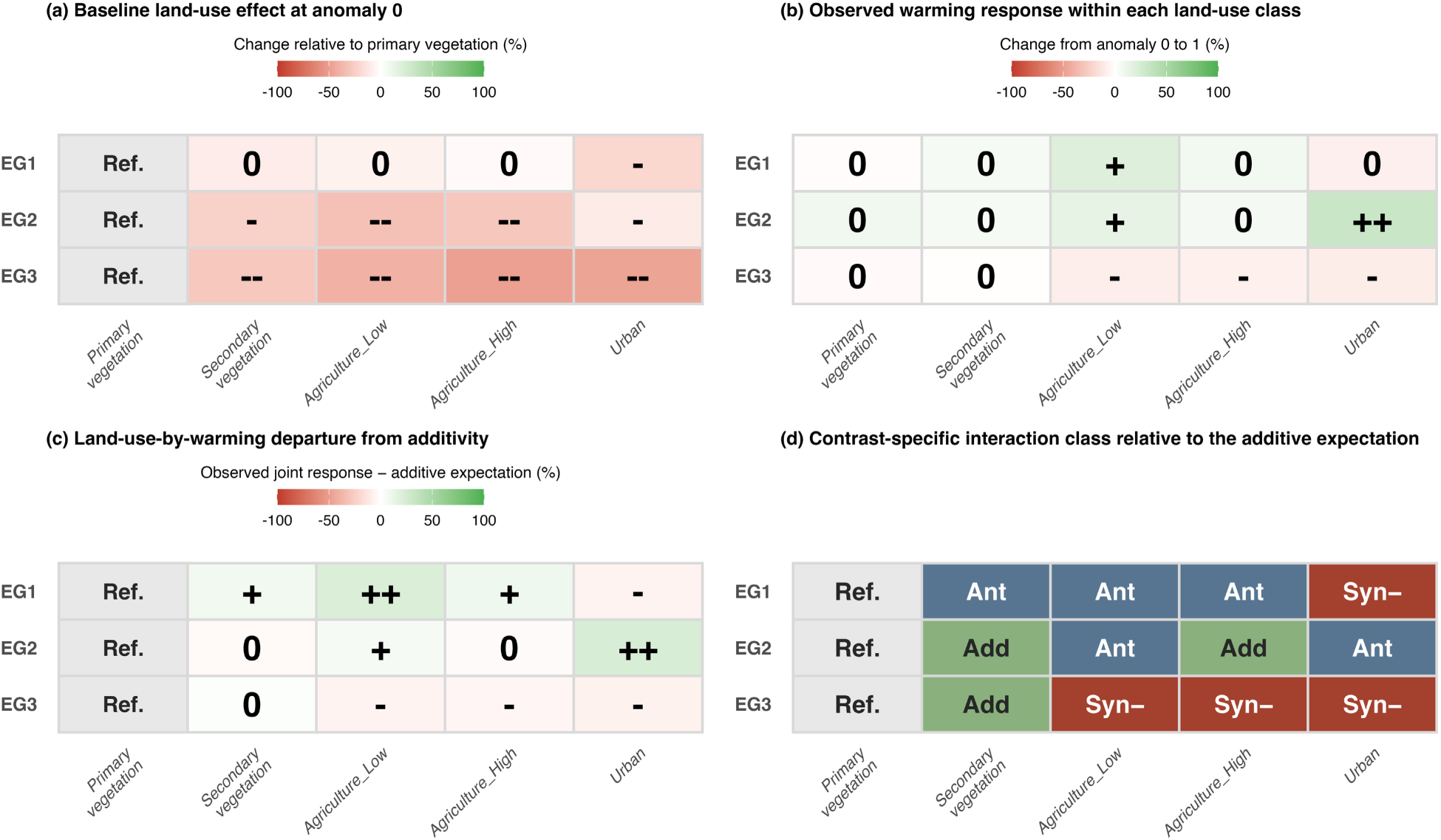
Response-scale decomposition and contrast-specific classification of land-use-by-warming interactions in bird occurrence across ecological groups. All contrasts are percentage differences in predicted occurrence probability relative to the corresponding ecological group in primary vegetation at a standardised mean-temperature anomaly of 0. a, Baseline land-use effects at anomaly 0, the difference between each modified land-use class and primary vegetation. b, Warming responses within each land-use class, the change in predicted occurrence probability from a climate anomaly of 0 to 1. c, Group-specific departures from additivity. The additive expectation is the sum of the baseline land-use effect and the warming effect in primary vegetation; the departure is the observed joint response minus this expectation, equivalently the difference between the warming response in each modified class and that in primary vegetation. Positive values indicate a more positive warming response than in primary vegetation. d, Contrast-specific interaction classification. Responses were classified as synergistic when the observed joint response extended further in the same direction than expected, and as antagonistic when the departure opposed the additive expectation. Departures below 5% were treated as additive, and primary vegetation is the reference and was not assigned a class. Across a–c, plus and minus signs indicate response direction, 0 denotes a change below the lowest threshold, and repeated symbols represent progressively larger magnitudes using graphical thresholds of 5%, 15% and 30%. These thresholds aid visual interpretation only; classes were assigned from the unrounded predicted contrasts and do not indicate statistical significance. The overall land-use-by-warming-by-ecological-group interaction was supported by a likelihood-ratio test (χ²₈ = 165.83, P < 0.001).

Baseline land-use effects at an anomaly of 0 were most pronounced for the range-restricted species of EG3, whose predicted occurrence was 28% lower in secondary vegetation, 40% lower in low-intensity agriculture, 49% lower in high-intensity agriculture and 47% lower in urban land than in primary vegetation (Fig. 6a). Predicted occurrence of the broad-niched species of EG2 was 24–32% lower in secondary vegetation and the two agricultural classes but only 10% lower in urban land, and the larger-bodied, slower-lived species of EG1 were least affected, with reductions of 3–10% across secondary vegetation and the agricultural classes and of 21% in urban land.

Warming responses between anomalies of 0 and 1 differed both among groups and among land-use classes (Fig. 5; Fig. 6b). Predicted occurrence of EG2 increased in every class, most steeply in urban land (32%) and low-intensity agriculture (14%) and by 6–9% elsewhere. EG1 increased in secondary vegetation (6%), low-intensity agriculture (20%) and high-intensity agriculture (8%), was flat in primary vegetation and declined by 8% in urban land. EG3 was flat or declining throughout, with reductions of 1–9% across classes.

Departures from the null additive expectation followed the same ordering (Fig. 6c). The null additive expectation is the occurrence expected if warming had the same effect in a modified land use as in primary vegetation, so a departure measures how far land use altered the warming response. Because occurrence was lower in every modified class than in primary vegetation for all three groups, a positive departure means that warming reduced, and in some cases reversed, this land-use deficit (an antagonistic interaction), whereas a negative departure means that warming deepened it (a synergistic interaction). Departures were positive for EG1 in secondary vegetation (+8%), low-intensity agriculture (+21%) and high-intensity agriculture (+10%) and for EG2 in low-intensity agriculture (+6%) and urban land (+23%), and negative for EG1 in urban land (−6%) and for EG3 in low-intensity agriculture (−6%), high-intensity agriculture (−5%) and urban land (−7%).

Classified against the additive expectation (Fig. 6d), five of the twelve modified-land-use contrasts were antagonistic, with warming buffering the negative land-use effect (EG1 in secondary vegetation, low-intensity agriculture and high-intensity agriculture; EG2 in low-intensity agriculture and urban land). Four were synergistic losses, with warming reinforcing the negative land-use effect (EG3 in low-intensity agriculture, high-intensity agriculture and urban land; EG1 in urban land). The remaining three contrasts (EG2 in secondary vegetation and high-intensity agriculture; EG3 in secondary vegetation) were approximately additive, so most departures from additivity were modest.

### Robustness of the land-use-by-warming interactions

The land-use-by-warming interactions for total abundance and species richness were robust to study influence, climate product and response distribution. They remained supported in all 518 leave-one-study-out refits (abundance, P ≤ 0.042; richness, P ≤ 1.63 × 10⁻⁴; Extended Data Fig. 5a; Supplementary Fig. S3; Supplementary Table S4a) and in harmonised comparisons under CRU TS, TerraClimate, ERA5-Land and CHELSA (abundance χ²₄ = 12.35–35.98, P ≤ 0.015, n = 5,750; richness χ²₄= 41.95–85.08, P ≤ 1.71 × 10⁻⁸, n = 6,386; Extended Data Fig. 5b; Supplementary Table S5a). Across these products, the negative interaction between urban land and warming for richness was significant in every case, whereas the other interaction terms, including that for low-intensity agriculture, varied in magnitude or significance (Extended Data Fig. 6; Supplementary Fig. S4; Supplementary Table S5b). COM–Poisson and generalised-Poisson specifications retained the focal interaction, and the pre-specified negative-binomial model was better supported than both (ΔAIC ≥ 202) while giving the most conservative test (χ²₄= 60.22 versus 65.22 and 72.10; Extended Data Fig. 5c; Supplementary Fig. S5; Supplementary Table S6). Within-study permutation tests found no residual spatial autocorrelation up to 500 km (Moran’s I = −0.0140 to −0.0004; 999 permutations; Benjamini–Hochberg-adjusted P ≥ 0.416; Extended Data Fig. 5d; Supplementary Table S7).

## Discussion

Our global synthesis shows that climate warming modifies how land-use change affects multiple dimensions of avian biodiversity. Land-use-by-warming interactions were supported for local abundance and richness, although individual contrasts varied in direction and strength. Greater warming exposure was also generally associated with lower pairwise compositional dissimilarity within land-use classes, consistent with warming-associated homogenisation. The most consistent cross-dimensional signal was ecological filtering: species with broader realised thermal and habitat niches and larger clutch sizes responded more positively to warming in modified environments, whereas range-restricted species with lower hand-wing indices and narrower diets were most strongly filtered in high-intensity agriculture and urban land.

Our first hypothesis was supported for species richness but not for abundance. Land use did not simply modify baseline biodiversity but mediated how assemblages responded to warming, so an equivalent degree of warming translated into contrasting outcomes across land-use classes. For richness, warming partly offset land-use-associated losses in low-intensity agriculture but compounded them in high-intensity agriculture and urban land. Land-use change shapes species’ exposure, sensitivity and adaptive capacity by altering habitat structure, microclimate, resources, disturbance and dispersal opportunities^14,16^. Low-intensity farmland may retain the shade, moisture and resources that buffer warming, whereas the simplified structure of high-intensity agriculture and urban land can amplify heat exposure^19,20^. Elevated urban abundance may instead reflect reliable anthropogenic food and water supplies favouring a few tolerant species. Joint effects span a continuum from additive to synergistic or antagonistic depending on taxon, scale and context: Suggitt et al.^50^ found additive effects dominated long-term distributional change across British taxa, whereas stronger non-additive responses have been reported for bird assemblages elsewhere^21,26,29^. Such interactions are thus context dependent rather than universal.

The ecological meaning of local diversity responses depends on which species are gained or lost. Apparent weak or positive changes in abundance or richness may reflect gains by disturbance-tolerant species, transient colonisation or shifts in dominance, while masking deeper reorganisation of species identity and function^9,33,34^. Conversely, negative abundance or richness responses in human-modified habitats point to combinations of warming and land-use intensity under which climatic stress and habitat simplification may reinforce each other^16,26,31^, potentially reducing bird-mediated functions such as pest control, seed dispersal and pollination^43,51^. Distinguishing additive, synergistic and antagonistic effects is therefore essential for accurate forecasts and for identifying where warming most amplifies biodiversity loss^21,30^.

Our results reinforce a broader message that local abundance or richness alone cannot fully diagnose ecological reassembly or conservation value. Conservation assessments based on richness alone may therefore overlook the loss or replacement of distinctive, threatened or functionally important species^8,33,52^. In birds, land-use change can replace narrow-ranged or habitat-specialist taxa with widespread, disturbance-tolerant winners without equivalent declines in local richness^9,48^, and such replacement can erode functional stability and ecosystem resilience^44,53^.

Our second hypothesis was only partly supported: greater warming exposure was associated with lower pairwise dissimilarity within land-use classes, but this association did not differ detectably among classes, indicating approximately additive effects on assemblage composition. The broadly consistent direction of these responses across several dissimilarity metrics supports emerging compositional convergence, although statistical support varied among components and land-use classes and was weaker where empirical coverage was limited, particularly in urban land. Importantly, the focal predictor represented the warming context of each site pair rather than the climatic difference between sites. Accordingly, the analysis tests whether assemblages in warmer contexts are more or less similar after accounting for geographic distance and climatic separation, indicating that compositional convergence is context dependent rather than universal. Recent syntheses similarly caution that biodiversity change involves multiple, scale-dependent dimensions rather than a single trajectory^54^. Rather than indicating universal homogenisation, our results suggest that greater warming exposure is associated with reduced loss-related compositional differences.

Our third hypothesis was supported, and ecological groups defined by shared ecological characteristics provided the strongest and most interpretable pattern. The positive responses of broad-niched, higher-fecundity generalists under modified and warmer conditions are consistent with human-modified landscapes favouring species able to exploit many habitats, whereas the reduced occurrence of range-restricted and diet-specialist strategies in high-intensity agriculture and urban land indicates filtering of species with limited breadth, dispersal capacity or specialised requirements. Warming counteracted land-use-associated occurrence losses for larger-bodied and broad-niched groups (EG1 and EG2) in low-intensity agriculture but compounded losses for range-restricted, diet-specialised species (EG3) across agricultural and urban land (Fig. 6). This aligns with comparative evidence that narrow ranges and habitat specialisation confer higher sensitivity to both drivers^38^. Consistent losses of sensitive specialists across sites, while generalists persist, may also explain the warming-associated decline in loss-related dissimilarity, as assemblages converge on similar subsets of widespread species. Trait composition, in turn, influences how communities respond to disturbance and which functions they support^40^, while intensive land use can reorganise avian functional diversity over long periods^27,55^. Our results extend this literature by showing that the functional signature of land-use modification is not fixed but is expressed in interaction with warming, which changes not only how many species occur locally but which functional strategies persist.

These findings have direct implications for biodiversity scenario forecasting and conservation. Models treating climate and land-use change as independent risk underestimating responses where joint effects amplify losses and overestimating them where interactions buffer losses. Here, an additive model would underestimate richness losses in high-intensity agriculture and urban land and occurrence losses of range-restricted species (EG3), but overestimate richness and abundance losses in low-intensity agriculture. Analyses based only on aggregate biodiversity metrics may also obscure contrasting responses among ecological groups that are not apparent from changes in richness or abundance alone. Previous work has shown that the two drivers jointly shape biodiversity outcomes^9,12,31^; our results indicate that the direction and magnitude of their interaction are themselves informative and should be reported alongside their separate effects. For conservation, climate-smart planning must protect more than species numbers. Maintaining primary and semi-natural vegetation, reducing modification intensity, retaining habitat heterogeneity and improving landscape permeability may buffer warming-driven community reassembly^5,28^, and integrated strategies combining conservation, restoration and land-system transformation are more likely to bend trajectories than single-driver interventions^56,57^. Safeguarding community distinctiveness and functional diversity is especially important for range-restricted and specialist species least likely to persist under continued warming.

Several limitations bound these inferences. First, PREDICTS is a space-for-time database rather than a long-term monitoring network, so our models evaluate spatial associations along contemporary warming gradients and cannot resolve lagged dynamics, extinction debts or colonisation credits documented in temporal studies^58,59^. Second, the analyses are observational and cannot fully separate direct warming effects from correlated differences in land-use history, management, habitat quality, or biotic interactions^60,61^. Third, temperature anomalies were derived from gridded macroclimatic data and do not capture microclimatic variation across canopy structure and topography^14,20^. Fourth, other climatic and anthropogenic pressures, including extreme climatic events, overexploitation and invasive species, may also influence assemblage change and covary with the focal drivers^62,63^. Finally, some urban land-use-by-ecological-group combinations had fewer observations and a narrower temperature-anomaly range, so those estimates should be interpreted cautiously.

Future studies should combine long-term monitoring with fine-resolution land-use and microclimate data, designs that account for imperfect detection, and mechanistic models linking environmental change to community dynamics^64,65^. Extending the framework to functional diversity and other taxa would test whether the winners and losers identified here are general signatures of assemblage reorganisation, and whether biotic homogenisation strengthens as pressures accumulate. Coupling these advances with land-use and climate scenarios is essential for projecting where biodiversity change will involve local biodiversity loss, compositional convergence, functional erosion or their combination, especially where rapid land-use conversion coincides with accelerating warming.

## Methods

### Overview

The PREDICTS database compiles biodiversity records from globally distributed sites sampled using consistent methods within individual studies, across different land-use classes and use-intensity classes^42,45,46^. For this work, we focused on bird records from the PREDICTS database with site-level estimates of recent warming, quantified as standardised temperature anomalies derived from CRU TS v4.09^47^, and species-level ecological characteristics compiled from AVONET and complementary published datasets^40^. We tested how land use and climate change interact to shape bird diversity across three complementary dimensions of ecological organisation (Fig. 1): (i) site-level total abundance and species richness (local biodiversity); (ii) pairwise dissimilarity in species composition among local assemblages and its metric-specific turnover, nestedness-resultant and abundance-gradient components (assemblage composition); and (iii) species-level occurrence within groups defined by shared ecological characteristics (ecological filtering).

### Biodiversity data and preprocessing

Biodiversity data were extracted from the PREDICTS (Projecting Responses of Ecological Diversity in Changing Terrestrial Systems) database (version 1.1, https://data.nhm.ac.uk/dataset/the-2016-release-of-the-predicts-database), a global compilation of spatial comparisons of local biodiversity across land-use types and intensities^42,45^. PREDICTS records are drawn from published sources or from unpublished datasets with published sampling protocols. Species-level abundance measurements comprise approximately 85% of records, occurrence records account for almost all of the remainder, and richness-only records constitute a very small fraction. We supplemented these with the November 2022 release of data added to the PREDICTS database, which extends the 2016 release with 115 additional data sources and is available from the Natural History Museum data portal^46^. The database has a hierarchical structure: each data source comprises one or more studies conducted using a consistent sampling methodology; within each study, sites are grouped into one or more spatial blocks that capture spatial clustering in the sampling design; and each block contains one or more georeferenced sites^42,45^.

We extracted all bird records from sites with known predominant land-use type and land-use intensity, excluding records assigned to “Cannot decide” land-use classes. Species identities were standardised to the BirdLife International/HBW taxonomy used by AVONET before trait matching^40,66^. From these records we derived three response datasets. Site-level species richness was the number of species recorded at each site, whereas total abundance was the sum of abundance-type measurements across species, including individual counts, point-count totals, transect counts and density estimates. Pairwise compositional dissimilarity was calculated among sites within the same study and land-use class, and species-by-site occurrence matrices were constructed relative to each study’s observed species pool. Bray–Curtis (abundance-based) and Jaccard and Sørensen (incidence-based) dissimilarities were calculated for each pair and partitioned into replacement- and loss-related components. Because sites recurred across pairs, models included random intercepts for both sites of each pair, and refits to subsamples in which each site entered only one pair tested sensitivity to this pseudo-replication (see Assemblage composition within land-use classes).

Where sampling effort varied among sites within a study, only raw count-based abundance measures expected to scale with sampling effort were corrected. Recorded sampling effort was scaled relative to the maximum effort within that study, and the corresponding abundance values were divided by relative effort, assuming a linear relationship between effort and observed abundance^3,26^. This correction affected 7.5% of non-zero abundance measurements (49,347 of 656,539 records). Effort-corrected site-level abundance was then rescaled within each study to the interval [0,1] by dividing each value by the maximum abundance observed in that study. This preserved within-study abundance contrasts while reducing differences in measurement scale among studies arising from variation in sampling units, protocols and taxonomic resolution. Rescaled abundance was then log-transformed after adding a small positive constant of 0.01 to accommodate zero values. The final analytical datasets comprised 5,817 site-level observations from 121 studies for total abundance and 6,457 observations from 138 studies for species richness.

### Land-use classification

PREDICTS classifies sites by predominant land-use type and land-use intensity using a standardised global scheme^42,45^. Original land-use categories are primary vegetation, secondary vegetation, plantation forest, cropland, pasture and urban. Primary vegetation comprises native forest or non-forest vegetation not known to have ever been modified, whereas secondary vegetation comprises vegetation recovering after substantial disturbance or clearance, with successional stage determined from vegetation structure. Cropland comprises land planted with herbaceous crops, pasture comprises land used regularly or permanently for livestock grazing, plantation forest comprises previously cleared land planted with crop trees or shrubs, and urban land comprises areas dominated by human settlements or buildings^26,42,45^. Within each land-use class, land-use intensity is classified as minimal, light or intense according to land-use-specific evidence of management and disturbance, including resource extraction, grazing, hunting, agrochemical inputs and mechanisation^42,45^. To obtain ecologically interpretable land-use classes with adequate sample sizes across the warming gradient, we aggregated the original PREDICTS land-use and land-use-intensity categories into five analytical classes: primary vegetation, secondary vegetation, low-intensity agriculture, high-intensity agriculture and urban land (Supplementary Methods, section S2). Primary vegetation was used as the reference class. Secondary vegetation comprised vegetation regenerating after substantial disturbance or clearance, with all successional stages pooled. Cropland and pasture under minimal or light use, together with minimally used plantation forest, were assigned to low-intensity agriculture; intensely used cropland and pasture, and plantation forest under light or intense use, were assigned to high-intensity agriculture^26^. Urban land was retained as a single class irrespective of recorded intensity because individual urban-intensity categories had insufficient sample sizes for robust estimation, but urban environments impose distinct structural, thermal and resource filters (including predominantly managed vegetation) that may produce biodiversity responses different from those in agricultural systems^9,53^.

### Climate-change variables

Recent warming exposure at each site was quantified using monthly mean and maximum temperature estimates from CRU TS v4.09, which provides monthly gridded climate estimates for 1901–2024 at 0.5° × 0.5° spatial resolution^47^. For each PREDICTS site, we extracted monthly temperature values from the corresponding CRU TS grid cell. Following Outhwaite et al.^26^, the historical baseline was defined as 1901–1930, and the recent climate window as the five years preceding the final year of biodiversity sampling at that site. This window captured recent warming most likely to influence observed assemblages while reducing sensitivity to short-term interannual temperature fluctuations^26,58^. The structure of this warming-exposure gradient within and among studies, for every analysis sample and warming metric, is summarised in Supplementary Fig. S6 and Supplementary Table S8.

The standardised mean-temperature anomaly for site *s* was computed as the difference between the recent-window monthly mean temperature and the 1901–1930 baseline mean, divided by the baseline standard deviation:

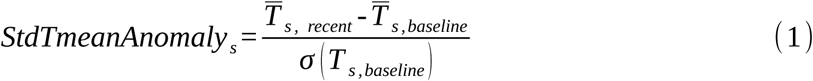

where *T̄ _s, recent_* is the mean monthly temperature during the five years preceding the final year of biodiversity sampling, *T̄ _s,baseline_* is the mean monthly temperature during the 1901–1930 baseline period, and *σ* ( *T _s,baseline_* ) is the standard deviation of monthly temperatures during that baseline period. A value of 1 therefore represents a recent temperature departure equivalent to one local historical standard deviation above the 1901–1930 mean. Because the baseline standard deviation was calculated from monthly temperatures across the full baseline period, this metric incorporates both seasonal and interannual variability and expresses warming relative to the historical climatic variability at each site, placing sites from contrasting climatic regions on a comparable, locally relative scale (Extended Data Fig. 1). An analogous standardised maximum-temperature anomaly was calculated from the monthly mean daily maximum temperatures of the three warmest months in each year, using the same five-year recent window and 1901–1930 baseline^26^; it therefore captures changes in warm-season daytime maxima rather than short-duration heat extremes. The standardised mean-temperature anomaly was the primary warming predictor, and the maximum-temperature anomaly was used in complementary analyses.

For analyses of compositional dissimilarity, each site pair_ij_ was assigned two climate predictors. Warming context was quantified as the mean standardised temperature anomaly of the two sites, whereas climatic separation was quantified as their absolute anomaly difference. All climate analyses were interpreted as space-for-time comparisons^67^. We could not infer temporal change at individual sites; instead, we quantified how biodiversity differed among sites, or among site pairs, when exposed to different levels of recent warming relative to their historical baselines.

### Species characteristics and ecological groups

We compiled eight ecologically informative characteristics expected to mediate species responses to land-use change and climate warming: body mass, geographic range size and clutch size^40^; habitat breadth, the number of major IUCN habitat classes occupied by each species^38^; diet breadth, the number of major dietary categories used^68^; thermal niche breadth, the difference between the upper and lower realised thermal limits across each species’ geographic range^69^; hand-wing index, a morphological proxy for dispersal capacity^70^; and generation length^71^. Together these characteristics summarise morphology, life history and ecological niche, all linked to global-change sensitivity in birds^37,39^.

Missing ecological-characteristic values were imputed using missForest^72^, restricting imputation to species with more than half of the eight focal characteristics observed. Only thermal niche breadth (113 of 2,946 species; 3.84%) and diet breadth (63 species; 2.14%) contained missing values. In total, 176 of 23,568 trait values (0.75%) were imputed across 114 species (3.87%). Imputation accuracy was evaluated using five repeats of ten-fold masked cross-validation, with normalised root-mean-square error (NRMSE) calculated as the root-mean-square prediction error divided by the standard deviation of the observed values. Cross-validated NRMSE was 0.45 for thermal niche breadth and 0.62 for diet breadth, with corresponding observed–predicted Pearson correlations of 0.91 and 0.82. These results indicate good predictive performance for both ecological characteristics. Reconciliation of the trait-imputation and occurrence-model species denominators is given in Supplementary Table S9.

After imputation, ecological characteristics were transformed to reduce skewness before PCA and clustering. Habitat breadth and diet breadth were square-root transformed to stabilise variance in these count variables, whereas the remaining right-skewed characteristics were natural-log transformed. All continuous characteristics were then winsorised at the 1st and 99th percentiles and standardised to zero mean and unit variance to prevent variables measured on larger scales from dominating multivariate distances. We aimed to identify recurrent multivariate ecological strategies, defined as groups of species sharing similar combinations of characteristics, rather than to evaluate each characteristic independently. To reduce collinearity and summarise the principal axes of variation, we projected the standardised trait matrix into principal component space and compared k-means, partitioning around medoids and Ward hierarchical clustering for K = 2–5. Candidate solutions were evaluated using average silhouette width, minimum bootstrap Jaccard stability across 100 nonparametric resamples and minimum cluster proportion^73^. The k-means K = 3 solution was retained as the largest candidate satisfying the pre-specified stability and balance criteria (Extended Data Fig. 4): minimum bootstrap Jaccard stability ≥ 0.55 for every cluster and minimum cluster proportion ≥ 0.20, while retaining an average silhouette width within 90% of the best-performing eligible solution^74,75^. Clustering-solution screening, principal-component loadings, explained variance and ecological-group profiles are reported in Supplementary Table S10.

### Statistical analyses

Mixed-effects models were used to estimate responses of local abundance, richness, within-land-use assemblage dissimilarity and species-level occurrence to land use, climate change and their interactions. Models were fitted by maximum likelihood in glmmTMB v1.1 using the Laplace approximation^76^ in R^77^. Study (SS) and spatial block (SSB) within study were included as random intercepts where appropriate to account for variation in sampling protocol, study scope and spatial arrangement. The occurrence model additionally included site (SSBS) and species random intercepts, whereas pairwise assemblage models included crossed site random intercepts for the two sites in each pair. Fixed effects were land use, standardised mean-temperature anomaly and, for occurrence, ecological strategy group. The focal inferential terms were the land-use-by-warming interaction for assemblage-level analyses and the land-use-by-warming-by-ecological-strategy interaction for occurrence (see below for equations). Sampling uncertainty in response-scale percentage change was quantified by parametric simulation. We drew 1,000 fixed-effect coefficient vectors from a multivariate normal distribution defined by the fitted estimates and their variance–covariance matrix. For each draw, marginal predictions were generated with random effects set to zero, back-transformed to the response scale, and expressed relative to primary vegetation at a standardised temperature anomaly of 0. The 2.5th and 97.5th percentiles provided 95% confidence intervals. Predictions were restricted to the observed anomaly range of each land-use class.

### Local biodiversity models

To address the first question, we modelled site-level total abundance and species richness as a function of land use, standardised mean-temperature anomaly and their interaction. We repeated the analyses using the standardised maximum-temperature anomaly as an alternative measure of warming exposure. Random intercepts for study (SS), spatial block within study (SSB), and site within study and block (SSBS) accounted for the hierarchical PREDICTS sampling design.

Total abundance was modelled as log-transformed within-study rescaled abundance using a Gaussian linear mixed-effects model:

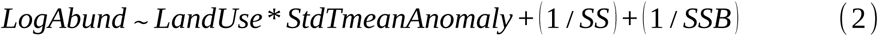

This model followed the PREDICTS framework for analysing abundance responses to land use and climate exposure^3,26^. Land use, standardised mean-temperature anomaly and their interaction were included as fixed effects.

Species richness was modelled as count data using a negative-binomial distribution with a log link. We specified a quadratic mean–variance relationship, Var(Y)=μ+μ^2^/θ, allowing variance to increase faster than the expected richness and thereby accommodating overdispersion. The negative-binomial model was preferred after preliminary Poisson fits showed residual overdispersion using DHARMa diagnostics^78^. The fitted model was:

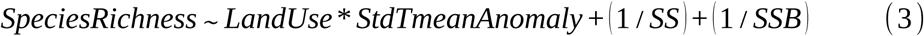

To assess whether associations with standardised temperature anomalies were confounded by long-term background climate, we refitted the local-biodiversity models with site-level climatological mean annual temperature for 1901–1930, derived from CRU TS v4.09^47^, as an additional covariate. The direction and statistical support of the focal land-use-by-warming interactions were qualitatively unchanged. Coefficient comparisons before and after this adjustment are given in Supplementary Fig. S7 and Supplementary Table S11.

### Assemblage composition within land-use classes

To address the second question, we tested whether warming exposure was associated with variation in assemblage composition within land-use classes. Pairwise compositional dissimilarity was calculated only between sites sampled within the same study and assigned to the same land-use class, allowing us to assess whether assemblages exposed to greater warming were more or less compositionally similar within a common land-use context.

Bray–Curtis dissimilarity was used as the primary abundance-based metric, with Jaccard and Sørensen dissimilarities used to assess consistency for incidence-based composition. Bray–Curtis dissimilarity was partitioned into balanced variation in abundance, representing reciprocal changes in species abundances, and abundance-gradient dissimilarity, representing predominantly unidirectional abundance differences between assemblages^49,79^. Jaccard and Sørensen dissimilarities were partitioned into turnover, representing species replacement, and nestedness-resultant dissimilarity, arising when one assemblage is largely a subset of another. We refer to turnover and balanced variation as replacement-related components, and to nestedness-resultant and abundance-gradient dissimilarity as loss-related components. Total dissimilarity and each component were modelled separately using ordered-beta mixed-effects models with a logit link, allowing exact zeros and ones to be retained^80^:

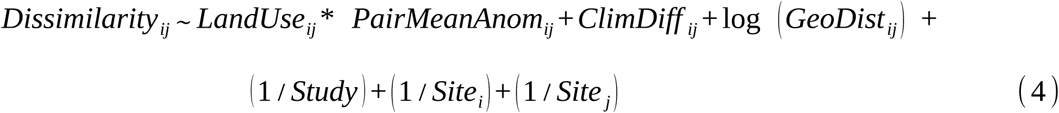

We considered only pairs of sites sampled within the same study and assigned to the same land-use class. Thus, LandUse_ij_ denotes the same land-use class of sites i and j, allowing us to assess warming-associated variation in assemblage dissimilarity within rather than between land-use classes. PairMeanAnom_ij_ is the mean standardised temperature anomaly of the two sites, ClimDiff _ij_ is their absolute difference in temperature anomaly, and log(GeoDist_ij_) accounts for distance decay in assemblage similarity. Random intercepts for study and the two site identities accounted for study-level heterogeneity and the repeated occurrence of individual sites across pairwise comparisons. The coefficient for PairMeanAnom_ij_ represents the warming–dissimilarity association in primary vegetation, whereas its interactions with land use test whether this association differs among the other land-use classes. A negative association between warming exposure and total dissimilarity indicated increasing compositional similarity, or biotic homogenisation. The robustness of this interpretation was assessed from the consistency of effect directions across abundance- and incidence-based metrics and the corresponding 95% confidence intervals. The decomposed components were used to distinguish species replacement from nestedness- or abundance-gradient-related differences among assemblages^49,79^. Of the 147,547 eligible within-study, within-land-use Bray–Curtis site pairs, 144,728 had complete covariate data and entered the fitted models: 52,276 in primary vegetation, 20,253 in secondary vegetation, 17,097 in low-intensity agriculture, 45,735 in high-intensity agriculture and 9,367 in urban land.

Because individual sites contributed to multiple pairwise comparisons, we assessed sensitivity to shared-site dependence using repeated strictly disjoint-pair subsampling, following Newbold et al.^81^. In each iteration, sites were randomly paired without replacement within each study-by-land-use stratum, ensuring that each site contributed to at most one comparison; one site was omitted when a stratum contained an odd number of sites. The nine ordered-beta models were refitted to each subsample with a study-level random intercept, and coefficients were summarised by their medians and 2.5th and 97.5th percentiles across iterations. Additionally, leave-one-study-out refits were used to identify results disproportionately influenced by individual studies.

### Ecological-strategy occurrence models

To address the third question, we tested whether groups of species characterised by different combinations of ecological characteristics differed in their occurrence responses to land use and climate change. Within each study, species detected at a site were coded as 1, whereas species detected elsewhere in the same study but not at the focal site were coded as 0. This study-specific species pool avoided treating species that were never sampled within a study as absences. We modelled occurrence rather than abundance to evaluate how species-level occurrence probabilities varied among ecological groups under different combinations of land use and warming exposure, while improving comparability among studies that used different abundance measures and sampling protocols. Occurrence was modelled with a binomial generalised linear mixed model and a logit link, incorporating the three-way interaction among land use, standardised mean-temperature anomaly and ecological group:

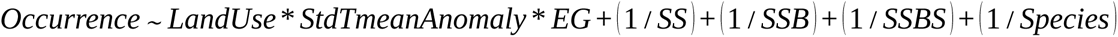

The site-level intercept additionally accounted for multiple species records within sites, while the species-level intercept accounted for repeated observations and baseline differences in occurrence among species. The model included 432,560 species–site records from 6,357 sites, 598 spatial blocks, 137 studies and 2,946 species assigned to ecological groups. Calibration and residual patterns for this model, and its sensitivity to the spatial-block random intercept, are reported in Supplementary Fig. S8 and Supplementary Table S12.

To interpret the three-way interaction, we estimated occurrence probabilities for each ecological group and land-use class at standardised temperature anomalies of 0 and 1. The warming response was defined as the change in predicted occurrence probability between these two anomaly levels. We then compared these responses pairwise among ecological groups within each land-use class. A positive contrast indicated that the focal group increased more, or declined less, with warming than the comparison group; a negative contrast indicated the opposite.

### Quantifying additive, synergistic and antagonistic responses

Each model included the land-use-by-warming interaction required to test whether the two drivers contributed independently. We defined the additive expectation as the response predicted when land use and warming acted independently on the relevant model scale. Statistical support for land-use-by-warming interactions was evaluated using likelihood-ratio tests comparing the full interaction model with the nested main-effects-only model^26^. Where the interaction model was supported, inference was based on fitted model coefficients, land-use-specific warming slopes, model-based contrasts and uncertainty intervals.

For ecological interpretation, we then evaluated joint responses on the response scale using model predictions. Interaction strength was calculated as the departure of the observed joint response from the additive expectation. Joint responses were interpreted as approximately additive when this departure was close to zero, as amplifying when the joint response was stronger than expected under additivity, and as buffering or offsetting when the joint response was weaker than expected. Because the ecological meaning of a positive or negative departure depends on the direction of the individual land-use and warming effects, response-scale classifications were interpreted relative to whether the expected outcome represented biodiversity loss or gain. Departures from additive expectations were classified according to their direction and magnitude. For detrimental responses, joint effects that were more negative than expected were classified as amplified losses, whereas less-negative effects indicated buffering and reversals to positive effects indicated offsetting. For beneficial responses, more-positive-than-expected effects were classified as amplified gains, whereas less-positive effects indicated buffering and reversals to negative effects indicated offsetting. When both individual effects acted in the same direction, amplified losses and gains corresponded to negative and positive synergistic effects, respectively, whereas buffering or offsetting indicated antagonistic effects. These response-scale categories were used for ecological interpretation and visualisation; statistical inference was based on model comparisons, contrasts and their uncertainty intervals.

To interpret the three-way interaction, we predicted occurrence probabilities for each ecological group and land-use class at standardised temperature anomalies of 0 and 1. Within each ecological group, primary vegetation at anomaly 0 served as the reference. The baseline land-use effect was defined as the contrast between each modified land-use class and primary vegetation at anomaly 0, and the warming response within each land-use class as the change in predicted occurrence from anomaly 0 to 1. For each modified land-use class, we then defined a reference-based additive expectation as the sum of its baseline land-use effect and the warming response in primary vegetation. This represents the joint response expected if warming produced the same response-scale change in the modified land-use class as in primary vegetation. The non-additive component was calculated as the observed joint response minus this additive expectation, equivalently the difference between the warming response in the modified land-use class and that in primary vegetation. Thus, the departure from additivity quantifies land-use-specific modification of the warming response relative to primary vegetation, rather than relative to an average warming response across land-use classes. All effects were expressed relative to the group-specific reference probability. For graphical classification, response-scale shifts of <5% were treated as near-zero, and shifts of 5–15%, 15–30% and >30% were labelled with one, two and three symbols, respectively; positive shifts were denoted by “+”, “++” and “+++”, and negative shifts by “−”, “−−” and “−−−”. These thresholds aided visual interpretation only: overall interaction terms were tested with likelihood-ratio tests, and uncertainty in specific model-based contrasts was represented by 95% confidence intervals. We applied the same decomposition to the total-abundance and species-richness models, using fixed-effect predictions at anomalies of 0 and 1 relative to primary vegetation at anomaly 0; 95% confidence intervals were obtained from 10,000 parametric simulations of the fixed-effect coefficients, and departures whose intervals excluded zero were classified as synergistic or antagonistic.

### Model diagnostics and sensitivity analyses

Model diagnostics and sensitivity analyses were tailored to response distributions and model structure. For Gaussian abundance, we inspected residual-versus-fitted plots, Q-Q plots and residual variance structure. For negative-binomial richness and binomial occurrence models, we used DHARMa simulation diagnostics to assess residual uniformity, overdispersion, zero inflation and outliers^78^. To assess sensitivity of the species-richness inference to the assumed count distribution, we refitted the richness model using Conway–Maxwell–Poisson (COM–Poisson) and generalised-Poisson distributions, retaining the same fixed- and random-effect structure, and compared model support and the land-use-by-warming interaction across distributions. Residual spatial autocorrelation was evaluated within studies across three distance bands: 0–50, 50–200 and 200–500 km, using Moran’s I with 999 within-study permutations. Influential studies were screened using leave-one-study-out refits of the local-biodiversity and assemblage-composition models. To assess sensitivity to climate-product choice and spatial resolution, we repeated the focal climate-exposure analyses using site-level standardised temperature anomalies from alternative gridded climate products. In addition to the primary CRU TS v4.09 monthly mean-temperature data at 0.5° resolution, standardised relative to the 1901–1930 baseline^47^, we repeated the analyses using TerraClimate at approximately 1/24° resolution with a 1958–1987 baseline^82^, ERA5-Land at approximately 0.1° resolution with a 1950–1979 baseline^83^, and CHELSA v2.1 at 30-arcsec resolution with a 1981–2000 baseline^84,85^. For each product, recent climate exposure was calculated from the 60 months preceding the end of biodiversity sampling at each site, and anomalies were standardised using the mean and standard deviation of monthly temperatures during the corresponding product-specific baseline. The qualitative direction of the focal land-use-by-warming associations was broadly consistent across products, although statistical support varied, with ERA5-Land showing the largest deviations. Simulation-based residual diagnostics and residual offsets across land-use classes and anomaly bins are reported in Supplementary Figs. S9 and S10 and Supplementary Tables S13 and S14, and product sensitivity for assemblage dissimilarity in Supplementary Fig. S11. Variance components for every fitted model, and a crosswalk from each principal claim to the corresponding supplementary evidence and its interpretation limits, are given in Supplementary Tables S15 and S16.

## Data availability

All data underlying this study are publicly available. Bird assemblage records were obtained from the PREDICTS database: the 2016 release (v1.1) and the November 2022 data release, both available from the Natural History Museum Data Portal (https://data.nhm.ac.uk/dataset/the-2016-release-of-the-predicts-database-v1-1; https://data.nhm.ac.uk/dataset/release-of-data-added-to-the-predicts-database-november-2022). Monthly mean and maximum temperature data were obtained from CRU TS v4.09 (https://crudata.uea.ac.uk/cru/data/hrg/cru_ts_4.09/). Alternative climate products used in sensitivity analyses are available from TerraClimate (https://doi.org/10.1038/sdata.2017.191), ERA5-Land (https://doi.org/10.24381/cds.68d2bb30) and CHELSA v2.1 (https://doi.org/10.16904/envidat.686). Species ecological characteristics were compiled from AVONET (body mass, geographic range size and clutch size; https://doi.org/10.6084/m9.figshare.16586228.v7), Etard and Newbold^38^ (habitat breadth; https://doi.org/10.1111/cobi.14208), BIRDBASE (diet breadth; https://doi.org/10.6084/m9.figshare.27051040.v1), Williams and Newbold^69^ (realised thermal niche breadth; https://doi.org/10.5061/dryad.sj3tx964n), Sheard et al.^70^ (hand-wing index; also available in AVONET) and Bird et al.^71^ (generation length; https://doi.org/10.1111/cobi.13486). Species names were standardised to the HBW–BirdLife Taxonomic Checklist v5.0 (https://datazone.birdlife.org/). Terrestrial biome boundaries shown in Fig. 2a follow Olson et al.^86^. The processed site-level analysis datasets, site-pair dissimilarity data and source data for all figures are archived at Zenodo (https://doi.org/10.5281/zenodo.22516089).

## Code availability

All code used for data processing, model fitting, sensitivity analyses and figures is available at Zenodo (https://doi.org/10.5281/zenodo.22516089).

## Author contributions

C.D., T.N., Z.W. and P.V. conceived and designed the study. C.D. assembled the datasets, performed all analyses and wrote the first draft of the manuscript. A.E., R.C., M.J. and K.J.B. contributed to the analytical framework, interpretation of results and manuscript revision. Z.J. contributed to interpretation of results and manuscript revision. Z.W. and P.V. supervised the research. All authors approved the final version.

## Competing interests

The authors declare no competing interests.

## Acknowledgements

We thank the many researchers who contributed biodiversity records to the PREDICTS database and the teams that compiled and maintained the species-level ecological datasets used in this study. We acknowledge the use of OpenAI’s ChatGPT to assist with language editing and to improve the clarity, conciseness and grammatical accuracy of the manuscript. The study conceptualisation, data analyses, interpretation and conclusions are entirely those of the authors, and all AI-assisted text was reviewed and verified by the authors.

## Funding

C.D. discloses support for the research of this work from the PKU–IIASA International Postdoctoral Fellowship Program, jointly funded by Peking University and the International Institute for Applied Systems Analysis. Z.W. discloses support for the research of this work from the National Natural Science Foundation of China [grant numbers 32125026 and 32588202] and the National Key Research and Development Program of China [grant number 2022YFF0802300].

---

**Extended Data Fig. 1.**
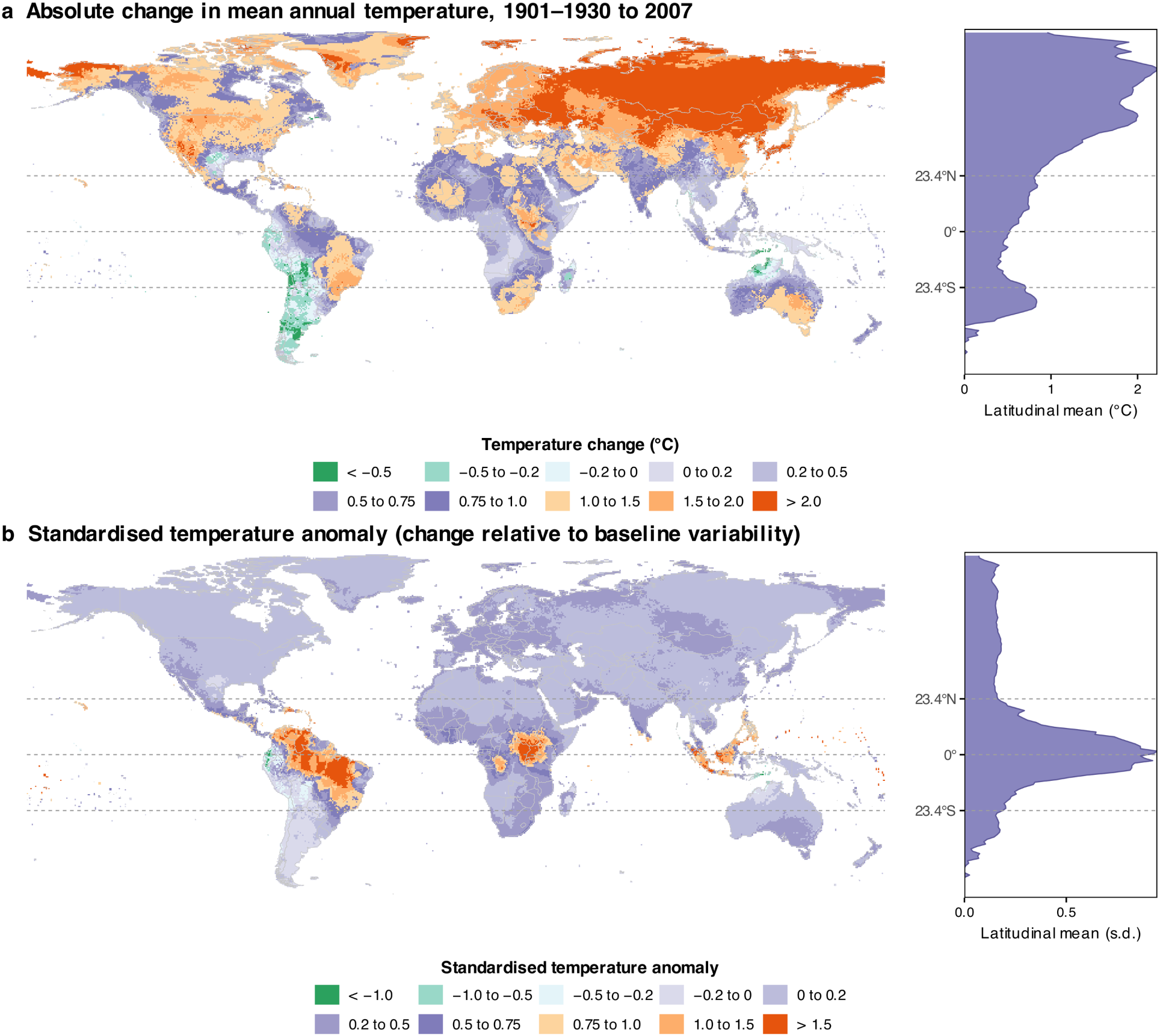
Global average changes in absolute and standardised temperatures. a, Change in mean annual temperature (°C) between the 1901–1930 baseline climatology and 2007 (the median mid-year of biodiversity sampling). b, The same change expressed as the standardised temperature anomaly used throughout our analyses: the absolute change divided by the standard deviation (s.d.) of monthly temperatures during the baseline period. Because standardisation rescales warming by local historical variability, regions with historically stable climates—most notably the humid tropics —show high relative exposure despite moderate absolute warming. Right-hand panels show the latitudinal mean of each metric; dashed lines mark the Equator and the Tropics of Cancer and Capricorn. All values were computed from CRU TS v4.09 monthly mean temperatures; maps show land grid cells (0.5° resolution) between 60° S and 85° N.

**Extended Data Fig. 2.**
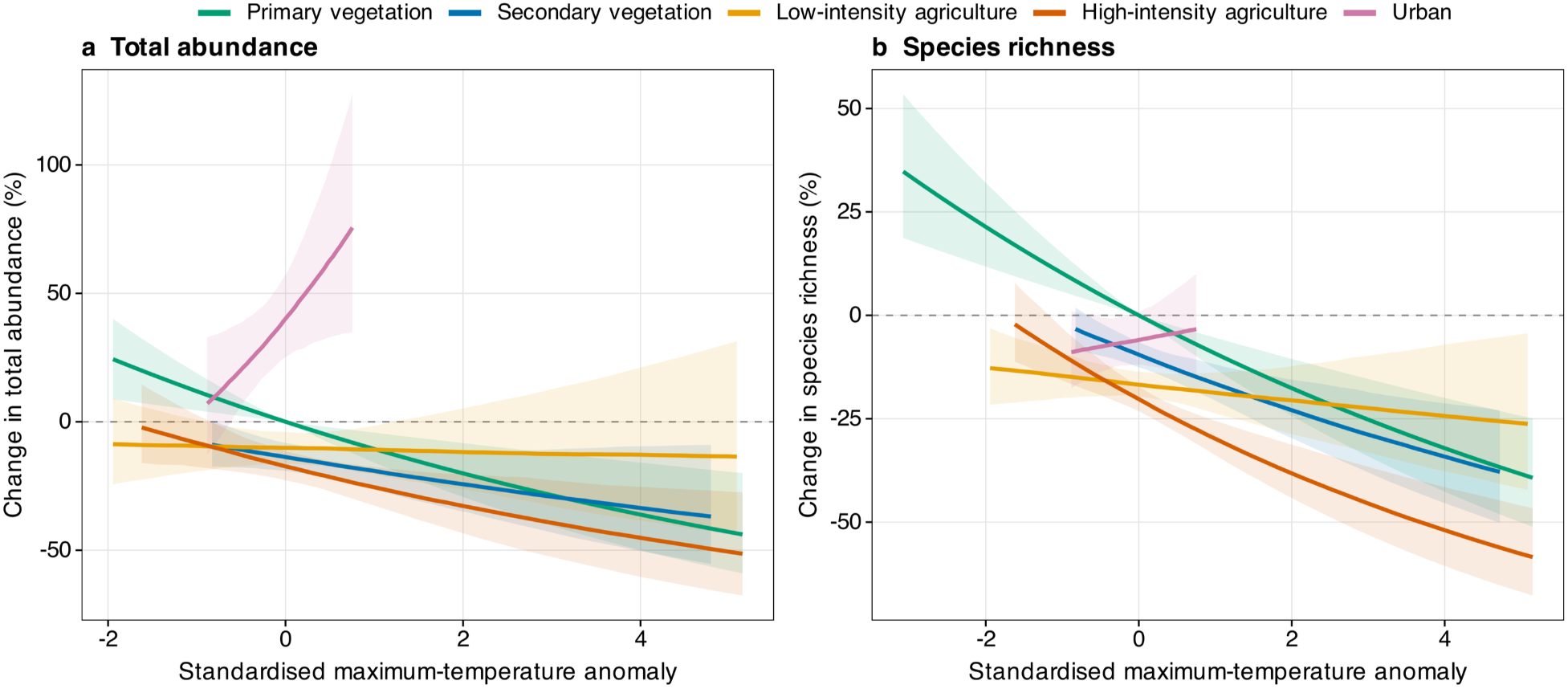
Local-biodiversity responses to the standardised maximum-temperature anomaly. **a**, Total abundance; **b**, species richness. Lines are marginal fixed-effect predictions on the response scale, expressed as the percentage difference from primary vegetation at a standardised maximum-temperature anomaly of zero; shaded bands are 95% confidence intervals obtained by parametric simulation of the fixed-effect variance–covariance matrix (1,000 draws). Colours denote land-use class. Predictions are restricted to the anomaly range observed within each land-use class, so curve lengths differ among classes and reflect data support rather than model truncation. Models are refitted with the maximum-temperature anomaly in place of the mean-temperature anomaly, on 5,817 sites from 121 studies (total abundance) and 6,457 sites from 138 studies (species richness). The omnibus land-use-by-warming interaction was supported for both responses (likelihood-ratio tests: abundance χ²₄ = 19.65, P = 5.84 × 10⁻⁴; richness χ²₄ = 33.12, P = 1.13 × 10⁻⁶).

**Extended Data Fig. 3.**
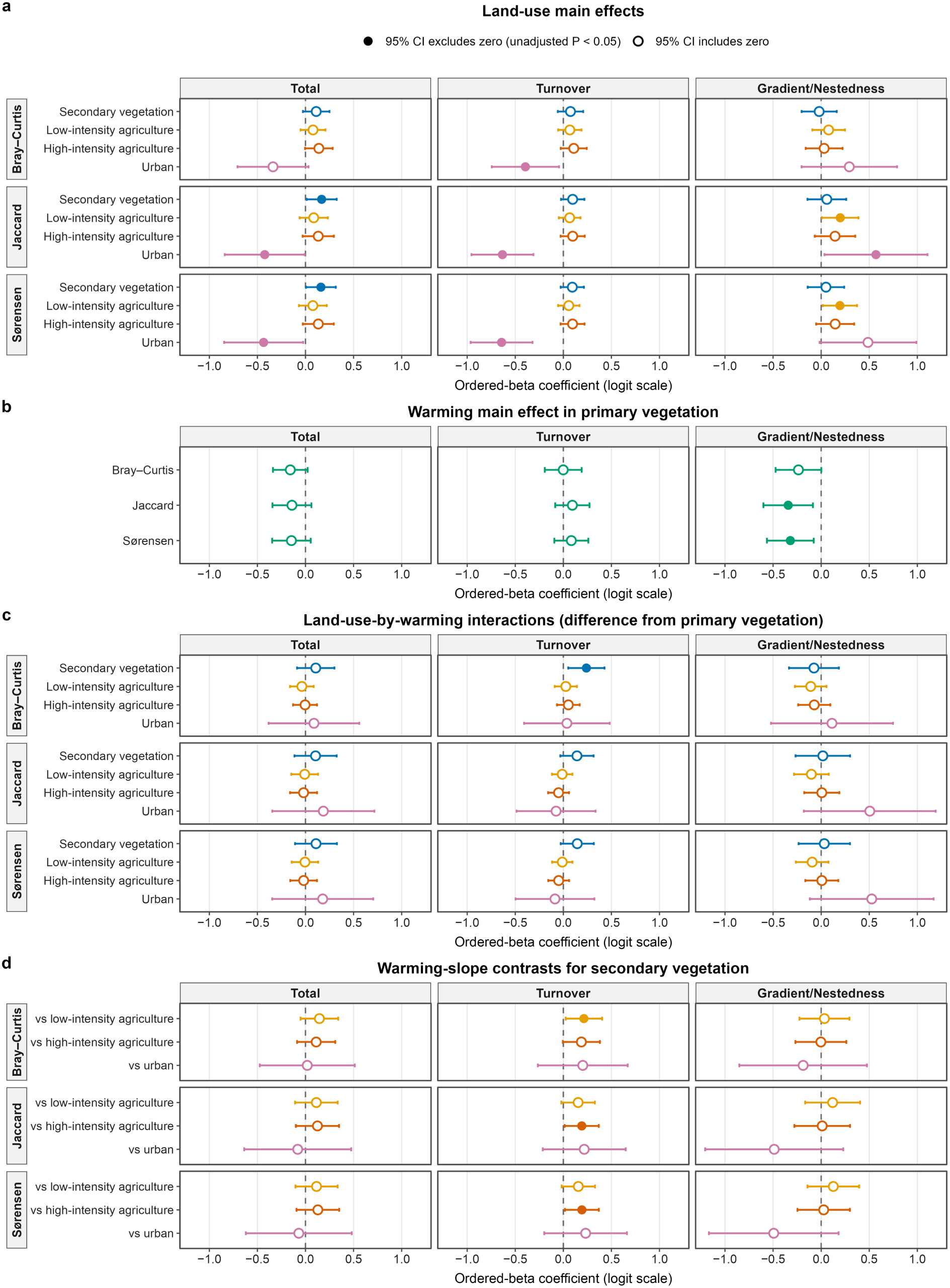
Land-use main effects, the primary-vegetation warming main effect, land-use-by-warming interaction coefficients and warming-slope contrasts for secondary vegetation from the nine ordered-beta dissimilarity models. a, Land-use main effects, that is contrasts of each modified land-use class with primary vegetation at a pair-mean anomaly of zero. b, The warming main effect in primary vegetation, that is the slope of pair-mean standardised temperature anomaly in the reference class. c, Land-use-by-warming interactions, that is the difference between the warming slope in each modified class and that in primary vegetation (omnibus likelihood-ratio tests of the four interaction terms are given in Supplementary Table S2b). d, Differences in warming slope between secondary vegetation and each of the other modified land-use classes, with positive values indicating a more positive slope in secondary vegetation; the corresponding contrast with primary vegetation is the secondary-vegetation interaction in c. Rows within each panel are the three dissimilarity indices (Bray–Curtis, Jaccard, Sørensen) and columns are total dissimilarity and its two additive components (turnover or balanced variation; nestedness-resultant or abundance-gradient). Points are conditional maximum-likelihood estimates on the ordered-beta (logit) model scale and horizontal lines are 95% Wald confidence intervals; filled symbols mark intervals that exclude zero (P < 0.05) and open symbols intervals that include zero. All models were fitted to the same 144,728 within-study, within-land-use site pairs from 88 studies and 4,809 sites.

**Extended Data Fig. 4.**
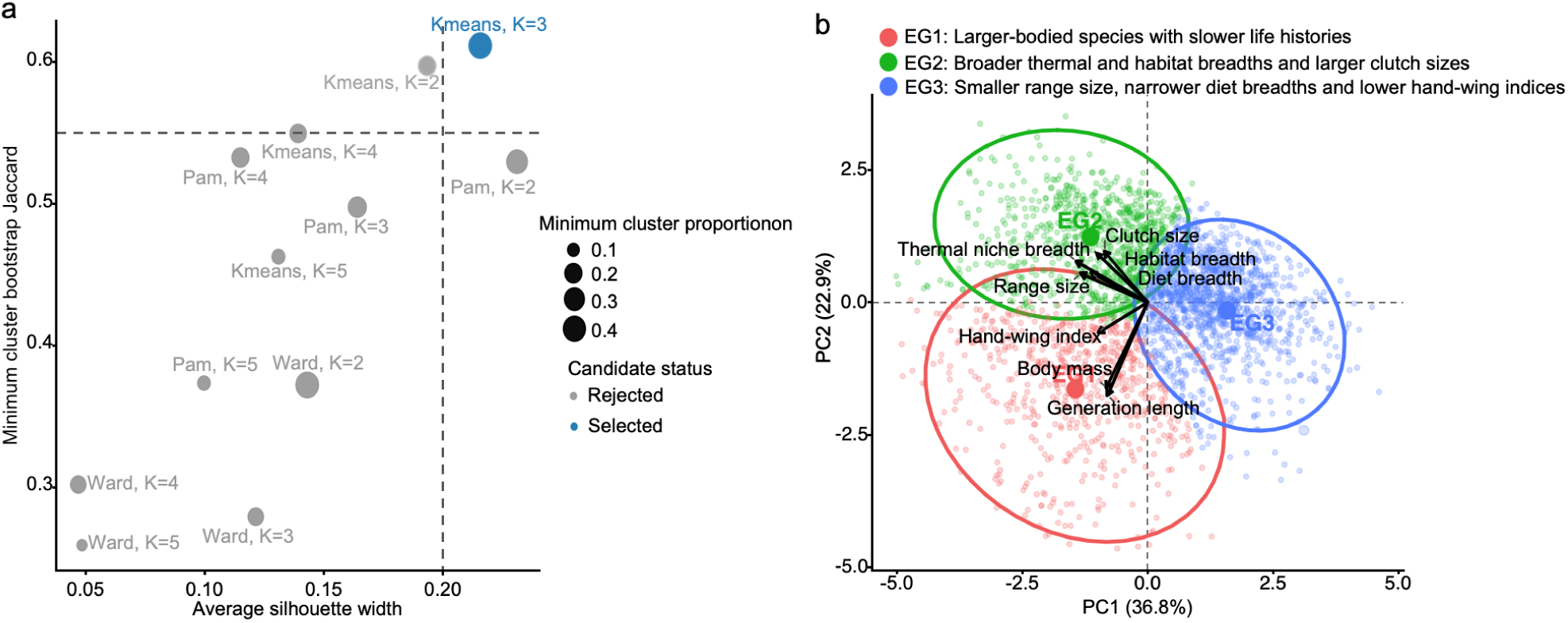
Selection and ecological differentiation of three avian ecological groups. a, Evaluation of candidate clustering solutions across three algorithms: k-means, partitioning around medoids (PAM) and Ward’s hierarchical clustering, and cluster numbers K = 2–5. Points are positioned according to average silhouette width and minimum bootstrap Jaccard stability; point size represents the minimum proportional size of any cluster. Dashed lines indicate the predefined eligibility thresholds, and the retained k-means solution with K = 3 is highlighted in blue. b, Distribution of bird species along the first two principal-component axes derived from eight standardised ecological characteristics. PC1 and PC2 explained 36.8% and 22.9% of the total variation, respectively. Points represent individual species and are coloured by ecological group; ellipses show 95% bivariate-normal envelopes, and arrows indicate the direction and relative contribution of each ecological characteristic. EG1 comprised predominantly larger-bodied species with slower life histories and longer generation lengths (n = 612); EG2 comprised species with broader temperature and habitat niches and larger clutch sizes (n = 954); and EG3 comprised range-restricted species with lower hand-wing indices and narrower diet breadths (n = 1,380).

**Extended Data Fig. 5.**
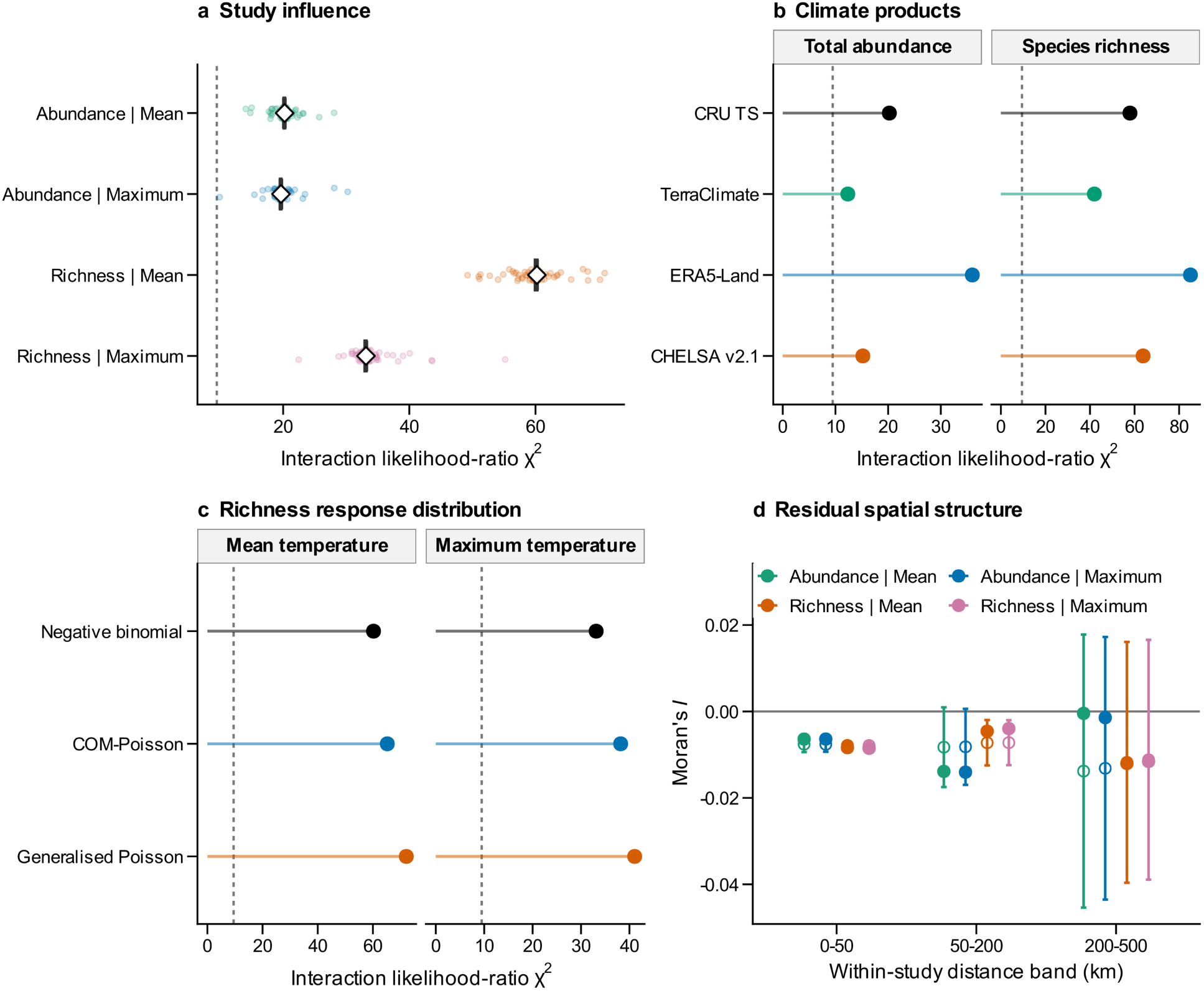
Robustness of the omnibus land-use-by-temperature interaction in the local-biodiversity models. **a**, Study influence. Faint points are the interaction likelihood-ratio χ² from each leave-one-study-out refit (518 refits in total: 121 for each abundance model and 138 for each richness model); open diamonds and vertical bars are the median and interquartile range across refits. **b**, Climate products. Points are the interaction χ² obtained under CRU TS, TerraClimate, ERA5-Land and CHELSA v2.1 using harmonised model structures and complete-case samples (n = 5,750 for abundance; n = 6,386 for richness). **c**, Response distribution. Points are the interaction χ² under negative-binomial, COM–Poisson and generalised-Poisson specifications of the richness model. **d**, Residual spatial structure. Points are Moran’s I of model residuals within studies in three distance bands, with 95% permutation envelopes from 999 within-study permutations; filled symbols denote observed values and open symbols the permutation medians. No within-study site pairs occur beyond 500 km, so wider bands cannot be evaluated. In **a**–**c**, dashed vertical lines mark the 5% critical value of the four-degree-of-freedom χ² distribution (9.49); every refit, product and distribution places the interaction beyond it.

**Extended Data Fig. 6.**
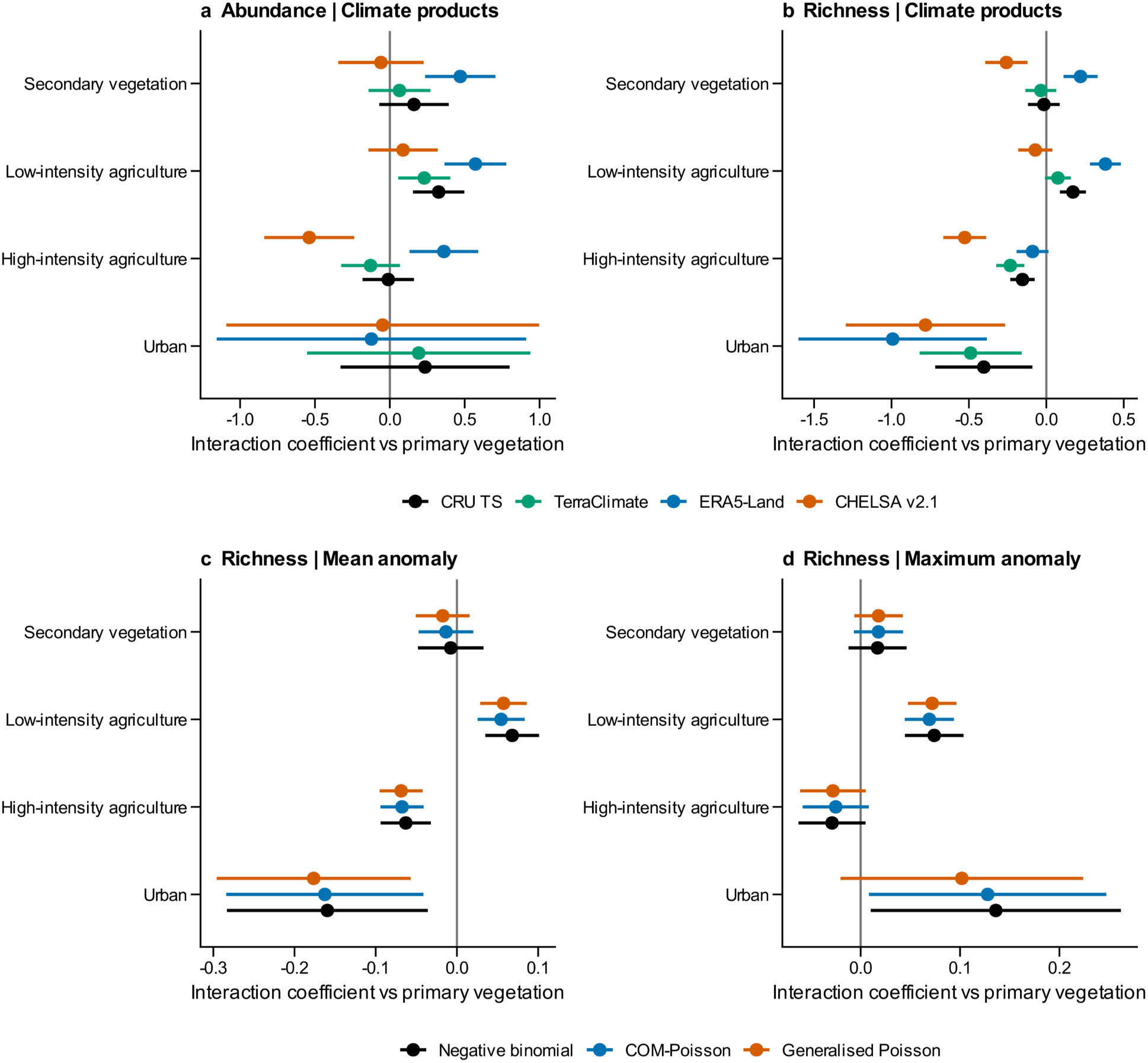
Coefficient-level sensitivity of the land-use-by-warming interaction in the local-biodiversity models. a, b, Interaction coefficients for total abundance (a) and species richness (b) estimated under four gridded climate products, using harmonised model structures and complete-case samples. c, d, Interaction coefficients for species richness under three count distributions, using the mean-temperature anomaly (c) and the maximum-temperature anomaly (d). In all panels, rows are the four modified land-use classes, points are maximum-likelihood estimates of the difference in warming slope relative to primary vegetation, and horizontal lines are 95% Wald confidence intervals; vertical grey lines mark zero. Colours denote climate product (a, b) or count distribution (c, d).

## Notes

https://doi.org/10.5281/zenodo.23237976

### Competing Interest Statement

The authors have declared no competing interest.

