## Supplemental Data 1 for "Land use and climate warming interact to reshape global avian biodiversity through ecological filtering and homogenisation"

Version: 7 October 2026

### Contents


**Supplementary Methods** **4**

S1. Data assembly, filtering and sample-size reconciliation 4

S2. Harmonisation of land-use classes 5

S3. Construction and interpretation of temperature anomalies 6

S4. Background climate and alternative climate products 7

S5. Ecological characteristics and missing-data imputation 9

S6. Definition and validation of ecological groups 9

S7. Local-biodiversity mixed-effects models 10

S8. Assemblage-dissimilarity ordered-beta models 11

S9. Dependence among site-pair comparisons 12

S10. Species-level occurrence model and response-scale decomposition 13

S11. Study influence, response-distribution diagnostics and spatial tests 14

S12. Scope of inference and interpretation boundaries 15

**Supplementary Figures** **16**

Supplementary Fig. S1 16

Supplementary Fig. S2 17

Supplementary Fig. S3 18

Supplementary Fig. S4 19

Supplementary Fig. S5 20

Supplementary Fig. S6 21

Supplementary Fig. S7 22

Supplementary Fig. S8 23

Supplementary Fig. S9 24

Supplementary Fig. S10 25

Supplementary Fig. S11 26

#### Contents (continued)


**Supplementary Tables** **27**

Table S1 27

Table S2 30

Table S3 40

Table S4 45

Table S5 46

Table S6 48

Table S7 49

Table S8 50

Table S9 51

Table S10 52

Table S11 52

Table S12 53

Table S13 54

Table S14 55

Table S15 56

Table S16 58

**Supplementary References** **59**

#### Supplementary Methods

##### S1. Data assembly, filtering and sample-size reconciliation

Bird assemblage records were obtained from PREDICTS version 1.1(Hudson et al., 2014, 2017), and its November 2022 supplementary release (Contu et al., 2022). Records were retained only when predominant land use and use intensity were resolved. Secondary-vegetation stages were pooled because the study tests a common secondary-vegetation class rather than a successional trajectory.

The sampling map contains 6,462 sites. Five sites lacked a CRU TS grid-cell value and were retained only for the descriptive map. The mean-temperature abundance model contains 5,817 sites from 121 studies and 503 spatial blocks; the richness model contains 6,457 sites from 138 studies and 599 blocks. The species-level occurrence model contains 432,560 species-site records from 6,357 sites, 137 studies, 598 spatial blocks and 2,946 species.

The ecological characteristic table and the fitted occurrence model use different denominators, and these are kept separate throughout the tables and figure legends. Eight ecological characteristics were assembled, giving 23,568 potential trait cells. Of these, 176 cells (0.75%) were missing and imputed, affecting 114 species (3.87%). Ecological groups and the fitted occurrence model use the 2,946 species, which is also the sum of the three ecological groups.

#### S2. Harmonisation of land-use classes

PREDICTS land-use and intensity combinations were mapped to five analytical classes: primary vegetation, secondary vegetation, low-intensity agriculture, high-intensity agriculture and urban land. Cropland and pasture under minimal or light use, and plantation forest under minimal use, were classified as low-intensity agriculture. Intense cropland or pasture and light or intense plantation forest were classified as high-intensity agriculture.

The five classes were modelled as nominal categories. Primary vegetation was the reference level, but this parameterisation does not imply an ordered gradient of human modification. Urban sites were not subdivided by intensity because individual urban-intensity combinations provided insufficient empirical support.

#### S3. Construction and interpretation of temperature anomalies

Monthly temperature was extracted from CRU TS v4.09 at 0.5-degree resolution (Harris et al., 2020). For each study, recent exposure was the mean of the 60 months preceding the final sampling year. The primary historical baseline was 1901–1930. The standardised anomaly was the recent-minus-baseline temperature difference divided by the standard deviation of baseline monthly temperatures.

Zero denotes recent temperature equal to the historical baseline mean; negative and positive values are both possible. The covariate therefore describes warming exposure in units of local historical temperature variability, not degrees Celsius. For site-pair models, PairMeanAnom is the mean anomaly of the two sites and absDelta is their absolute anomaly difference.

Most anomaly variation is among studies rather than within studies because paired PREDICTS sites are often geographically close. This structure motivates study-level influence analyses and prevents interpretation of the models as direct temporal trajectories at individual sites.

Support for the exposure gradient. Because PREDICTS studies sample a limited area under a consistent protocol, and because CRU TS has a 0.5-degree grid, the standardised anomaly varies far more among studies than within them. In the species-richness sample, 58 of 138 studies (42.0%) had a within-study span of exactly zero, most often because all sites fell within a single grid cell. The median within-study span was 0.006 anomaly units per study, or 0.015 units for a typical site, and an intercept-only mixed model attributed 98.3% of the variation in site-level anomalies to differences among studies. The corresponding values for the total-abundance sample were 52 of 121 studies (43.0%) and 98.0%. Variation in the maximum-temperature anomaly is somewhat less concentrated among studies (94.4%), but the same structure applies (Table S8; Supplementary Fig. S6).

This is a property of the PREDICTS sampling design rather than of the models, and it has one consequence that applies to every analysis reported here. Warming coefficients and land-use-by-warming interactions are identified largely by contrasts between studies located in regions with different warming histories, rather than by gradients traversed within a single sampling landscape. Because the study random intercept absorbs part of the same contrast, the effective information available for the warming terms is smaller than the total number of sites would suggest, and between-study differences in species pool, biogeography and sampling protocol cannot be separated from warming exposure by design. Estimates are therefore interpreted as regional-scale spatial associations (Supplementary Methods S12).

#### S4. Background climate and alternative climate products

The standardised anomaly measures how far recent temperatures at a site have departed from that site's own 1901–1930 baseline, expressed in units of its baseline variability. It is therefore a relative measure: two sites can share the same anomaly while differing widely in absolute mean temperature. Because absolute background temperature is itself a correlate of avian assemblage composition and covaries with latitude, elevation, biome and the global distribution of land-use classes, an association attributed to warming exposure could instead reflect position on the global temperature gradient. The land-use-by-anomaly interaction is vulnerable to the same confounding.

To test this, both local-biodiversity models were refitted with the site-level 1901–1930 climatological mean annual temperature, standardised across sites and added as a single additive covariate. This quantity is taken from the same CRU TS v4.09 series and the same baseline period used to derive the anomalies, so it describes the historical thermal setting of each site and, unlike a contemporary climatology, contains no late-twentieth-century warming signal. The land-use classification, the land-use-by-anomaly interaction, the random-effects structure and the response distributions were identical to the primary models, so any change in the focal interaction is attributable to the covariate. The comparison is diagnostic rather than inferential: if the focal interaction were an artefact of position on the global temperature gradient, adding background temperature should attenuate the interaction coefficients or remove their statistical support.

The baseline mean annual temperature is defined for every site with an anomaly, so the adjusted models retain the complete samples of 5,817 abundance and 6,457 richness observations (site mean 16.02 and 15.80 °C; s.d. 8.72 and 8.59). The omnibus interaction remained supported for abundance ($\chi^2$ = 20.13, d.f. = 4, P = 4.71 × $10^{-4}$) and richness ($\chi^2$ = 59.81, d.f. = 4, P = 3.19 × $10^{-12}$), and the principal interaction coefficients retained their directions and statistical support (Table S11; Supplementary Fig. S7). Thus, adjustment for absolute macroclimatic context did not account for the focal interaction.

This test addresses confounding by long-term background macroclimate only. It does not exclude confounding by variables that covary with warming exposure at finer spatial or thematic scales, including microclimate, land-use history, management intensity, habitat quality and biotic interactions, and it does not convert the space-for-time design into a temporal inference.

Climate-product sensitivity used harmonised complete-case samples and identical model structures for CRU TS, TerraClimate, ERA5-Land and CHELSA (Abatzoglou et al., 2018; Muñoz-Sabater et al., 2021; Karger et al., 2017, 2021). Product choice jointly changes time-series length, feasible baseline period, gridding method and spatial resolution. It is therefore not interpreted as a resolution-only test.

On common complete-case samples, the omnibus interaction was supported under all four products for abundance (n = 5,750; $\chi^2$ = 12.35–35.98; P ≤ 0.0149) and richness (n = 6,386; $\chi^2$ = 41.95–85.08; P ≤

$1.71 \times 10^{-8}$), whereas several individual land-use contrasts were product-sensitive. This distinction is retained in the Results: climate-product choice did not alter the overall conclusion that warming associations varied among land-use classes, but it did affect the magnitude or support of some class-specific coefficients.

#### S5. Ecological characteristics and missing-data imputation

Eight ecological characteristics were assembled. Body mass, geographic range size and clutch size were obtained from AVONET (Tobias et al., 2022); habitat breadth followed Etard and Newbold (2024); realised thermal niche breadth followed Williams and Newbold (2021); hand-wing index followed Sheard et al. (2020); and generation length followed Bird et al. (2020). Diet breadth is the number of major dietary categories used, derived from the BIRDBASE diet-breadth variable (Sekercioglu et al., 2025). Missing values occurred only for realised thermal niche breadth (113 species) and diet breadth (63 species). In total, 176 of 23,568 trait cells (0.75%) were missing across 113 of 2,946 species (3.84%).

Missing values were imputed with missForest (Stekhoven and Bühlmann, 2012). Internal out-of-bag NRMSE was recorded, and predictive accuracy was evaluated independently using five repeats of ten-fold masked cross-validation. Cross-validated NRMSE was 0.45 for realised thermal niche breadth and 0.62 for diet breadth, with corresponding observed–predicted Pearson correlations of 0.91 and 0.82. These results indicate good predictive performance for both ecological characteristics.

#### S6. Definition and validation of ecological groups

Ecological characteristics were transformed as specified in the analysis code, winsorised at the 1st and 99th percentiles and standardised. Principal-component scores were clustered using k-means, partitioning around medoids and Ward hierarchical clustering for K = 2–5. Candidate solutions were assessed from silhouette width, minimum bootstrap Jaccard stability and the size of the smallest group.

The retained eight-characteristic solution was k-means with K = 3 (silhouette width 0.215; minimum bootstrap Jaccard stability 0.607; minimum group proportion 0.394). EG1, EG2 and EG3 contained 612, 954 and 1,380 species, respectively. PC1 and PC2 explained 36.8% and 22.9% of variance (Table S10).

#### S7. Local-biodiversity mixed-effects models

Within-study rescaled and log-transformed total abundance was modelled with a Gaussian linear mixed-effects model. Species richness was modelled with a negative-binomial mixed-effects model under the nbinom2 parameterisation and a log link. Both models included land use, standardised temperature anomaly and their interaction, together with random intercepts for study and spatial block.

The primary interaction was evaluated by a four-degree-of-freedom likelihood-ratio test comparing the full interaction model with a nested model that retained both main effects. Land-use-specific warming slopes were obtained as linear combinations of the primary- vegetation anomaly coefficient and the corresponding interaction coefficient. Maximum-temperature models were fitted as complementary exposure analyses.

For mean-temperature anomaly, the omnibus interaction was supported for total abundance ($\chi^2 = 20.23$, d.f. = 4, $P = 4.50 \times 10^{-4}$) and species richness ($\chi^2 = 60.22$, d.f. = 4, $P = 2.61 \times 10^{-12}$). Full coefficients, confidence intervals, random-effect variances and land-use-specific slopes are given in Table S1.

**S8. Assemblage-dissimilarity ordered-beta models**

Bray-Curtis, Jaccard and Sorensen total dissimilarity and their two additive components were calculated for pairs of sites sampled within the same study and land-use class (Baselga, 2010, 2012, 2013). Exact zeros and ones were retained with ordered-beta mixed-effects models using a logit link (Kubinec, 2023; Brooks et al., 2017).

For sites i and j, the conditional model was D_ij ~ LandUse_ij × PairMeanAnom_ij + |ClimDiff_ij| + log(GeoDist_ij) + (1|Study) + (1|Site_i) + (1|Site_j). Random intercepts account for study heterogeneity and repeated occurrence of each site in multiple pairwise comparisons.

The primary Bray-Curtis data contained 144,728 complete site pairs from 88 studies and 4,809 sites. A negative within-land-use warming slope indicates increasing compositional similarity with warming exposure. The land-use-by-warming interactions were not supported for any of the nine dissimilarity responses; class-specific slopes are therefore described without claiming that they differ from primary vegetation (Table S2).

Support for the focal predictor. Restricting comparisons to sites within the same study and land-use class makes eligible pairs both geographically and climatically proximate. Of the 144,728 pairs entering the fitted models, 94.5% were separated by no more than 100 km, with a median separation of 10.2 km, and 84.8% differed by no more than 0.05 standardised anomaly units, that is by less than one-twentieth of a local baseline standard deviation.

Two consequences follow. First, |ClimDiff_ij| has limited leverage: within-pair climatic separation is small for most comparisons, so this term is estimated over a narrow range and functions as a control for residual climatic separation rather than as a substantive test of climatic divergence. Second, the focal predictor is effectively a study-level attribute. Site-level anomalies vary far more among studies than within them: 58 of 138 studies had a within-study span of zero, most often because all sites fell within the same 0.5-degree grid cell; the median within-study span was 0.006 anomaly units per study and 0.015 units for a typical site; and variance partitioning attributed 98.3% of the variation to differences among studies (Table S8; Supplementary Fig. S6). Pair-level mean anomaly averages two such sites and inherits this structure.

The warming-dissimilarity associations are therefore identified largely by contrasts between studies in regions with different warming histories, rather than by gradients traversed within a single sampling landscape. Because the study random intercept absorbs part of the same contrast that identifies the focal term, the effective information available for these estimates is limited. Weak support for most terms should be read as limited resolution under this design rather than as evidence that warming is unrelated to assemblage composition (Supplementary Methods S12).

#### S9. Dependence among site-pair comparisons

Repeated strictly disjoint-pair subsampling addresses dependence caused by shared sites. In each of 1,000 iterations, sites were paired without replacement within each study-by- land-use stratum, so that each site contributed to at most one comparison. The nine ordered-beta models were refitted with a study random intercept, and empirical medians and 2.5th-97.5th percentile intervals were calculated (Table S3a; Supplementary Fig. S1).

Separately, study-level cluster bootstrap resampling assessed sensitivity to the finite and uneven representation of studies. Studies were sampled with replacement in 1,000 iterations and Gaussian mixed models of bounded logit-transformed dissimilarity were refitted. Leave-one-study-out refits of the same Gaussian-logit sensitivity models were used to identify influential studies (Tables S3b,c; Supplementary Fig. S2). These analyses are distributional and dependence sensitivities, not replacement confidence intervals for the primary ordered-beta models.

**S10. Species-level occurrence model and response-scale decomposition**

Occurrence was modelled with a binomial-logit mixed model containing the three-way interaction among land use, standardised mean-temperature anomaly and ecological group. Random intercepts were fitted for study, spatial block, site and species. The primary block-augmented model had AIC 283,795, compared with 284,013 for the model without the spatial-block term.

Predictions were calculated on the probability scale for each ecological-group-by-land- use combination. Land-use effects at anomaly zero, within-land-use warming responses and departures from the additive expectation were kept distinct. Uncertainty was propagated from the full fixed-effect variance-covariance matrix.

The block and no-block models yielded the same qualitative classification for all reported response-scale departures. The block model is therefore retained as primary because it matches the hierarchical design and has the lower AIC.

**S11. Study influence, response-distribution diagnostics and spatial tests**

Each of the four local-diversity models was refitted after omitting one study at a time. All 518 refits were inferentially usable, and the omnibus land-use-by-anomaly interaction remained supported in every omission. Coefficient-level support was less uniform, so omnibus and individual-coefficient stability are reported separately (Table S4; Fig. S3).

DHARMa diagnostics used 500 simulations (Hartig, 2022). Abundance models showed no dispersion departure but non-uniform residuals and more outliers than expected. Negative-binomial richness models showed non-uniform residuals and marked underdispersion. Residual medians also varied among land-use strata, and residual-anomaly associations were modest but detectable in three of the four models, particularly the richness models and the maximum-temperature analyses. Conway–Maxwell–Poisson (COM-Poisson) and generalised-Poisson sensitivity models retained the focal omnibus interactions and the principal mean-temperature coefficient directions, but did not fully remove the diagnostic departures. We therefore retained the pre-specified negative-binomial models for comparability with the primary PREDICTS workflow while treating agreement across count families as the relevant robustness evidence and explicitly acknowledging incomplete residual fit (Tables S6, S13 and S14; Supplementary Figs. S5, S9 and S10).

Moran tests used Pearson residuals and 999 within-study permutations. No residual spatial autocorrelation was detected at 0–50, 50–200 or 200–500 km after multiplicity adjustment. No within-study site pairs occurred at 500–2,000 km, so no inference was made for that distance band (Table S7).

#### S12. Scope of inference and interpretation boundaries

PREDICTS is a globally replicated space-for-time database rather than a longitudinal monitoring network. The fitted associations therefore describe biodiversity variation along spatial gradients in recent warming exposure and land-use context. They do not directly estimate temporal population trajectories, causal warming effects, extinction debts or colonisation credits.

For RQ1, a supported omnibus interaction establishes that warming associations vary among land-use classes; it does not imply that every interaction coefficient is supported. For RQ2, a supported slope within one class does not establish a difference from primary vegetation when the corresponding interaction or omnibus test is unsupported.

Predictions should be restricted to observed covariate support. Urban ecological-group combinations contain fewer observations and narrower anomaly ranges than several other combinations, so estimates near the edge of urban support are interpreted cautiously.

### Supplementary Figures

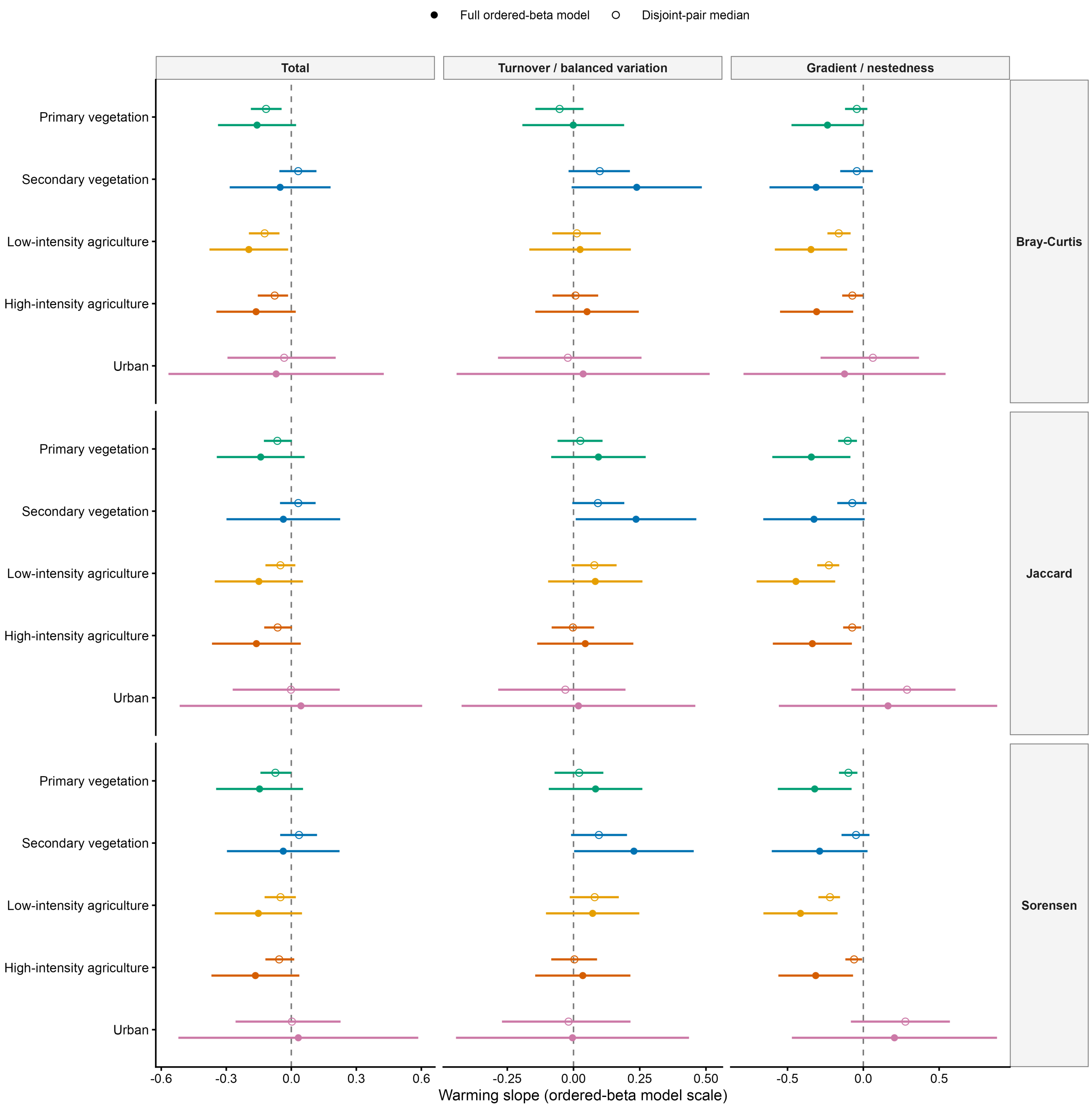


**Supplementary Fig. S1 |** Repeated strictly disjoint-pair subsampling for the nine ordered-beta assemblage-dissimilarity models. Filled circles and lines show full-data land-use-specific warming slopes and 95% Wald confidence intervals; open circles and lines show medians and 2.5th-97.5th percentiles across 1,000 subsamples in which each site entered at most one pair. The vertical line denotes zero.

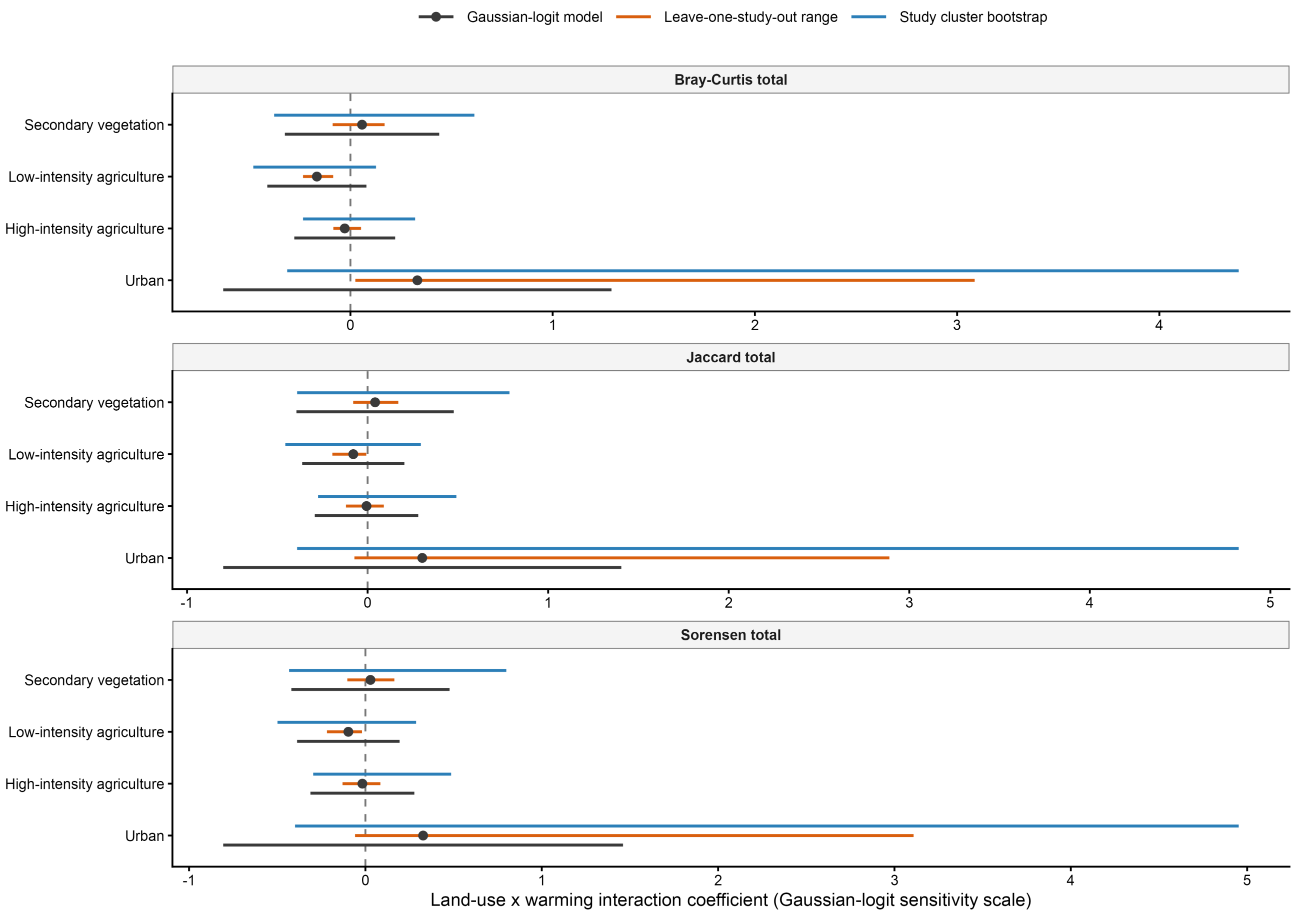


**Supplementary Fig. S2 |** Study-level dependence and influence sensitivity for total Bray-Curtis, Jaccard and Sorensen dissimilarity. Diamonds show coefficients and 95% Wald confidence intervals from Gaussian mixed models of bounded logit-transformed dissimilarity; thick lines show leave-one-study-out ranges; open circles and thin lines show medians and 2.5th-97.5th percentiles from 1,000 study-cluster bootstrap resamples. These estimates are on the Gaussian-logit sensitivity scale and are not confidence intervals for the primary ordered-beta models.

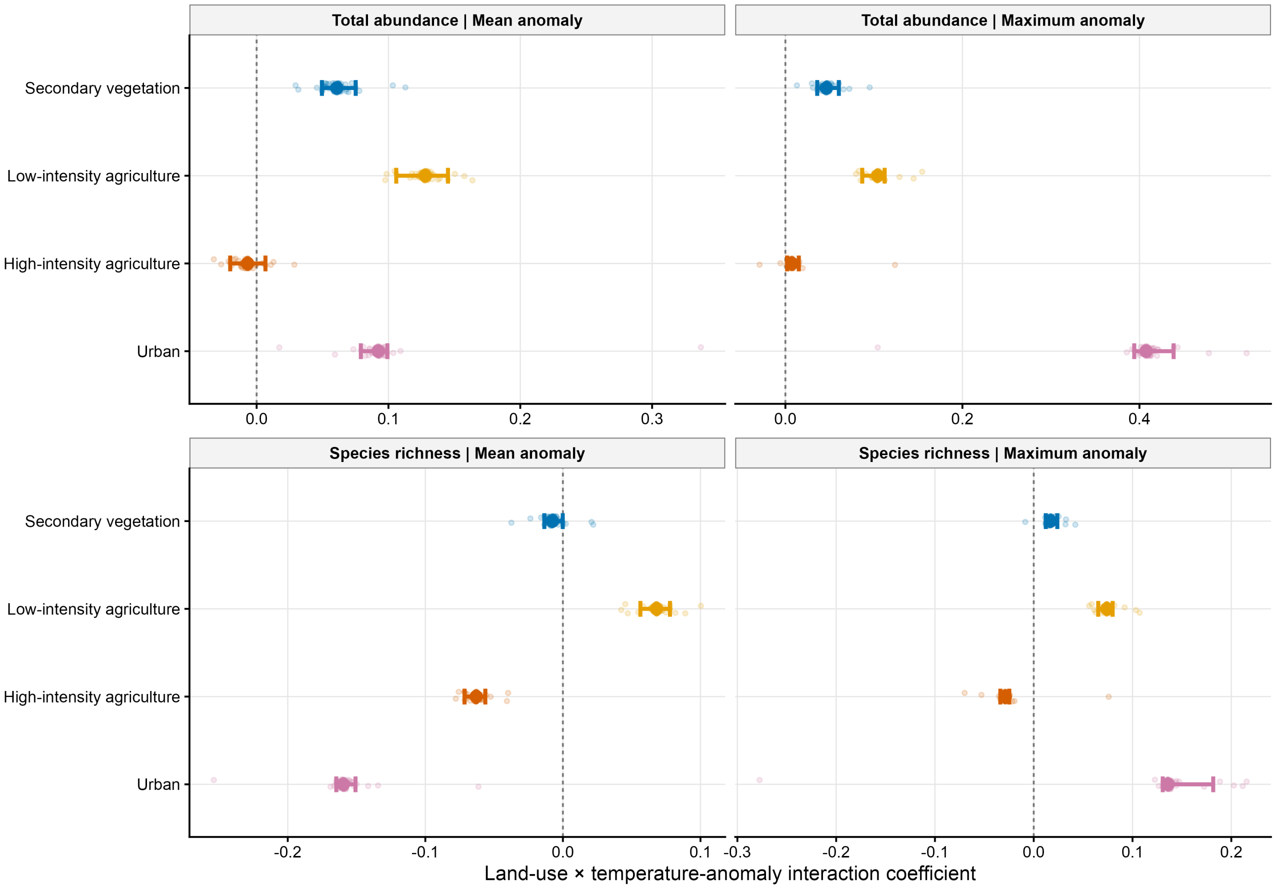


**Supplementary Fig. S3 |** Leave-one-study-out envelopes for the land-use-by-temperature interaction coefficients in the four local-diversity models.

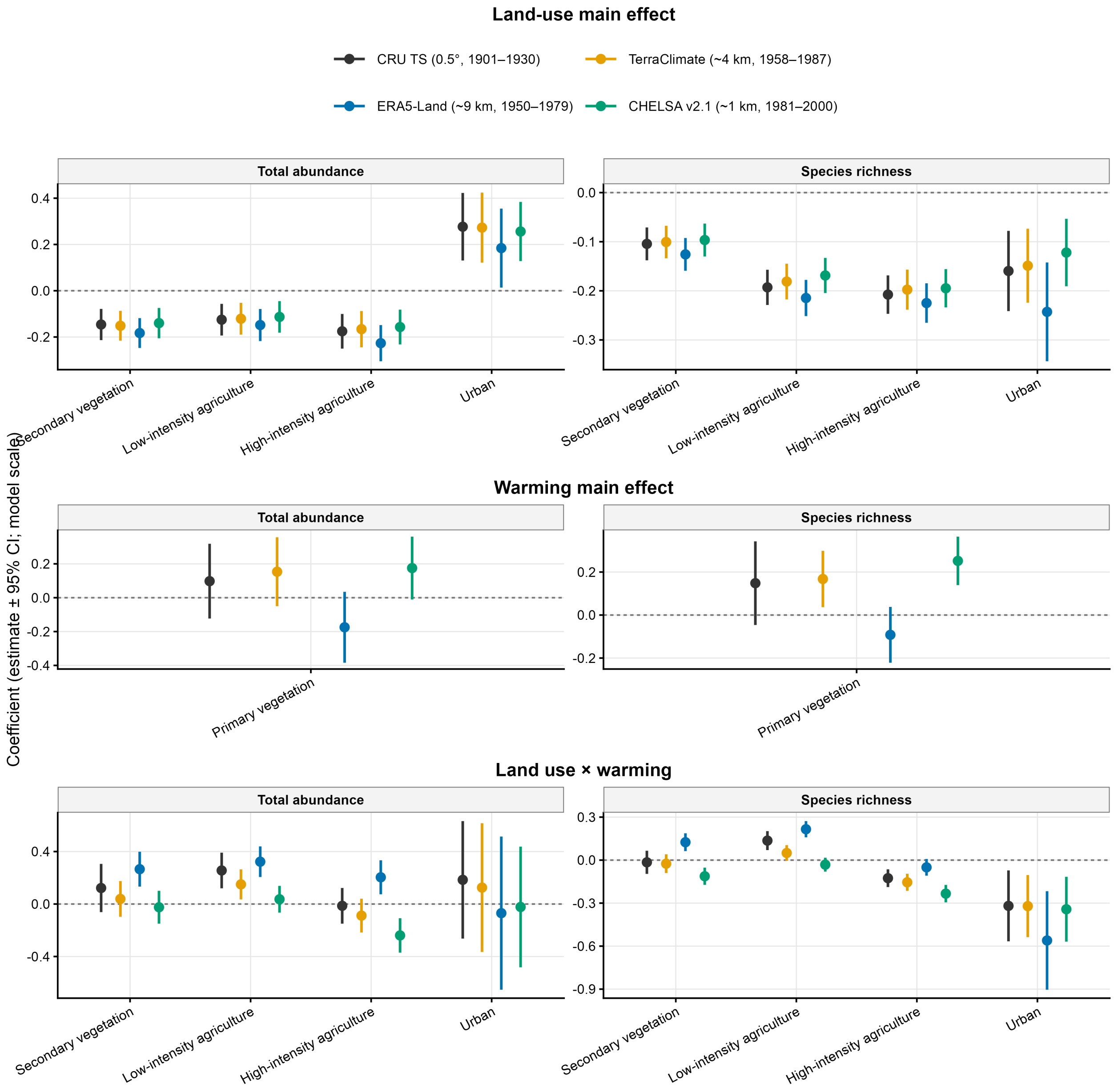


**Supplementary Fig. S4 |** Climate-product sensitivity of local-biodiversity coefficients under harmonised model structures and complete-case samples.

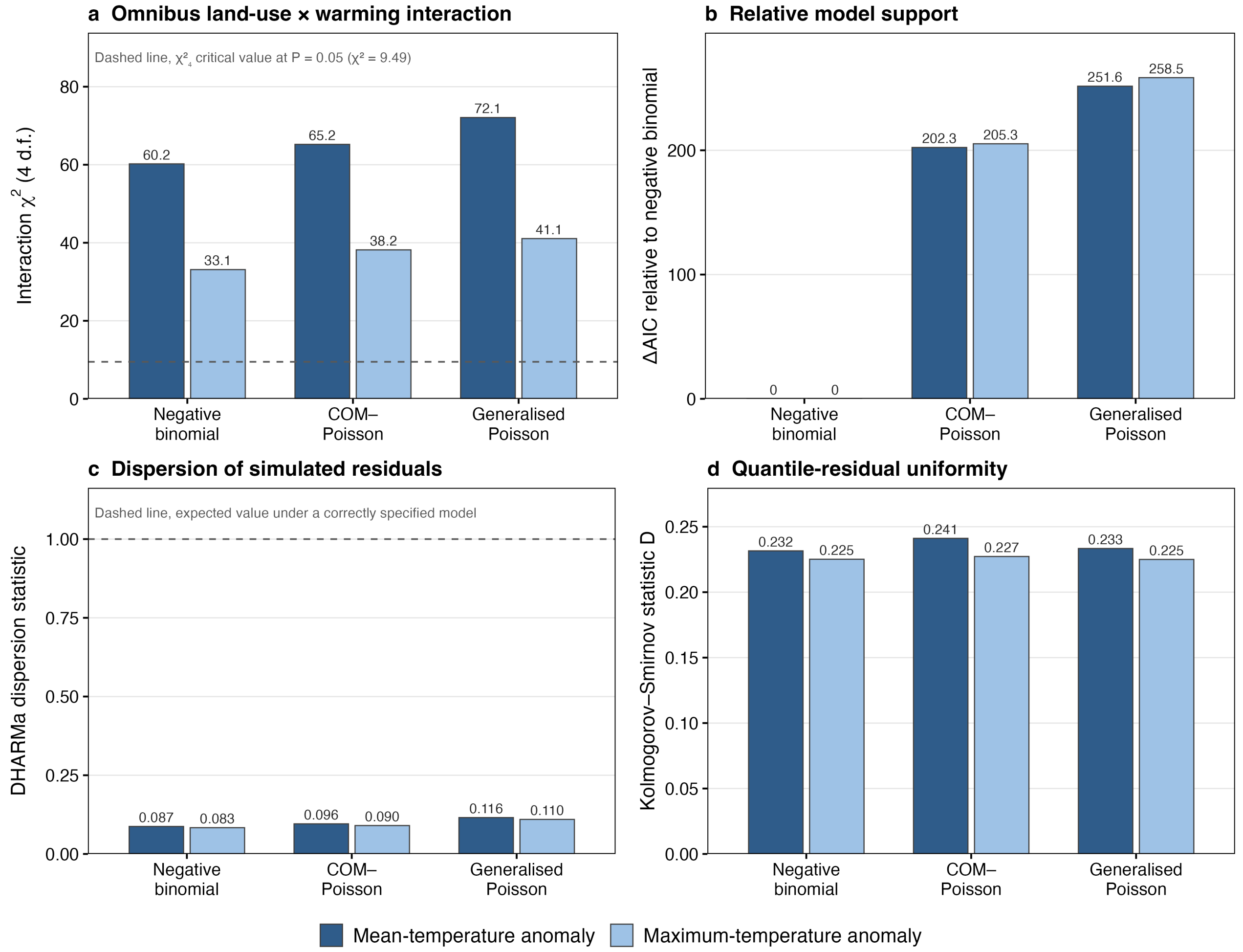


**Supplementary Fig. S5 |** Comparison of negative-binomial, COM-Poisson and generalised-Poisson richness models. Alternative families retain the focal interaction but do not eliminate all residual departures.

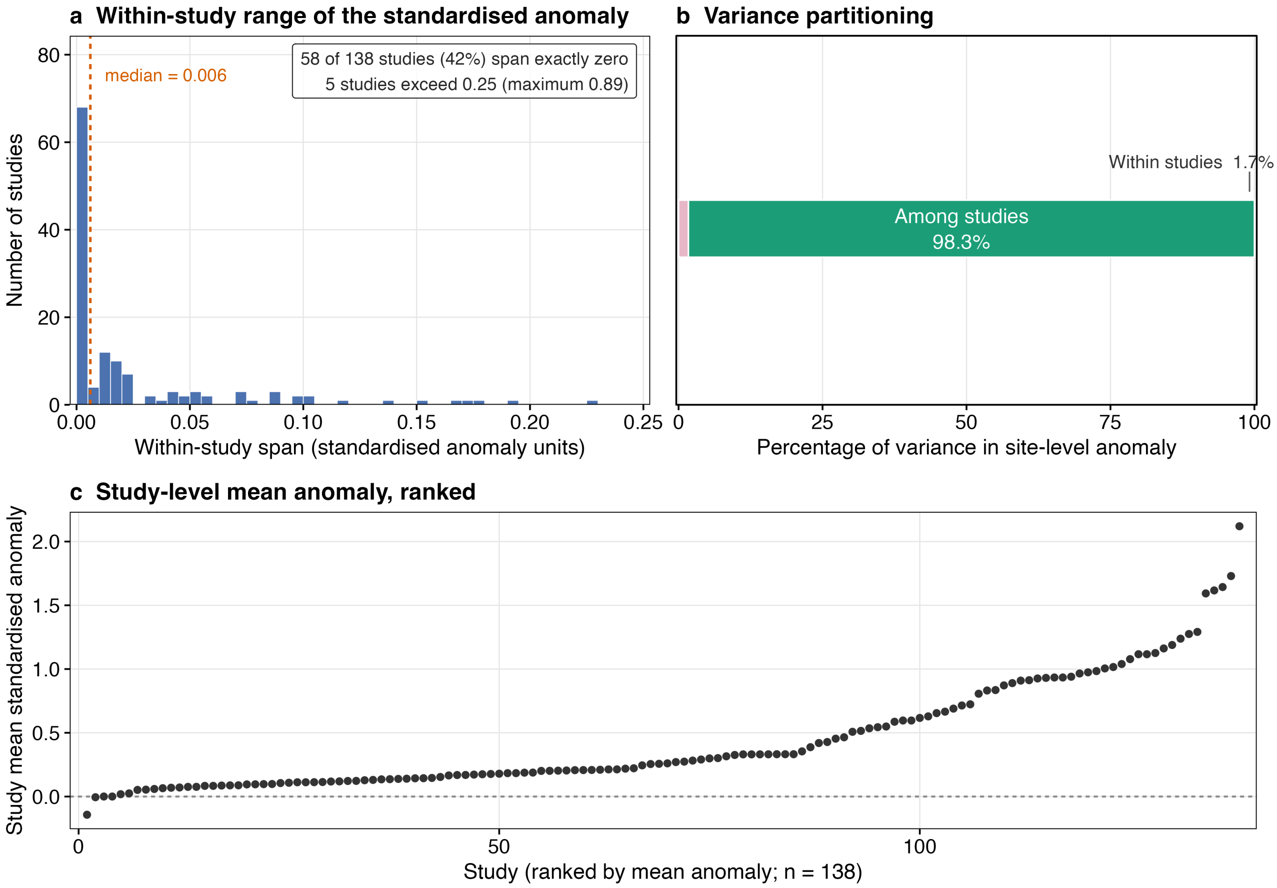


**Supplementary Fig. S6 |** Distribution of the standardised temperature anomaly within and among studies. Most temperature-exposure variation occurs among rather than within studies: 58 of 138 studies had a within-study span of exactly zero, the median within-study span was 0.006 anomaly units per study, and an intercept-only mixed model attributed 98.3% of the variation in site-level anomalies to differences among studies (Table S8).

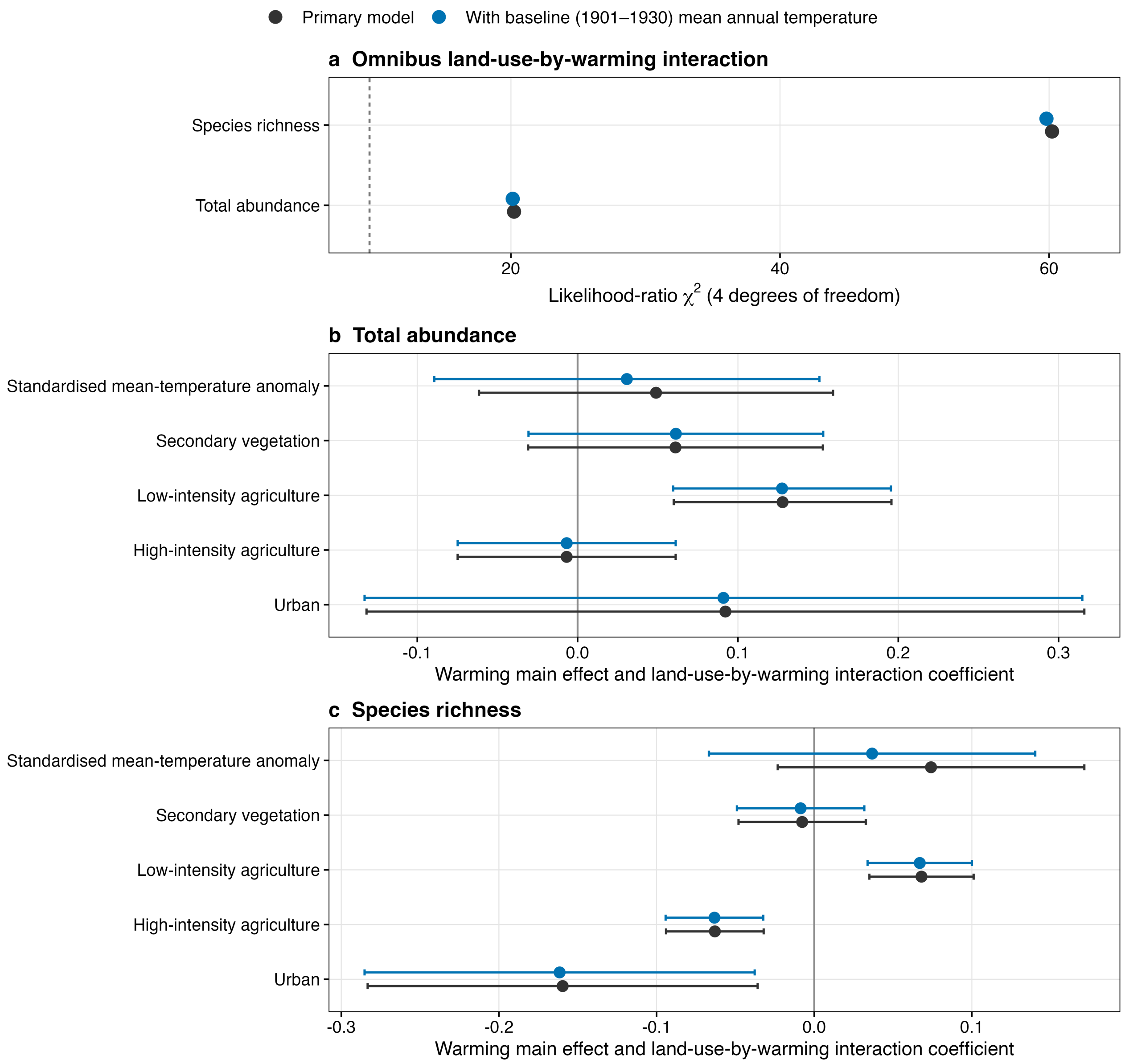


**Supplementary Fig. S7 |** Sensitivity of the local-biodiversity interactions to adjustment for the site-level 1901–1930 climatological mean annual temperature. Points and lines show estimates and 95% confidence intervals before and after adjustment. The covariate is defined for every site with an anomaly, so the adjusted models retain the complete samples of 5,817 abundance and 6,457 richness observations. The omnibus interaction remained supported for both responses, and no interaction coefficient changed sign or crossed the conventional threshold of statistical support.

a

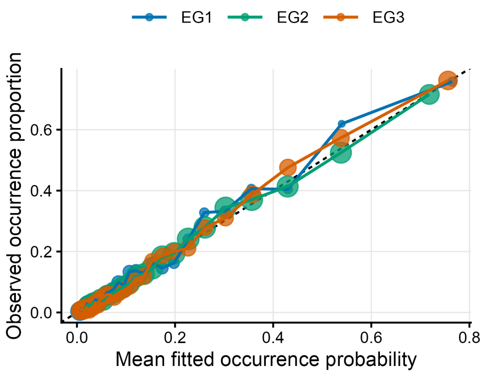


b

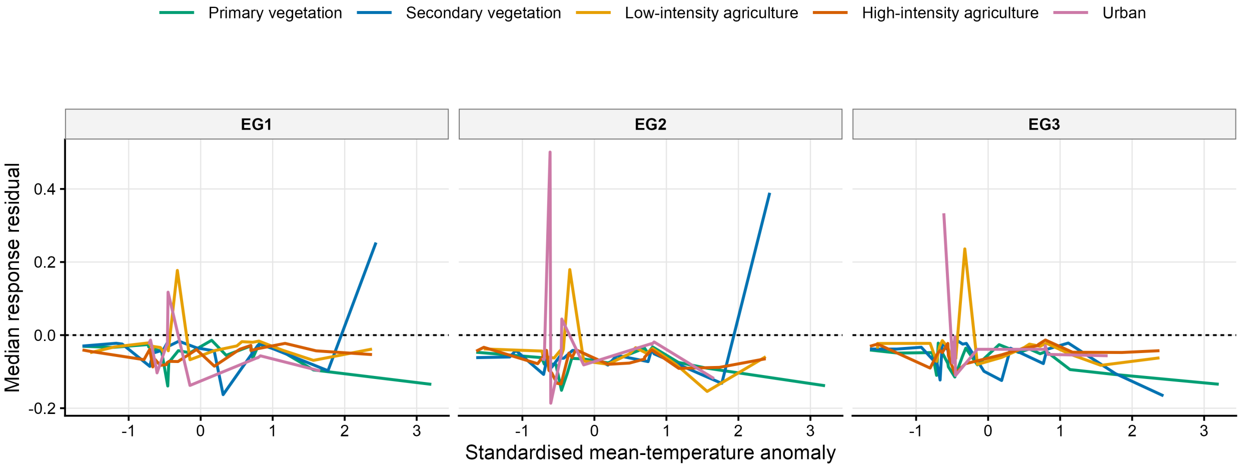


**Supplementary Fig. S8 |** Calibration and residual patterns for the block-augmented species-occurrence model. Calibration is summarised by ecological group; residual patterns are shown across land-use classes and warming exposure.

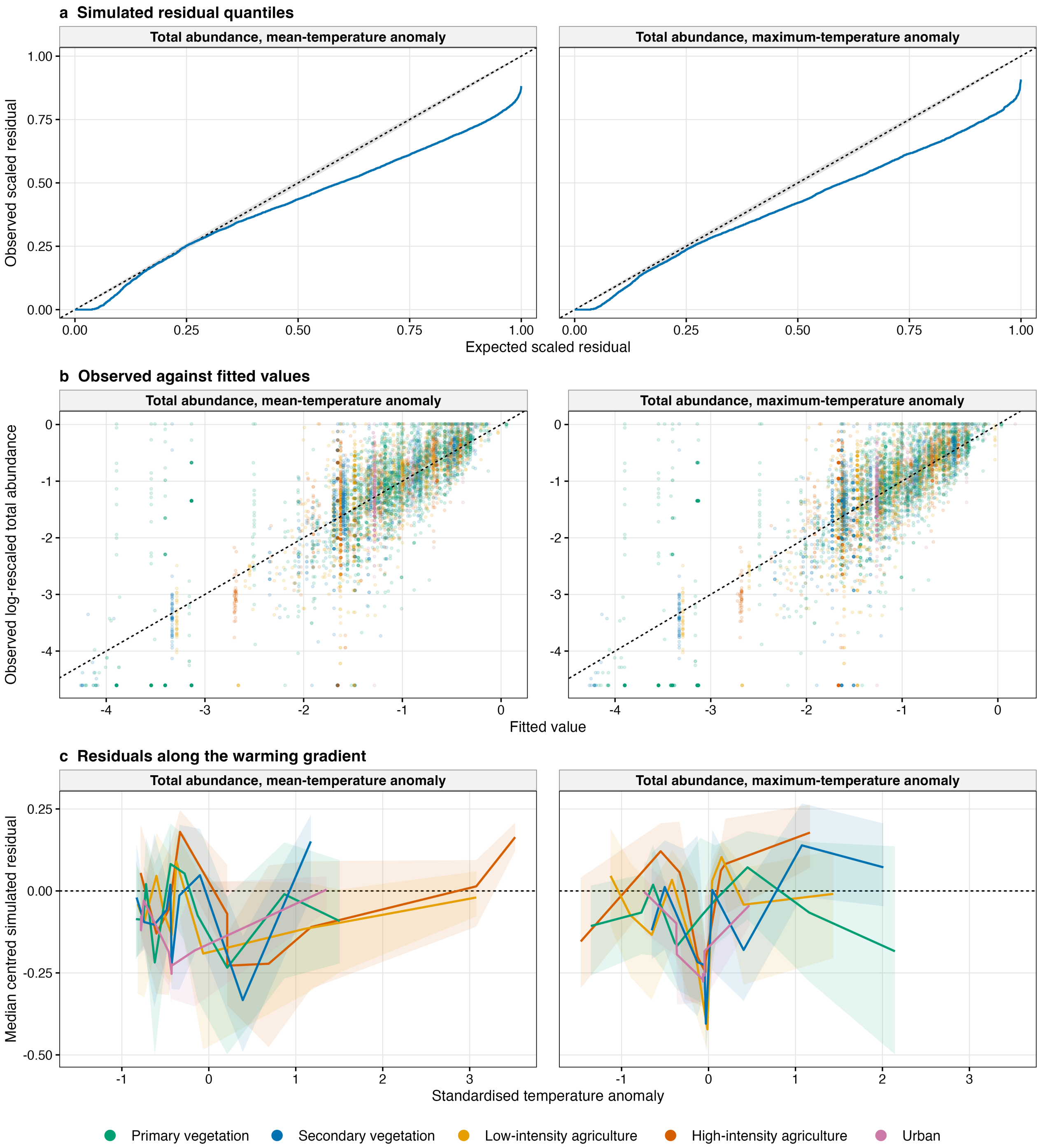


**Supplementary Fig. S9 |** Diagnostics for total-abundance models using mean- and maximum-temperature anomalies. Panels show scaled-residual quantiles, observed versus fitted values and binned residual patterns across land use and anomaly.

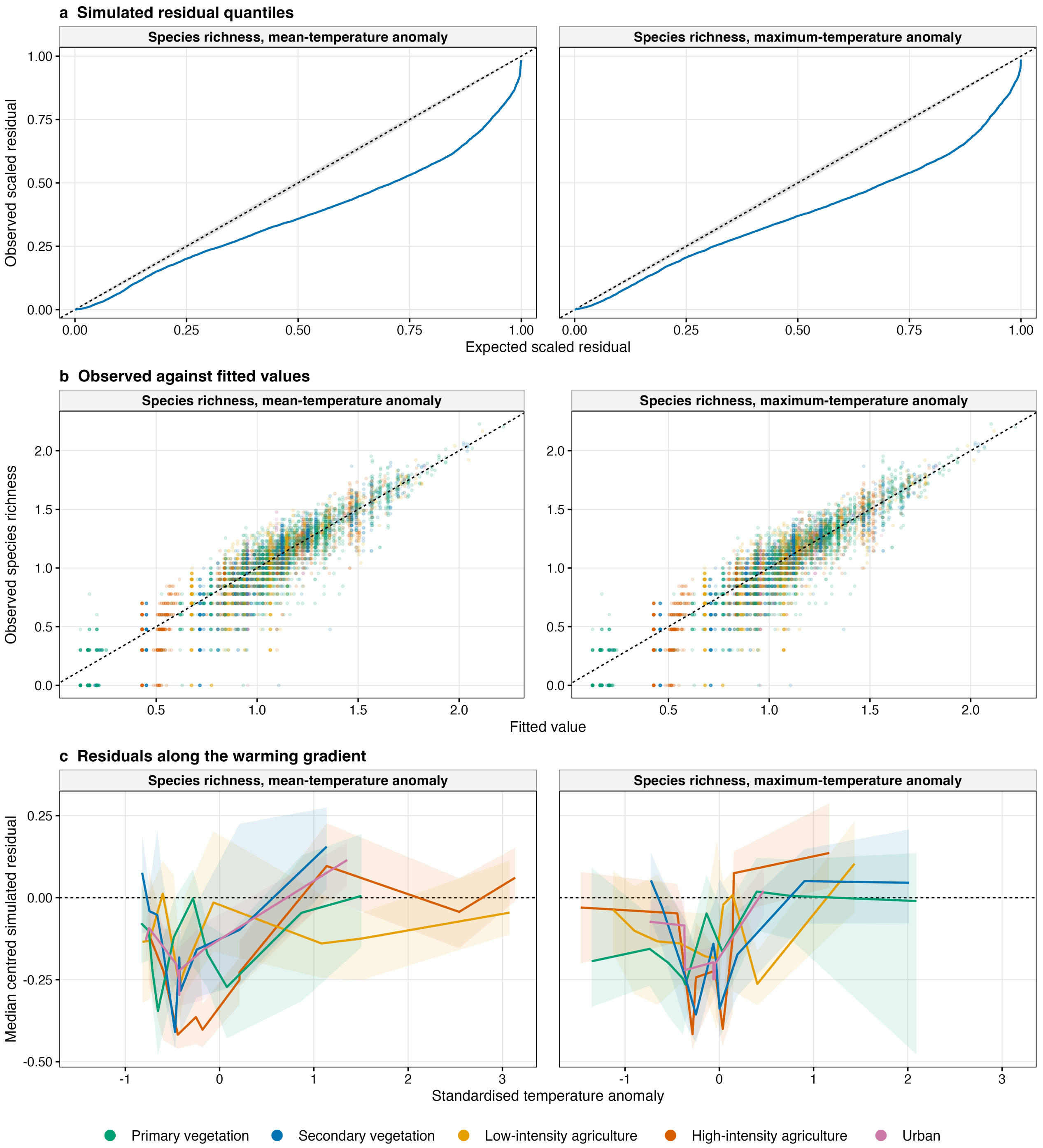


**Supplementary Fig. S10 |** Diagnostics for species-richness models using mean- and maximum-temperature anomalies. Departures from the assumed negative-binomial distributions are retained explicitly.

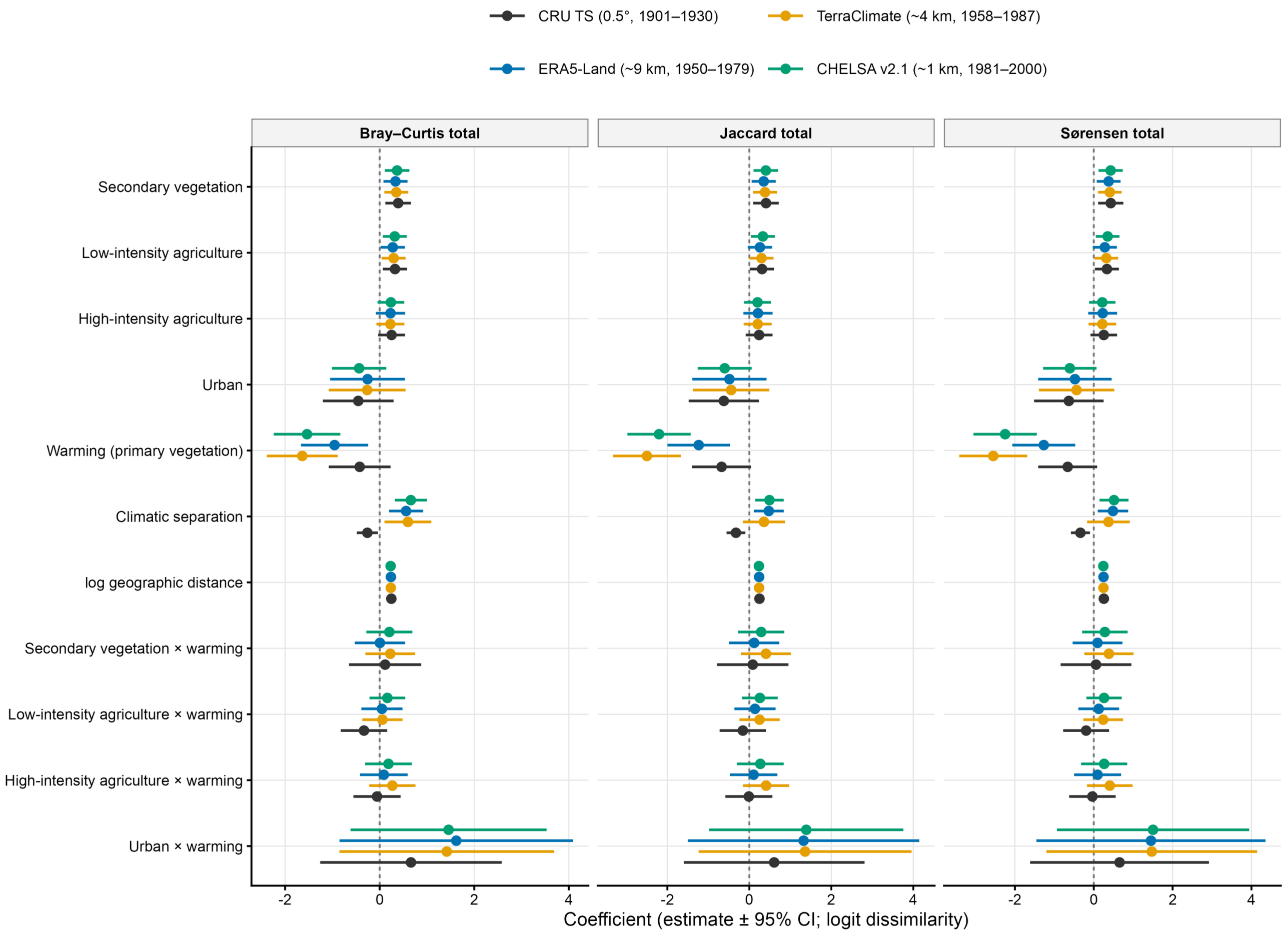


**Supplementary Fig. S11 |** Climate-product sensitivity of assemblage-dissimilarity associations.

### Supplementary Tables

**Table S1 |** Parameters, omnibus interaction tests, land-use-specific slopes and random effects for the four local-biodiversity models.

**Table S1a RQ1 fixed effects**

**Total abundance, mean-temperature anomaly**

| Predictor | Estimate | s.e. | 95% CI | z | P |
|---|---|---|---|---|---|
| Intercept: primary vegetation at anomaly = 0 | −0.854 | 0.0750 | [−1.001, −0.707] | −11.4 | 4.8e-30 |
| Secondary vegetation | −0.146 | 0.0344 | [−0.213, −0.079] | −4.24 | 2.2e-05 |
| Low-intensity agriculture | −0.125 | 0.0349 | [−0.194, −0.057] | −3.59 | 3.4e-04 |
| High-intensity agriculture | −0.175 | 0.0382 | [−0.250, −0.100] | −4.59 | 4.5e-06 |
| Urban | 0.277 | 0.0745 | [0.131, 0.423] | 3.71 | 2.0e-04 |
| Standardised mean-temperature anomaly | 0.0489 | 0.0563 | [−0.061, 0.159] | 0.868 | 0.385 |
| Secondary vegetation × mean-temperature anomaly | 0.0611 | 0.0469 | [−0.031, 0.153] | 1.30 | 0.193 |
| Low-intensity agriculture × mean-temperature anomaly | 0.128 | 0.0346 | [0.060, 0.196] | 3.69 | 2.2e-04 |
| High-intensity agriculture × mean-temperature anomaly | −0.00683 | 0.0347 | [−0.075, 0.061] | −0.197 | 0.844 |
| Urban × mean-temperature anomaly | 0.0922 | 0.114 | [−0.132, 0.316] | 0.807 | 0.420 |

**Total abundance, maximum-temperature anomaly**

| Predictor | Estimate | s.e. | 95% CI | z | P |
|---|---|---|---|---|---|
| Intercept: primary vegetation at anomaly = 0 | −0.845 | 0.0751 | [−0.992, −0.698] | −11.2 | 2.3e-29 |
| Secondary vegetation | −0.147 | 0.0338 | [−0.213, −0.080] | −4.34 | 1.4e-05 |
| Low-intensity agriculture | −0.107 | 0.0351 | [−0.176, −0.038] | −3.05 | 0.002 |
| High-intensity agriculture | −0.189 | 0.0376 | [−0.263, −0.116] | −5.03 | 4.8e-07 |
| Urban | 0.338 | 0.0620 | [0.216, 0.460] | 5.45 | 5.0e-08 |
| Standardised maximum-temperature anomaly | −0.112 | 0.0342 | [−0.179, −0.045] | −3.28 | 0.001 |
| Secondary vegetation × maximum-temperature anomaly | 0.0464 | 0.0338 | [−0.020, 0.113] | 1.37 | 0.170 |
| Low-intensity agriculture × maximum-temperature anomaly | 0.104 | 0.0324 | [0.041, 0.168] | 3.22 | 0.001 |
| High-intensity agriculture × maximum-temperature anomaly | 0.00739 | 0.0314 | [−0.054, 0.069] | 0.236 | 0.814 |
| Urban × maximum-temperature anomaly | 0.408 | 0.127 | [0.158, 0.657] | 3.21 | 0.001 |

**Species richness, mean-temperature anomaly**

| Predictor | Estimate | s.e. | 95% CI | z | P |
|---|---|---|---|---|---|
| Intercept: primary vegetation at anomaly = 0 | 2.81 | 0.0915 | [2.636, 2.994] | 30.8 | 7.0e-208 |
| Secondary vegetation | −0.104 | 0.0171 | [−0.138, −0.071] | −6.12 | 9.6e-10 |
| Low-intensity agriculture | −0.193 | 0.0183 | [−0.229, −0.157] | −10.5 | 5.3e-26 |
| High-intensity agriculture | −0.208 | 0.0200 | [−0.247, −0.169] | −10.4 | 2.3e-25 |
| Urban | −0.160 | 0.0417 | [−0.241, −0.078] | −3.83 | 1.3e-04 |
| Standardised mean-temperature anomaly | 0.0741 | 0.0496 | [−0.023, 0.171] | 1.49 | 0.135 |
| Secondary vegetation × mean-temperature anomaly | −0.00763 | 0.0206 | [−0.048, 0.033] | −0.370 | 0.711 |
| Low-intensity agriculture × mean-temperature anomaly | 0.0681 | 0.0169 | [0.035, 0.101] | 4.03 | 5.5e-05 |
| High-intensity agriculture × mean-temperature anomaly | −0.0630 | 0.0158 | [−0.094, −0.032] | −3.99 | 6.6e-05 |
| Urban × mean-temperature anomaly | −0.160 | 0.0631 | [−0.283, −0.036] | −2.53 | 0.011 |

**Species richness, maximum-temperature anomaly**

| Predictor | Estimate | s.e. | 95% CI | z | P |
|---|---|---|---|---|---|
| Intercept: primary vegetation at anomaly = 0 | 2.82 | 0.0928 | [2.637, 3.000] | 30.4 | 1.0e-202 |
| Secondary vegetation | −0.100 | 0.0174 | [−0.134, −0.066] | −5.75 | 8.7e-09 |
| Low-intensity agriculture | −0.183 | 0.0183 | [−0.219, −0.147] | −9.99 | 1.7e-23 |
| High-intensity agriculture | −0.227 | 0.0196 | [−0.266, −0.189] | −11.6 | 4.2e-31 |
| Urban | −0.0590 | 0.0339 | [−0.126, 0.007] | −1.74 | 0.082 |
| Standardised maximum-temperature anomaly | −0.0984 | 0.0209 | [−0.139, −0.057] | −4.71 | 2.5e-06 |
| Secondary vegetation × maximum-temperature anomaly | 0.0170 | 0.0149 | [−0.012, 0.046] | 1.14 | 0.255 |
| Low-intensity agriculture × maximum-temperature anomaly | 0.0740 | 0.0150 | [0.045, 0.103] | 4.92 | 8.7e-07 |
| High-intensity agriculture × maximum-temperature anomaly | −0.0288 | 0.0172 | [−0.063, 0.005] | −1.67 | 0.094 |

| Predictor | Estimate | s.e. | 95% CI | z | P |
|---|---|---|---|---|---|
| Urban × maximum-temperature anomaly | 0.136 | 0.0642 | [0.010, 0.262] | 2.12 | 0.034 |

**Table S1b RQ1 omnibus LRT**

| Model | n | $\chi^2$ | d.f. | P |
|---|---|---|---|---|
| Total abundance, mean-temperature anomaly | 5817. | 20.2 | 4.00 | 4.5e-04 |
| Total abundance, maximum-temperature anomaly | 5817. | 19.7 | 4.00 | 5.8e-04 |
| Species richness, mean-temperature anomaly | 6457. | 60.2 | 4.00 | 2.6e-12 |
| Species richness, maximum-temperature anomaly | 6457. | 33.1 | 4.00 | 1.1e-06 |

**Table S1c RQ1 landuse specific slopes**

| Response | Warming metric | Land use | Estimate | s.e. | 95% CI | P |
|---|---|---|---|---|---|---|
| Total abundance | Mean-temperature anomaly | Primary vegetation | 0.0489 | 0.0563 | [−0.061, 0.159] | 0.385 |
| Total abundance | Mean-temperature anomaly | Secondary vegetation | 0.110 | 0.0639 | [−0.015, 0.235] | 0.085 |
| Total abundance | Mean-temperature anomaly | Low-intensity agriculture | 0.177 | 0.0582 | [0.063, 0.291] | 0.002 |
| Total abundance | Mean-temperature anomaly | High-intensity agriculture | 0.0421 | 0.0580 | [−0.072, 0.156] | 0.468 |
| Total abundance | Mean-temperature anomaly | Urban | 0.141 | 0.124 | [−0.101, 0.384] | 0.254 |
| Total abundance | Maximum-temperature anomaly | Primary vegetation | −0.112 | 0.0342 | [−0.179, −0.045] | 0.001 |
| Total abundance | Maximum-temperature anomaly | Secondary vegetation | −0.0658 | 0.0424 | [−0.149, 0.017] | 0.121 |
| Total abundance | Maximum-temperature anomaly | Low-intensity agriculture | −0.00782 | 0.0430 | [−0.092, 0.076] | 0.856 |
| Total abundance | Maximum-temperature anomaly | High-intensity agriculture | −0.105 | 0.0425 | [−0.188, −0.021] | 0.014 |
| Total abundance | Maximum-temperature anomaly | Urban | 0.295 | 0.131 | [0.039, 0.551] | 0.024 |
| Species richness | Mean-temperature anomaly | Primary vegetation | 0.0741 | 0.0496 | [−0.023, 0.171] | 0.135 |
| Species richness | Mean-temperature anomaly | Secondary vegetation | 0.0665 | 0.0522 | [−0.036, 0.169] | 0.203 |
| Species richness | Mean-temperature anomaly | Low-intensity agriculture | 0.142 | 0.0507 | [0.043, 0.241] | 0.005 |
| Species richness | Mean-temperature anomaly | High-intensity agriculture | 0.0111 | 0.0505 | [−0.088, 0.110] | 0.826 |
| Species richness | Mean-temperature anomaly | Urban | −0.0855 | 0.0797 | [−0.242, 0.071] | 0.283 |
| Species richness | Maximum-temperature anomaly | Primary vegetation | −0.0984 | 0.0209 | [−0.139, −0.057] | 2.5e-06 |
| Species richness | Maximum-temperature anomaly | Secondary vegetation | −0.0814 | 0.0239 | [−0.128, −0.035] | 6.5e-04 |
| Species richness | Maximum-temperature anomaly | Low-intensity agriculture | −0.0244 | 0.0244 | [−0.072, 0.023] | 0.317 |
| Species richness | Maximum-temperature anomaly | High-intensity agriculture | −0.127 | 0.0257 | [−0.178, −0.077] | 7.4e-07 |
| Species richness | Maximum-temperature anomaly | Urban | 0.0375 | 0.0668 | [−0.093, 0.168] | 0.574 |

**Table S1d RQ1 random effects and fit**

**Total abundance, mean-temperature anomaly**

| Random effect | Variance | s.d. | Levels | Observations | AIC |
|---|---|---|---|---|---|
| Study | 0.555 | 0.745 | 121. | 5817. | 12301. |
| Spatial block within study | 0.0637 | 0.252 | 503. | 5817. | 12301. |
| Residual | 0.434 | 0.659 | | 5817. | 12301. |

**Total abundance, maximum-temperature anomaly**

| Random effect | Variance | s.d. | Levels | Observations | AIC |
|---|---|---|---|---|---|
| Study | 0.561 | 0.749 | 121. | 5817. | 12297. |
| Spatial block within study | 0.0668 | 0.259 | 503. | 5817. | 12297. |
| Residual | 0.433 | 0.658 | | 5817. | 12297. |

**Species richness, mean-temperature anomaly**

| Random effect | Variance | s.d. | Levels | Observations | AIC |
|---|---|---|---|---|---|
| Study | 1.09 | 1.04 | 138. | 6457. | 35510. |
| Spatial block within study | 0.0454 | 0.213 | 599. | 6457. | 35510. |
| Negative-binomial dispersion | | 28.9 | | 6457. | 35510. |

**Species richness, maximum-temperature anomaly**

| Random effect | Variance | s.d. | Levels | Observations | AIC |
|---|---|---|---|---|---|
| Study | 1.13 | 1.06 | 138. | 6457. | 35520. |
| Spatial block within study | 0.0483 | 0.220 | 599. | 6457. | 35520. |
| Negative-binomial dispersion | | 29.1 | | 6457. | 35520. |

**Table S2 |** Full ordered-beta model output for the nine assemblage-dissimilarity responses.

**Table S2a RQ2 ordered beta fixed effects**

**Bray-Curtis total dissimilarity**

| Predictor | Estimate | s.e. | 95% CI | z | P |
|---|---|---|---|---|---|
| Intercept: primary vegetation at anomaly = 0 | 0.419 | 0.154 | [0.118, 0.720] | 2.73 | 0.006 |
| Secondary vegetation | 0.111 | 0.0704 | [−0.026, 0.249] | 1.58 | 0.113 |
| Low-intensity agriculture | 0.0781 | 0.0660 | [−0.051, 0.208] | 1.18 | 0.237 |
| High-intensity agriculture | 0.139 | 0.0732 | [−0.005, 0.282] | 1.90 | 0.058 |
| Urban | −0.338 | 0.188 | [−0.707, 0.031] | −1.79 | 0.073 |
| Pair-level mean standardised temperature anomaly | −0.158 | 0.0919 | [−0.338, 0.022] | −1.72 | 0.086 |
| Absolute within-pair anomaly difference | −0.0487 | 0.0225 | [−0.093, −0.005] | −2.16 | 0.031 |
| Log geographic distance | 0.124 | 0.00266 | [0.119, 0.129] | 46.6 | 0.0e+00 |
| Secondary vegetation × pair-level mean anomaly | 0.107 | 0.0991 | [−0.088, 0.301] | 1.08 | 0.282 |
| Low-intensity agriculture × pair-level mean anomaly | −0.0380 | 0.0627 | [−0.161, 0.085] | −0.606 | 0.545 |
| High-intensity agriculture × pair-level mean anomaly | −0.00454 | 0.0636 | [−0.129, 0.120] | −0.0714 | 0.943 |
| Urban × pair-level mean anomaly | 0.0880 | 0.241 | [−0.383, 0.559] | 0.366 | 0.714 |

**Bray-Curtis balanced variation**

| Predictor | Estimate | s.e. | 95% CI | z | P |
|---|---|---|---|---|---|
| Intercept: primary vegetation at anomaly = 0 | −0.825 | 0.209 | [−1.234, −0.415] | −3.95 | 7.9e-05 |
| Secondary vegetation | 0.0743 | 0.0672 | [−0.058, 0.206] | 1.10 | 0.269 |
| Low-intensity agriculture | 0.0677 | 0.0621 | [−0.054, 0.189] | 1.09 | 0.275 |
| High-intensity agriculture | 0.108 | 0.0691 | [−0.027, 0.244] | 1.57 | 0.116 |
| Urban | −0.395 | 0.178 | [−0.744, −0.045] | −2.21 | 0.027 |
| Pair-level mean standardised temperature anomaly | −0.00101 | 0.0980 | [−0.193, 0.191] | −0.0103 | 0.992 |
| Absolute within-pair anomaly difference | −0.00114 | 0.0246 | [−0.049, 0.047] | −0.0461 | 0.963 |
| Log geographic distance | 0.125 | 0.00311 | [0.118, 0.131] | 40.0 | 0.0e+00 |
| Secondary vegetation × pair-level mean anomaly | 0.240 | 0.0961 | [0.051, 0.428] | 2.49 | 0.013 |
| Low-intensity agriculture × pair-level mean anomaly | 0.0258 | 0.0590 | [−0.090, 0.141] | 0.438 | 0.662 |
| High-intensity agriculture × pair-level mean anomaly | 0.0520 | 0.0602 | [−0.066, 0.170] | 0.864 | 0.388 |
| Urban × pair-level mean anomaly | 0.0373 | 0.227 | [−0.408, 0.483] | 0.164 | 0.870 |

**Bray-Curtis abundance-gradient dissimilarity**

| Predictor | Estimate | s.e. | 95% CI | z | P |
|---|---|---|---|---|---|
| Intercept: primary vegetation at anomaly = 0 | −1.74 | 0.197 | [−2.123, −1.352] | −8.83 | 1.1e-18 |
| Secondary vegetation | −0.0207 | 0.0932 | [−0.203, 0.162] | −0.222 | 0.824 |
| Low-intensity agriculture | 0.0777 | 0.0866 | [−0.092, 0.247] | 0.897 | 0.370 |
| High-intensity agriculture | 0.0311 | 0.0977 | [−0.160, 0.223] | 0.318 | 0.750 |
| Urban | 0.293 | 0.253 | [−0.204, 0.789] | 1.16 | 0.248 |
| Pair-level mean standardised temperature anomaly | −0.236 | 0.121 | [−0.474, 0.001] | −1.95 | 0.051 |
| Absolute within-pair anomaly difference | −0.116 | 0.0303 | [−0.175, −0.057] | −3.82 | 1.3e-04 |
| Log geographic distance | −0.0155 | 0.00375 | [−0.023, −0.008] | −4.13 | 3.6e-05 |
| Secondary vegetation × pair-level mean anomaly | −0.0750 | 0.132 | [−0.334, 0.184] | −0.567 | 0.571 |
| Low-intensity agriculture × pair-level mean anomaly | −0.109 | 0.0836 | [−0.273, 0.055] | −1.30 | 0.193 |
| High-intensity agriculture × pair-level mean anomaly | −0.0718 | 0.0852 | [−0.239, 0.095] | −0.842 | 0.400 |
| Urban × pair-level mean anomaly | 0.112 | 0.324 | [−0.522, 0.747] | 0.347 | 0.728 |

**Jaccard total dissimilarity**

| Predictor | Estimate | s.e. | 95% CI | z | P |
|---|---|---|---|---|---|
| Intercept: primary vegetation at anomaly = 0 | 0.583 | 0.165 | [0.259, 0.906] | 3.53 | 4.2e-04 |
| Secondary vegetation | 0.168 | 0.0797 | [0.012, 0.324] | 2.11 | 0.035 |
| Low-intensity agriculture | 0.0856 | 0.0747 | [−0.061, 0.232] | 1.15 | 0.252 |
| High-intensity agriculture | 0.133 | 0.0828 | [−0.030, 0.295] | 1.60 | 0.109 |
| Urban | −0.424 | 0.212 | [−0.840, −0.009] | −2.00 | 0.045 |

| Predictor | Estimate | s.e. | 95% CI | z | P |
|---|---|---|---|---|---|
| Pair-level mean standardised temperature anomaly | −0.141 | 0.103 | [−0.344, 0.062] | −1.36 | 0.172 |
| Absolute within-pair anomaly difference | −0.0404 | 0.0220 | [−0.084, 0.003] | −1.84 | 0.067 |
| Log geographic distance | 0.113 | 0.00244 | [0.109, 0.118] | 46.5 | 0.0e+00 |
| Secondary vegetation × pair-level mean anomaly | 0.104 | 0.112 | [−0.115, 0.324] | 0.931 | 0.352 |
| Low-intensity agriculture × pair-level mean anomaly | −0.00843 | 0.0709 | [−0.147, 0.130] | −0.119 | 0.905 |
| High-intensity agriculture × pair-level mean anomaly | −0.0199 | 0.0718 | [−0.161, 0.121] | −0.277 | 0.782 |
| Urban × pair-level mean anomaly | 0.186 | 0.271 | [−0.346, 0.717] | 0.685 | 0.494 |

**Jaccard turnover**

| Predictor | Estimate | s.e. | 95% CI | z | P |
|---|---|---|---|---|---|
| Intercept: primary vegetation at anomaly = 0 | −0.513 | 0.198 | [−0.901, −0.125] | −2.59 | 0.010 |
| Secondary vegetation | 0.0970 | 0.0621 | [−0.025, 0.219] | 1.56 | 0.118 |
| Low-intensity agriculture | 0.0657 | 0.0575 | [−0.047, 0.178] | 1.14 | 0.253 |
| High-intensity agriculture | 0.0974 | 0.0640 | [−0.028, 0.223] | 1.52 | 0.128 |
| Urban | −0.633 | 0.165 | [−0.956, −0.311] | −3.85 | 1.2e-04 |
| Pair-level mean standardised temperature anomaly | 0.0941 | 0.0910 | [−0.084, 0.273] | 1.03 | 0.301 |
| Absolute within-pair anomaly difference | −0.0230 | 0.0227 | [−0.068, 0.022] | −1.01 | 0.312 |
| Log geographic distance | 0.116 | 0.00277 | [0.110, 0.121] | 41.8 | 0.0e+00 |
| Secondary vegetation × pair-level mean anomaly | 0.142 | 0.0883 | [−0.031, 0.315] | 1.61 | 0.108 |
| Low-intensity agriculture × pair-level mean anomaly | −0.0120 | 0.0543 | [−0.118, 0.095] | −0.220 | 0.826 |
| High-intensity agriculture × pair-level mean anomaly | −0.0498 | 0.0555 | [−0.159, 0.059] | −0.897 | 0.370 |
| Urban × pair-level mean anomaly | −0.0755 | 0.210 | [−0.486, 0.335] | −0.360 | 0.719 |

**Jaccard nestedness-resultant dissimilarity**

| Predictor | Estimate | s.e. | 95% CI | z | P |
|---|---|---|---|---|---|
| Intercept: primary vegetation at anomaly = 0 | −1.82 | 0.205 | [−2.225, −1.419] | −8.87 | 7.4e-19 |
| Secondary vegetation | 0.0590 | 0.103 | [−0.143, 0.261] | 0.573 | 0.567 |
| Low-intensity agriculture | 0.198 | 0.0966 | [0.009, 0.388] | 2.05 | 0.040 |
| High-intensity agriculture | 0.144 | 0.107 | [−0.066, 0.355] | 1.35 | 0.178 |
| Urban | 0.570 | 0.273 | [0.034, 1.106] | 2.08 | 0.037 |
| Pair-level mean standardised temperature anomaly | −0.343 | 0.131 | [−0.601, −0.085] | −2.61 | 0.009 |
| Absolute within-pair anomaly difference | −0.0344 | 0.0299 | [−0.093, 0.024] | −1.15 | 0.249 |
| Log geographic distance | −0.0367 | 0.00366 | [−0.044, −0.030] | −10.0 | 1.1e-23 |
| Secondary vegetation × pair-level mean anomaly | 0.0174 | 0.145 | [−0.266, 0.301] | 0.120 | 0.904 |
| Low-intensity agriculture × pair-level mean anomaly | −0.102 | 0.0919 | [−0.282, 0.078] | −1.11 | 0.268 |
| High-intensity agriculture × pair-level mean anomaly | 0.00667 | 0.0931 | [−0.176, 0.189] | 0.0716 | 0.943 |
| Urban × pair-level mean anomaly | 0.506 | 0.350 | [−0.180, 1.191] | 1.45 | 0.148 |

**Sørensen total dissimilarity**

| Predictor | Estimate | s.e. | 95% CI | z | P |
|---|---|---|---|---|---|
| Intercept: primary vegetation at anomaly = 0 | −0.0960 | 0.164 | [−0.418, 0.226] | −0.584 | 0.559 |
| Secondary vegetation | 0.160 | 0.0788 | [0.006, 0.315] | 2.03 | 0.042 |
| Low-intensity agriculture | 0.0767 | 0.0737 | [−0.068, 0.221] | 1.04 | 0.298 |
| High-intensity agriculture | 0.133 | 0.0819 | [−0.027, 0.294] | 1.63 | 0.104 |
| Urban | −0.435 | 0.210 | [−0.847, −0.024] | −2.08 | 0.038 |
| Pair-level mean standardised temperature anomaly | −0.146 | 0.102 | [−0.347, 0.054] | −1.43 | 0.153 |
| Absolute within-pair anomaly difference | −0.0445 | 0.0207 | [−0.085, −0.004] | −2.15 | 0.032 |
| Log geographic distance | 0.119 | 0.00251 | [0.114, 0.123] | 47.3 | 0.0e+00 |
| Secondary vegetation × pair-level mean anomaly | 0.109 | 0.111 | [−0.108, 0.327] | 0.983 | 0.325 |
| Low-intensity agriculture × pair-level mean anomaly | −0.00558 | 0.0701 | [−0.143, 0.132] | −0.0796 | 0.937 |
| High-intensity agriculture × pair-level mean anomaly | −0.0193 | 0.0711 | [−0.159, 0.120] | −0.272 | 0.786 |
| Urban × pair-level mean anomaly | 0.179 | 0.268 | [−0.347, 0.704] | 0.667 | 0.505 |

**Sørensen turnover**

| Predictor | Estimate | s.e. | 95% CI | z | P |
|---|---|---|---|---|---|
| Intercept: primary vegetation at anomaly = 0 | −1.18 | 0.198 | [−1.565, −0.790] | −5.95 | 2.6e-09 |
| Secondary vegetation | 0.0933 | 0.0616 | [−0.027, 0.214] | 1.52 | 0.130 |
| Low-intensity agriculture | 0.0569 | 0.0569 | [−0.055, 0.168] | 1.00 | 0.318 |
| High-intensity agriculture | 0.0960 | 0.0635 | [−0.029, 0.221] | 1.51 | 0.131 |
| Urban | −0.642 | 0.164 | [−0.964, −0.320] | −3.91 | 9.2e-05 |
| Pair-level mean standardised temperature anomaly | 0.0832 | 0.0901 | [−0.093, 0.260] | 0.923 | 0.356 |
| Absolute within-pair anomaly difference | −0.0323 | 0.0214 | [−0.074, 0.010] | −1.51 | 0.131 |
| Log geographic distance | 0.122 | 0.00282 | [0.116, 0.127] | 43.2 | 0.0e+00 |
| Secondary vegetation × pair-level mean anomaly | 0.145 | 0.0878 | [−0.027, 0.317] | 1.65 | 0.099 |
| Low-intensity agriculture × pair-level mean anomaly | −0.0111 | 0.0539 | [−0.117, 0.095] | −0.205 | 0.838 |
| High-intensity agriculture × pair-level mean anomaly | −0.0482 | 0.0552 | [−0.156, 0.060] | −0.874 | 0.382 |
| Urban × pair-level mean anomaly | −0.0871 | 0.209 | [−0.497, 0.323] | −0.416 | 0.677 |

**Sørensen nestedness-resultant dissimilarity**

| Predictor | Estimate | s.e. | 95% CI | z | P |
|---|---|---|---|---|---|
| Intercept: primary vegetation at anomaly = 0 | −1.96 | 0.195 | [−2.343, −1.579] | −10.1 | 8.8e-24 |
| Secondary vegetation | 0.0498 | 0.0969 | [−0.140, 0.240] | 0.515 | 0.607 |
| Low-intensity agriculture | 0.195 | 0.0908 | [0.017, 0.373] | 2.15 | 0.032 |
| High-intensity agriculture | 0.146 | 0.101 | [−0.052, 0.343] | 1.44 | 0.149 |
| Urban | 0.487 | 0.257 | [−0.017, 0.990] | 1.90 | 0.058 |
| Pair-level mean standardised temperature anomaly | −0.321 | 0.124 | [−0.564, −0.077] | −2.58 | 0.010 |
| Absolute within-pair anomaly difference | −0.0365 | 0.0271 | [−0.090, 0.017] | −1.35 | 0.178 |
| Log geographic distance | −0.0149 | 0.00332 | [−0.021, −0.008] | −4.50 | 6.8e-06 |
| Secondary vegetation × pair-level mean anomaly | 0.0326 | 0.136 | [−0.234, 0.300] | 0.239 | 0.811 |
| Low-intensity agriculture × pair-level mean anomaly | −0.0938 | 0.0864 | [−0.263, 0.076] | −1.08 | 0.278 |
| High-intensity agriculture × pair-level mean anomaly | 0.00667 | 0.0876 | [−0.165, 0.178] | 0.0762 | 0.939 |
| Urban × pair-level mean anomaly | 0.526 | 0.328 | [−0.117, 1.169] | 1.60 | 0.109 |

**Table S2b RQ2 omnibus interaction LRT**

| Model | $\chi^2$ | d.f. | P |
|---|---|---|---|
| Bray-Curtis total dissimilarity | 2.26 | 4.00 | 0.687 |
| Bray-Curtis balanced variation | 6.53 | 4.00 | 0.163 |
| Bray-Curtis abundance-gradient dissimilarity | 1.94 | 4.00 | 0.747 |
| Jaccard total dissimilarity | 1.61 | 4.00 | 0.807 |
| Jaccard turnover | 4.71 | 4.00 | 0.318 |
| Jaccard nestedness-resultant dissimilarity | 4.49 | 4.00 | 0.344 |
| Sørensen total dissimilarity | 1.69 | 4.00 | 0.793 |
| Sørensen turnover | 4.87 | 4.00 | 0.301 |
| Sørensen nestedness-resultant dissimilarity | 5.09 | 4.00 | 0.278 |

**Table S2c RQ2 landuse specific warming slopes**

**Bray-Curtis total dissimilarity**

| Model family | Estimate | s.e. | 95% CI | P |
|---|---|---|---|---|
| ordered_beta | −0.158 | 0.0919 | [−0.338, 0.022] | 0.086 |
| ordered_beta | −0.0514 | 0.119 | [−0.284, 0.181] | 0.665 |
| ordered_beta | −0.196 | 0.0926 | [−0.377, −0.014] | 0.034 |
| ordered_beta | −0.163 | 0.0934 | [−0.346, 0.021] | 0.082 |
| ordered_beta | −0.0699 | 0.253 | [−0.567, 0.427] | 0.783 |
| gaussian_logit | −0.212 | 0.167 | [−0.538, 0.115] | 0.205 |
| gaussian_logit | −0.154 | 0.219 | [−0.584, 0.275] | 0.481 |
| gaussian_logit | −0.378 | 0.168 | [−0.706, −0.049] | 0.024 |
| gaussian_logit | −0.240 | 0.168 | [−0.569, 0.089] | 0.153 |
| gaussian_logit | 0.121 | 0.509 | [−0.877, 1.119] | 0.813 |

**Bray-Curtis balanced variation**

| Model family | Estimate | s.e. | 95% CI | P |
|---|---|---|---|---|
| ordered_beta | −0.00101 | 0.0980 | [−0.193, 0.191] | 0.992 |
| ordered_beta | 0.239 | 0.125 | [−0.007, 0.484] | 0.057 |
| ordered_beta | 0.0248 | 0.0978 | [−0.167, 0.216] | 0.800 |
| ordered_beta | 0.0510 | 0.0997 | [−0.144, 0.246] | 0.609 |
| ordered_beta | 0.0363 | 0.244 | [−0.441, 0.514] | 0.882 |
| gaussian_logit | 0.155 | 0.179 | [−0.195, 0.505] | 0.385 |
| gaussian_logit | 0.435 | 0.233 | [−0.022, 0.893] | 0.062 |
| gaussian_logit | 0.192 | 0.179 | [−0.159, 0.544] | 0.284 |
| gaussian_logit | 0.200 | 0.182 | [−0.156, 0.557] | 0.271 |
| gaussian_logit | 0.176 | 0.521 | [−0.844, 1.197] | 0.735 |

**Bray-Curtis abundance-gradient dissimilarity**

| Model family | Estimate | s.e. | 95% CI | P |
|---|---|---|---|---|
| ordered_beta | −0.236 | 0.121 | [−0.474, 0.001] | 0.051 |
| ordered_beta | −0.311 | 0.157 | [−0.619, −0.004] | 0.047 |
| ordered_beta | −0.345 | 0.122 | [−0.584, −0.107] | 0.005 |
| ordered_beta | −0.308 | 0.123 | [−0.549, −0.067] | 0.012 |
| ordered_beta | −0.124 | 0.340 | [−0.790, 0.543] | 0.716 |
| gaussian_logit | −0.494 | 0.217 | [−0.918, −0.070] | 0.023 |
| gaussian_logit | −0.472 | 0.281 | [−1.023, 0.079] | 0.093 |
| gaussian_logit | −0.426 | 0.218 | [−0.853, 0.001] | 0.050 |
| gaussian_logit | −0.418 | 0.220 | [−0.849, 0.012] | 0.057 |
| gaussian_logit | −0.393 | 0.628 | [−1.625, 0.839] | 0.532 |

**Jaccard total dissimilarity**

| Model family | Estimate | s.e. | 95% CI | P |
|---|---|---|---|---|
| ordered_beta | −0.141 | 0.103 | [−0.344, 0.062] | 0.172 |
| ordered_beta | −0.0367 | 0.134 | [−0.299, 0.226] | 0.784 |
| ordered_beta | −0.150 | 0.104 | [−0.353, 0.054] | 0.150 |
| ordered_beta | −0.161 | 0.105 | [−0.366, 0.044] | 0.124 |
| ordered_beta | 0.0444 | 0.285 | [−0.515, 0.604] | 0.876 |
| gaussian_logit | −0.338 | 0.184 | [−0.699, 0.022] | 0.066 |
| gaussian_logit | −0.297 | 0.243 | [−0.773, 0.180] | 0.222 |
| gaussian_logit | −0.418 | 0.185 | [−0.781, −0.054] | 0.024 |
| gaussian_logit | −0.344 | 0.184 | [−0.705, 0.016] | 0.061 |
| gaussian_logit | −0.0351 | 0.582 | [−1.176, 1.106] | 0.952 |

**Jaccard turnover**

| Model family | Estimate | s.e. | 95% CI | P |
|---|---|---|---|---|
| ordered_beta | 0.0941 | 0.0910 | [−0.084, 0.273] | 0.301 |
| ordered_beta | 0.236 | 0.116 | [0.008, 0.464] | 0.042 |
| ordered_beta | 0.0821 | 0.0908 | [−0.096, 0.260] | 0.366 |
| ordered_beta | 0.0443 | 0.0926 | [−0.137, 0.226] | 0.632 |
| ordered_beta | 0.0186 | 0.225 | [−0.423, 0.460] | 0.934 |
| gaussian_logit | 0.289 | 0.189 | [−0.082, 0.661] | 0.127 |
| gaussian_logit | 0.509 | 0.247 | [0.024, 0.994] | 0.040 |
| gaussian_logit | 0.367 | 0.190 | [−0.006, 0.740] | 0.054 |
| gaussian_logit | 0.217 | 0.193 | [−0.161, 0.595] | 0.261 |
| gaussian_logit | −0.174 | 0.552 | [−1.256, 0.908] | 0.753 |

**Jaccard nestedness-resultant dissimilarity**

| Model family | Estimate | s.e. | 95% CI | P |
|---|---|---|---|---|
| ordered_beta | −0.343 | 0.131 | [−0.601, −0.085] | 0.009 |

| Model family | Estimate | s.e. | 95% CI | P |
| --- | --- | --- | --- | --- |
| ordered_beta | −0.325 | 0.171 | [−0.660, 0.009] | 0.057 |
| ordered_beta | −0.445 | 0.132 | [−0.704, −0.185] | 7.7e-04 |
| ordered_beta | −0.336 | 0.133 | [−0.597, −0.075] | 0.012 |
| ordered_beta | 0.163 | 0.367 | [−0.557, 0.883] | 0.658 |
| gaussian_logit | −0.657 | 0.237 | [−1.122, −0.192] | 0.006 |
| gaussian_logit | −0.580 | 0.310 | [−1.187, 0.027] | 0.061 |
| gaussian_logit | −0.518 | 0.239 | [−0.986, −0.051] | 0.030 |
| gaussian_logit | −0.486 | 0.240 | [−0.956, −0.017] | 0.042 |
| gaussian_logit | 0.0451 | 0.706 | [−1.340, 1.430] | 0.949 |

**Sørensen total dissimilarity**

| Model family | Estimate | s.e. | 95% CI | P |
| --- | --- | --- | --- | --- |
| ordered_beta | −0.146 | 0.102 | [−0.347, 0.054] | 0.153 |
| ordered_beta | −0.0371 | 0.133 | [−0.297, 0.223] | 0.779 |
| ordered_beta | −0.152 | 0.103 | [−0.353, 0.049] | 0.139 |
| ordered_beta | −0.166 | 0.103 | [−0.368, 0.037] | 0.109 |
| ordered_beta | 0.0324 | 0.282 | [−0.521, 0.586] | 0.909 |
| gaussian_logit | −0.330 | 0.191 | [−0.704, 0.044] | 0.084 |
| gaussian_logit | −0.301 | 0.252 | [−0.795, 0.192] | 0.231 |
| gaussian_logit | −0.427 | 0.192 | [−0.804, −0.050] | 0.026 |
| gaussian_logit | −0.347 | 0.191 | [−0.722, 0.028] | 0.069 |
| gaussian_logit | −0.00280 | 0.599 | [−1.177, 1.172] | 0.996 |

**Sørensen turnover**

| Model family | Estimate | s.e. | 95% CI | P |
| --- | --- | --- | --- | --- |
| ordered_beta | 0.0832 | 0.0901 | [−0.093, 0.260] | 0.356 |
| ordered_beta | 0.228 | 0.115 | [0.002, 0.454] | 0.048 |
| ordered_beta | 0.0722 | 0.0899 | [−0.104, 0.248] | 0.422 |
| ordered_beta | 0.0350 | 0.0917 | [−0.145, 0.215] | 0.703 |
| ordered_beta | −0.00386 | 0.224 | [−0.444, 0.436] | 0.986 |
| gaussian_logit | 0.298 | 0.187 | [−0.069, 0.665] | 0.111 |
| gaussian_logit | 0.499 | 0.245 | [0.019, 0.979] | 0.042 |
| gaussian_logit | 0.342 | 0.188 | [−0.026, 0.710] | 0.069 |
| gaussian_logit | 0.210 | 0.190 | [−0.163, 0.582] | 0.270 |
| gaussian_logit | −0.0952 | 0.552 | [−1.178, 0.988] | 0.863 |

**Sørensen nestedness-resultant dissimilarity**

| Model family | Estimate | s.e. | 95% CI | P |
| --- | --- | --- | --- | --- |
| ordered_beta | −0.321 | 0.124 | [−0.564, −0.077] | 0.010 |
| ordered_beta | −0.288 | 0.161 | [−0.604, 0.028] | 0.074 |
| ordered_beta | −0.414 | 0.125 | [−0.659, −0.170] | 9.0e-04 |
| ordered_beta | −0.314 | 0.126 | [−0.560, −0.068] | 0.012 |
| ordered_beta | 0.205 | 0.345 | [−0.471, 0.882] | 0.552 |
| gaussian_logit | −0.670 | 0.237 | [−1.134, −0.205] | 0.005 |
| gaussian_logit | −0.566 | 0.309 | [−1.172, 0.040] | 0.067 |
| gaussian_logit | −0.508 | 0.238 | [−0.975, −0.041] | 0.033 |
| gaussian_logit | −0.497 | 0.239 | [−0.965, −0.028] | 0.038 |
| gaussian_logit | 0.0223 | 0.706 | [−1.361, 1.406] | 0.975 |

**Table S2d RQ2 dispersion random effects and fit**

**Bray-Curtis total dissimilarity**

| Component | Predictor | Estimate | scale | Observations | Studies | AIC |
| --- | --- | --- | --- | --- | --- | --- |
| Random effect | SS | 1.80 | variance | 144728. | 88.0 | −118055. |
| Random effect | s1 | 0.671 | variance | 144728. | 88.0 | −118055. |

| Component | Predictor | Estimate | scale | Observations | Studies | AIC |
|---|---|---|---|---|---|---|
| Random effect | s2 | 0.655 | variance | 144728. | 88.0 | −118055. |
| Dispersion model | Intercept | 2.85 | log precision | 144728. | 88.0 | −118055. |

**Bray-Curtis balanced variation**

| Component | Predictor | Estimate | scale | Observations | Studies | AIC |
|---|---|---|---|---|---|---|
| Random effect | SS | 3.51 | variance | 144728. | 88.0 | −27393. |
| Random effect | s1 | 0.575 | variance | 144728. | 88.0 | −27393. |
| Random effect | s2 | 0.552 | variance | 144728. | 88.0 | −27393. |
| Dispersion model | Intercept | 2.42 | log precision | 144728. | 88.0 | −27393. |

**Bray-Curtis abundance-gradient dissimilarity**

| Component | Predictor | Estimate | scale | Observations | Studies | AIC |
|---|---|---|---|---|---|---|
| Random effect | SS | 2.93 | variance | 144728. | 88.0 | −198371. |
| Random effect | s1 | 1.17 | variance | 144728. | 88.0 | −198371. |
| Random effect | s2 | 1.19 | variance | 144728. | 88.0 | −198371. |
| Dispersion model | Intercept | 2.64 | log precision | 144728. | 88.0 | −198371. |

**Jaccard total dissimilarity**

| Component | Predictor | Estimate | scale | Observations | Studies | AIC |
|---|---|---|---|---|---|---|
| Random effect | SS | 2.05 | variance | 144728. | 88.0 | −149954. |
| Random effect | s1 | 0.856 | variance | 144728. | 88.0 | −149954. |
| Random effect | s2 | 0.873 | variance | 144728. | 88.0 | −149954. |
| Dispersion model | Intercept | 3.12 | log precision | 144728. | 88.0 | −149954. |

**Jaccard turnover**

| Component | Predictor | Estimate | scale | Observations | Studies | AIC |
|---|---|---|---|---|---|---|
| Random effect | SS | 3.14 | variance | 144728. | 88.0 | −51524. |
| Random effect | s1 | 0.475 | variance | 144728. | 88.0 | −51524. |
| Random effect | s2 | 0.488 | variance | 144728. | 88.0 | −51524. |
| Dispersion model | Intercept | 2.77 | log precision | 144728. | 88.0 | −51524. |

**Jaccard nestedness-resultant dissimilarity**

| Component | Predictor | Estimate | scale | Observations | Studies | AIC |
|---|---|---|---|---|---|---|
| Random effect | SS | 3.15 | variance | 144728. | 88.0 | −168045. |
| Random effect | s1 | 1.40 | variance | 144728. | 88.0 | −168045. |
| Random effect | s2 | 1.46 | variance | 144728. | 88.0 | −168045. |
| Dispersion model | Intercept | 2.87 | log precision | 144728. | 88.0 | −168045. |

**Sørensen total dissimilarity**

| Component | Predictor | Estimate | scale | Observations | Studies | AIC |
|---|---|---|---|---|---|---|
| Random effect | SS | 2.03 | variance | 144728. | 88.0 | −98044. |
| Random effect | s1 | 0.835 | variance | 144728. | 88.0 | −98044. |
| Random effect | s2 | 0.848 | variance | 144728. | 88.0 | −98044. |
| Dispersion model | Intercept | 2.91 | log precision | 144728. | 88.0 | −98044. |

**Sørensen turnover**

| Component | Predictor | Estimate | scale | Observations | Studies | AIC |
|---|---|---|---|---|---|---|
| Random effect | SS | 3.14 | variance | 144728. | 88.0 | −24690. |
| Random effect | s1 | 0.465 | variance | 144728. | 88.0 | −24690. |
| Random effect | s2 | 0.477 | variance | 144728. | 88.0 | −24690. |
| Dispersion model | Intercept | 2.65 | log precision | 144728. | 88.0 | −24690. |

**Sørensen nestedness-resultant dissimilarity**

| Component | Predictor | Estimate | scale | Observations | Studies | AIC |
|---|---|---|---|---|---|---|

| | | | | | | |
|---|---|---|---|---|---|---|
| Random effect | SS | 2.85 | variance | 144728. | 88.0 | −177698. |
| Random effect | s1 | 1.23 | variance | 144728. | 88.0 | −177698. |
| Random effect | s2 | 1.30 | variance | 144728. | 88.0 | −177698. |
| Dispersion model | Intercept | 3.15 | log precision | 144728. | 88.0 | −177698. |

**Table S2e RQ2 boundary and sample summary**

| Model | Pairs | Studies | Sites | Zeros | Ones | Interior | Boundary fraction |
|---|---|---|---|---|---|---|---|
| Bray-Curtis total dissimilarity | 144728. | 88.0 | 4809. | 814. | 18898. | 125016. | 0.136 |
| Bray-Curtis balanced variation | 144728. | 88.0 | 4809. | 11986. | 12473. | 120269. | 0.169 |
| Bray-Curtis abundance-gradient dissimilarity | 144728. | 88.0 | 4809. | 18077. | 6425. | 120226. | 0.169 |
| Jaccard total dissimilarity | 144728. | 88.0 | 4809. | 1769. | 18898. | 124061. | 0.143 |
| Jaccard turnover | 144728. | 88.0 | 4809. | 15892. | 12473. | 116363. | 0.196 |
| Jaccard nestedness-resultant dissimilarity | 144728. | 88.0 | 4809. | 26449. | 6425. | 111854. | 0.227 |
| Sørensen total dissimilarity | 144728. | 88.0 | 4809. | 1769. | 18898. | 124061. | 0.143 |
| Sørensen turnover | 144728. | 88.0 | 4809. | 15892. | 12473. | 116363. | 0.196 |
| Sørensen nestedness-resultant dissimilarity | 144728. | 88.0 | 4809. | 26449. | 6425. | 111854. | 0.227 |

**Table S2f RQ2 additive ordered beta fixed effects**

Reduced models from the likelihood-ratio tests, without the land-use-by-warming interaction. Because the omnibus interaction was not supported for any response, these models provide the common warming slope across land-use classes; the land-use-specific slopes reported in the main text and shown in Fig. 4 are from the full interaction models (Tables S2a,c). P (BH), Benjamini–Hochberg-adjusted across the nine responses separately for each predictor.

**Bray-Curtis total dissimilarity**

| Predictor | Estimate | s.e. | 95% CI | z | P | P (BH) |
|---|---|---|---|---|---|---|
| Intercept: primary vegetation at anomaly = 0 | 0.431 | 0.153 | [0.131, 0.731] | 2.81 | 0.005 | – |
| Secondary vegetation | 0.0753 | 0.0649 | [−0.052, 0.203] | 1.16 | 0.246 | 0.737 |
| Low-intensity agriculture | 0.0715 | 0.0650 | [−0.056, 0.199] | 1.10 | 0.272 | 0.322 |
| High-intensity agriculture | 0.138 | 0.0698 | [0.001, 0.275] | 1.98 | 0.048 | 0.207 |
| Urban | −0.397 | 0.116 | [−0.624, −0.169] | −3.41 | <0.001 | <0.001 |
| Pair-level mean standardised temperature anomaly | −0.157 | 0.0849 | [−0.324, 0.009] | −1.85 | 0.064 | 0.144 |
| Absolute within-pair anomaly difference | −0.0484 | 0.0225 | [−0.093, −0.004] | −2.15 | 0.032 | 0.096 |
| Log geographic distance | 0.124 | 0.00265 | [0.119, 0.129] | 46.6 | <0.001 | <0.001 |

**Bray-Curtis balanced variation**

| Predictor | Estimate | s.e. | 95% CI | z | P | P (BH) |
|---|---|---|---|---|---|---|
| Intercept: primary vegetation at anomaly = 0 | −0.803 | 0.208 | [−1.211, −0.395] | −3.86 | <0.001 | – |
| Secondary vegetation | 0.0123 | 0.0614 | [−0.108, 0.133] | 0.200 | 0.841 | 0.841 |
| Low-intensity agriculture | 0.0688 | 0.0611 | [−0.051, 0.189] | 1.13 | 0.260 | 0.322 |
| High-intensity agriculture | 0.111 | 0.0657 | [−0.018, 0.240] | 1.69 | 0.091 | 0.207 |
| Urban | −0.422 | 0.109 | [−0.635, −0.209] | −3.88 | <0.001 | <0.001 |
| Pair-level mean standardised temperature anomaly | 0.0359 | 0.0924 | [−0.145, 0.217] | 0.390 | 0.698 | 0.698 |
| Absolute within-pair anomaly difference | −0.000946 | 0.0246 | [−0.049, 0.047] | −0.0400 | 0.969 | 0.969 |
| Log geographic distance | 0.124 | 0.00311 | [0.118, 0.131] | 40.0 | <0.001 | <0.001 |

**Bray-Curtis abundance-gradient dissimilarity**

| Predictor | Estimate | s.e. | 95% CI | z | P | P (BH) |
|---|---|---|---|---|---|---|
| Intercept: primary vegetation at anomaly = 0 | −1.74 | 0.196 | [−2.122, −1.352] | −8.84 | <0.001 | – |
| Secondary vegetation | −0.0208 | 0.0890 | [−0.195, 0.154] | −0.234 | 0.815 | 0.841 |
| Low-intensity agriculture | 0.0623 | 0.0896 | [−0.113, 0.238] | 0.695 | 0.487 | 0.487 |
| High-intensity agriculture | 0.0257 | 0.0935 | [−0.157, 0.209] | 0.275 | 0.783 | 0.783 |
| Urban | 0.219 | 0.156 | [−0.086, 0.524] | 1.41 | 0.160 | 0.180 |
| Pair-level mean standardised temperature anomaly | −0.292 | 0.112 | [−0.511, −0.073] | −2.62 | 0.009 | 0.027 |
| Absolute within-pair anomaly difference | −0.116 | 0.0303 | [−0.175, −0.056] | −3.81 | <0.001 | 0.001 |
| Log geographic distance | −0.0155 | 0.00376 | [−0.023, −0.008] | −4.12 | <0.001 | <0.001 |

**Jaccard total dissimilarity**

| Predictor | Estimate | s.e. | 95% CI | z | P | P (BH) |
|---|---|---|---|---|---|---|
| Intercept: primary vegetation at anomaly = 0 | 0.594 | 0.165 | [0.272, 0.917] | 3.61 | <0.001 | – |
| Secondary vegetation | 0.132 | 0.0736 | [−0.012, 0.276] | 1.79 | 0.073 | 0.404 |
| Low-intensity agriculture | 0.0856 | 0.0736 | [−0.059, 0.230] | 1.16 | 0.245 | 0.322 |
| High-intensity agriculture | 0.117 | 0.0790 | [−0.038, 0.272] | 1.48 | 0.138 | 0.207 |
| Urban | −0.540 | 0.132 | [−0.798, −0.282] | −4.11 | <0.001 | <0.001 |
| Pair-level mean standardised temperature anomaly | −0.136 | 0.0952 | [−0.323, 0.050] | −1.43 | 0.152 | 0.227 |
| Absolute within-pair anomaly difference | −0.0403 | 0.0220 | [−0.083, 0.003] | −1.83 | 0.067 | 0.152 |
| Log geographic distance | 0.113 | 0.00244 | [0.109, 0.118] | 46.5 | <0.001 | <0.001 |

**Jaccard turnover**

| Predictor | Estimate | s.e. | 95% CI | z | P | P (BH) |
|---|---|---|---|---|---|---|
| Intercept: primary vegetation at anomaly = 0 | −0.500 | 0.197 | [−0.887, −0.113] | −2.53 | 0.011 | – |
| Secondary vegetation | 0.0492 | 0.0568 | [−0.062, 0.160] | 0.870 | 0.386 | 0.763 |
| Low-intensity agriculture | 0.0749 | 0.0565 | [−0.036, 0.186] | 1.33 | 0.185 | 0.322 |
| High-intensity agriculture | 0.0705 | 0.0608 | [−0.049, 0.190] | 1.16 | 0.246 | 0.280 |
| Urban | −0.589 | 0.100 | [−0.786, −0.393] | −5.87 | <0.001 | <0.001 |
| Pair-level mean standardised temperature anomaly | 0.0882 | 0.0859 | [−0.080, 0.257] | 1.03 | 0.305 | 0.392 |
| Absolute within-pair anomaly difference | −0.0229 | 0.0227 | [−0.067, 0.022] | −1.01 | 0.314 | 0.353 |
| Log geographic distance | 0.116 | 0.00277 | [0.110, 0.121] | 41.8 | <0.001 | <0.001 |

**Jaccard nestedness-resultant dissimilarity**

| Predictor | Estimate | s.e. | 95% CI | z | P | P (BH) |
|---|---|---|---|---|---|---|
| Intercept: primary vegetation at anomaly = 0 | −1.81 | 0.205 | [−2.211, −1.407] | −8.82 | <0.001 | – |
| Secondary vegetation | 0.0356 | 0.0950 | [−0.150, 0.222] | 0.375 | 0.707 | 0.841 |
| Low-intensity agriculture | 0.168 | 0.0953 | [−0.019, 0.355] | 1.76 | 0.078 | 0.322 |
| High-intensity agriculture | 0.160 | 0.102 | [−0.041, 0.360] | 1.56 | 0.119 | 0.207 |
| Urban | 0.253 | 0.170 | [−0.081, 0.586] | 1.49 | 0.137 | 0.177 |
| Pair-level mean standardised temperature anomaly | −0.365 | 0.121 | [−0.602, −0.128] | −3.02 | 0.003 | 0.013 |
| Absolute within-pair anomaly difference | −0.0340 | 0.0299 | [−0.093, 0.024] | −1.14 | 0.254 | 0.327 |
| Log geographic distance | −0.0367 | 0.00366 | [−0.044, −0.029] | −10.0 | <0.001 | <0.001 |

**Sørensen total dissimilarity**

| Predictor | Estimate | s.e. | 95% CI | z | P | P (BH) |
|---|---|---|---|---|---|---|
| Intercept: primary vegetation at anomaly = 0 | −0.0841 | 0.164 | [−0.405, 0.237] | −0.510 | 0.607 | – |
| Secondary vegetation | 0.123 | 0.0726 | [−0.019, 0.266] | 1.70 | 0.090 | 0.404 |
| Low-intensity agriculture | 0.0774 | 0.0727 | [−0.065, 0.220] | 1.07 | 0.286 | 0.322 |
| High-intensity agriculture | 0.117 | 0.0781 | [−0.036, 0.270] | 1.50 | 0.133 | 0.207 |
| Urban | −0.548 | 0.130 | [−0.803, −0.293] | −4.21 | <0.001 | <0.001 |
| Pair-level mean standardised temperature anomaly | −0.140 | 0.0942 | [−0.325, 0.044] | −1.49 | 0.136 | 0.227 |
| Absolute within-pair anomaly difference | −0.0444 | 0.0207 | [−0.085, −0.004] | −2.14 | 0.032 | 0.096 |
| Log geographic distance | 0.119 | 0.00251 | [0.114, 0.123] | 47.3 | <0.001 | <0.001 |

**Sørensen turnover**

| Predictor | Estimate | s.e. | 95% CI | z | P | P (BH) |
|---|---|---|---|---|---|---|
| Intercept: primary vegetation at anomaly = 0 | −1.16 | 0.197 | [−1.551, −0.778] | −5.90 | <0.001 | – |
| Secondary vegetation | 0.0449 | 0.0562 | [−0.065, 0.155] | 0.800 | 0.424 | 0.763 |
| Low-intensity agriculture | 0.0663 | 0.0559 | [−0.043, 0.176] | 1.18 | 0.236 | 0.322 |
| High-intensity agriculture | 0.0695 | 0.0603 | [−0.049, 0.188] | 1.15 | 0.249 | 0.280 |
| Urban | −0.591 | 0.0995 | [−0.786, −0.396] | −5.94 | <0.001 | <0.001 |
| Pair-level mean standardised temperature anomaly | 0.0780 | 0.0850 | [−0.089, 0.245] | 0.920 | 0.359 | 0.404 |
| Absolute within-pair anomaly difference | −0.0322 | 0.0214 | [−0.074, 0.010] | −1.50 | 0.132 | 0.238 |
| Log geographic distance | 0.122 | 0.00282 | [0.116, 0.127] | 43.2 | <0.001 | <0.001 |

**Sørensen nestedness-resultant dissimilarity**

| Predictor | Estimate | s.e. | 95% CI | z | P | P (BH) |
|---|---|---|---|---|---|---|
| Intercept: primary vegetation at anomaly = 0 | −1.95 | 0.195 | [−2.329, −1.565] | −10.0 | <0.001 | – |
| Secondary vegetation | 0.0237 | 0.0897 | [−0.152, 0.200] | 0.264 | 0.792 | 0.841 |
| Low-intensity agriculture | 0.167 | 0.0896 | [−0.009, 0.342] | 1.86 | 0.063 | 0.322 |
| High-intensity agriculture | 0.159 | 0.0964 | [−0.030, 0.347] | 1.65 | 0.100 | 0.207 |
| Urban | 0.158 | 0.160 | [−0.155, 0.472] | 0.988 | 0.323 | 0.323 |

| Predictor | Estimate | s.e. | 95% CI | z | P | P (BH) |
|---|---|---|---|---|---|---|
| Pair-level mean standardised temperature anomaly | −0.339 | 0.114 | [−0.563, −0.116] | −2.97 | 0.003 | 0.013 |
| Absolute within-pair anomaly difference | −0.0361 | 0.0271 | [−0.089, 0.017] | −1.33 | 0.183 | 0.274 |
| Log geographic distance | −0.0149 | 0.00332 | [−0.021, −0.008] | −4.49 | <0.001 | <0.001 |

**Table S2g RQ2 warming-slope contrasts for secondary vegetation**

Differences in the warming–dissimilarity slope (logit scale) between secondary vegetation and each other land-use class, from the interaction models (Extended Data Fig. 3d). These are post hoc coefficient-level contrasts; the omnibus likelihood-ratio tests (Table S2b) remain the primary test of land-use dependence. P, unadjusted, as reported in the main text; P (BH), Benjamini–Hochberg-adjusted across all 36 contrasts.

**Bray-Curtis total dissimilarity**

| Contrast | Estimate | s.e. | 95% CI | z | P | P (BH) |
|---|---|---|---|---|---|---|
| Secondary vegetation vs primary vegetation | 0.107 | 0.0991 | [−0.088, 0.301] | 1.08 | 0.282 | 0.558 |
| Secondary vegetation vs low-intensity agriculture | 0.145 | 0.0999 | [−0.051, 0.340] | 1.45 | 0.148 | 0.496 |
| Secondary vegetation vs high-intensity agriculture | 0.111 | 0.101 | [−0.087, 0.310] | 1.10 | 0.273 | 0.558 |
| Secondary vegetation vs urban | 0.0185 | 0.252 | [−0.475, 0.512] | 0.0736 | 0.941 | 0.969 |

**Bray-Curtis balanced variation**

| Contrast | Estimate | s.e. | 95% CI | z | P | P (BH) |
|---|---|---|---|---|---|---|
| Secondary vegetation vs primary vegetation | 0.240 | 0.0961 | [0.051, 0.428] | 2.49 | 0.013 | 0.309 |
| Secondary vegetation vs low-intensity agriculture | 0.214 | 0.0967 | [0.024, 0.403] | 2.21 | 0.027 | 0.309 |
| Secondary vegetation vs high-intensity agriculture | 0.188 | 0.0984 | [−0.005, 0.380] | 1.91 | 0.057 | 0.407 |
| Secondary vegetation vs urban | 0.202 | 0.239 | [−0.265, 0.670] | 0.848 | 0.396 | 0.595 |

**Bray-Curtis abundance-gradient dissimilarity**

| Contrast | Estimate | s.e. | 95% CI | z | P | P (BH) |
|---|---|---|---|---|---|---|
| Secondary vegetation vs primary vegetation | −0.0750 | 0.132 | [−0.334, 0.184] | −0.567 | 0.571 | 0.773 |
| Secondary vegetation vs low-intensity agriculture | 0.0338 | 0.133 | [−0.226, 0.294] | 0.255 | 0.799 | 0.941 |
| Secondary vegetation vs high-intensity agriculture | −0.00325 | 0.135 | [−0.268, 0.262] | −0.0240 | 0.981 | 0.981 |
| Secondary vegetation vs urban | −0.187 | 0.339 | [−0.851, 0.476] | −0.554 | 0.580 | 0.773 |

**Jaccard total dissimilarity**

| Contrast | Estimate | s.e. | 95% CI | z | P | P (BH) |
|---|---|---|---|---|---|---|
| Secondary vegetation vs primary vegetation | 0.104 | 0.112 | [−0.115, 0.324] | 0.931 | 0.352 | 0.559 |
| Secondary vegetation vs low-intensity agriculture | 0.113 | 0.113 | [−0.108, 0.334] | 0.999 | 0.318 | 0.558 |
| Secondary vegetation vs high-intensity agriculture | 0.124 | 0.114 | [−0.100, 0.349] | 1.09 | 0.278 | 0.558 |
| Secondary vegetation vs urban | −0.0811 | 0.284 | [−0.638, 0.475] | −0.286 | 0.775 | 0.941 |

**Jaccard turnover**

| Contrast | Estimate | s.e. | 95% CI | z | P | P (BH) |
|---|---|---|---|---|---|---|
| Secondary vegetation vs primary vegetation | 0.142 | 0.0883 | [−0.031, 0.315] | 1.61 | 0.108 | 0.433 |
| Secondary vegetation vs low-intensity agriculture | 0.154 | 0.0889 | [−0.020, 0.328] | 1.73 | 0.084 | 0.430 |
| Secondary vegetation vs high-intensity agriculture | 0.192 | 0.0906 | [0.014, 0.369] | 2.12 | 0.034 | 0.309 |
| Secondary vegetation vs urban | 0.217 | 0.220 | [−0.214, 0.649] | 0.988 | 0.323 | 0.558 |

**Jaccard nestedness-resultant dissimilarity**

| Contrast | Estimate | s.e. | 95% CI | z | P | P (BH) |
|---|---|---|---|---|---|---|
| Secondary vegetation vs primary vegetation | 0.0174 | 0.145 | [−0.266, 0.301] | 0.120 | 0.904 | 0.969 |

| Contrast | Estimate | s.e. | 95% CI | z | P | P (BH) |
|---|---|---|---|---|---|---|
| Secondary vegetation vs low-intensity agriculture | 0.119 | 0.146 | [−0.166, 0.405] | 0.818 | 0.413 | 0.595 |
| Secondary vegetation vs high-intensity agriculture | 0.0107 | 0.148 | [−0.279, 0.300] | 0.0726 | 0.942 | 0.969 |
| Secondary vegetation vs urban | −0.488 | 0.366 | [−1.206, 0.230] | −1.33 | 0.182 | 0.547 |

**Sørensen total dissimilarity**

| Contrast | Estimate | s.e. | 95% CI | z | P | P (BH) |
|---|---|---|---|---|---|---|
| Secondary vegetation vs primary vegetation | 0.109 | 0.111 | [−0.108, 0.327] | 0.983 | 0.325 | 0.558 |
| Secondary vegetation vs low-intensity agriculture | 0.115 | 0.112 | [−0.104, 0.334] | 1.03 | 0.305 | 0.558 |
| Secondary vegetation vs high-intensity agriculture | 0.128 | 0.113 | [−0.094, 0.351] | 1.13 | 0.257 | 0.558 |
| Secondary vegetation vs urban | −0.0696 | 0.281 | [−0.620, 0.481] | −0.248 | 0.804 | 0.941 |

**Sørensen turnover**

| Contrast | Estimate | s.e. | 95% CI | z | P | P (BH) |
|---|---|---|---|---|---|---|
| Secondary vegetation vs primary vegetation | 0.145 | 0.0878 | [−0.027, 0.317] | 1.65 | 0.099 | 0.433 |
| Secondary vegetation vs low-intensity agriculture | 0.156 | 0.0884 | [−0.017, 0.329] | 1.76 | 0.078 | 0.430 |
| Secondary vegetation vs high-intensity agriculture | 0.193 | 0.0901 | [0.017, 0.370] | 2.14 | 0.032 | 0.309 |
| Secondary vegetation vs urban | 0.232 | 0.219 | [−0.198, 0.662] | 1.06 | 0.291 | 0.558 |

**Sørensen nestedness-resultant dissimilarity**

| Contrast | Estimate | s.e. | 95% CI | z | P | P (BH) |
|---|---|---|---|---|---|---|
| Secondary vegetation vs primary vegetation | 0.0326 | 0.136 | [−0.234, 0.300] | 0.239 | 0.811 | 0.941 |
| Secondary vegetation vs low-intensity agriculture | 0.126 | 0.137 | [−0.143, 0.395] | 0.921 | 0.357 | 0.559 |
| Secondary vegetation vs high-intensity agriculture | 0.0260 | 0.139 | [−0.247, 0.299] | 0.187 | 0.852 | 0.958 |
| Secondary vegetation vs urban | −0.493 | 0.344 | [−1.168, 0.181] | −1.43 | 0.151 | 0.496 |

**Table S3 |** Dependence and influence analyses for assemblage dissimilarity. Table S3a reports 1,000 strictly disjoint-pair ordered-beta refits addressing shared-site dependence. Tables S3b,c report 1,000 study-cluster bootstrap resamples and leave-one-study-out ranges from Gaussian mixed models of bounded logit-transformed dissimilarity, addressing study representation and influence as a separate distributional sensitivity analysis.

**Table S3a RQ2 disjoint pair subsampling**

| metric | n_success | median | Empirical 95% interval | main_estimate | Main-model 95% CI | direction_match | empirical_interval_excludes_zero |
|---|---|---|---|---|---|---|---|
| Bray-Curtis abundance gradient | 1000. | −0.0724 | [−0.139, −0.002] | −0.308 | [−0.549, −0.067] | TRUE | TRUE |
| Bray-Curtis abundance gradient | 1000. | −0.162 | [−0.237, −0.084] | −0.345 | [−0.584, −0.107] | TRUE | TRUE |
| Bray-Curtis abundance gradient | 1000. | −0.0432 | [−0.121, 0.027] | −0.236 | [−0.474, 0.001] | TRUE | FALSE |
| Bray-Curtis abundance gradient | 1000. | −0.0428 | [−0.152, 0.063] | −0.311 | [−0.619, −0.004] | TRUE | FALSE |
| Bray-Curtis abundance gradient | 1000. | 0.0632 | [−0.282, 0.367] | −0.124 | [−0.790, 0.543] | FALSE | FALSE |
| Bray-Curtis total dissimilarity | 1000. | −0.0767 | [−0.154, −0.015] | −0.163 | [−0.346, 0.021] | TRUE | TRUE |
| Bray-Curtis total dissimilarity | 1000. | −0.123 | [−0.196, −0.055] | −0.196 | [−0.377, −0.014] | TRUE | TRUE |

| metric | n_success | median | Empirical 95% interval | main_estimate | Main-model 95% CI | direction_match | empirical_interval_excludes_zero |
|---|---|---|---|---|---|---|---|
| Bray-Curtis total dissimilarity | 1000. | −0.116 | [−0.186, −0.045] | −0.158 | [−0.338, 0.022] | TRUE | TRUE |
| Bray-Curtis total dissimilarity | 1000. | 0.0319 | [−0.055, 0.116] | −0.0514 | [−0.284, 0.181] | FALSE | FALSE |
| Bray-Curtis total dissimilarity | 1000. | −0.0331 | [−0.295, 0.205] | −0.0699 | [−0.567, 0.427] | TRUE | FALSE |
| Bray-Curtis balanced variation | 1000. | 0.00750 | [−0.079, 0.093] | 0.0510 | [−0.144, 0.246] | TRUE | FALSE |
| Bray-Curtis balanced variation | 1000. | 0.0128 | [−0.080, 0.103] | 0.0248 | [−0.167, 0.216] | TRUE | FALSE |
| Bray-Curtis balanced variation | 1000. | −0.0526 | [−0.144, 0.038] | −0.00101 | [−0.193, 0.191] | TRUE | FALSE |
| Bray-Curtis balanced variation | 1000. | 0.0991 | [−0.019, 0.213] | 0.239 | [−0.007, 0.484] | TRUE | FALSE |
| Bray-Curtis balanced variation | 1000. | −0.0215 | [−0.285, 0.257] | 0.0363 | [−0.441, 0.514] | FALSE | FALSE |
| Jaccard nestedness-resultant | 1000. | −0.0733 | [−0.133, −0.014] | −0.336 | [−0.597, −0.075] | TRUE | TRUE |
| Jaccard nestedness-resultant | 1000. | −0.227 | [−0.304, −0.159] | −0.445 | [−0.704, −0.185] | TRUE | TRUE |
| Jaccard nestedness-resultant | 1000. | −0.103 | [−0.166, −0.042] | −0.343 | [−0.601, −0.085] | TRUE | TRUE |
| Jaccard nestedness-resultant | 1000. | −0.0729 | [−0.172, 0.022] | −0.325 | [−0.660, 0.009] | TRUE | FALSE |
| Jaccard nestedness-resultant | 1000. | 0.288 | [−0.079, 0.608] | 0.163 | [−0.557, 0.883] | TRUE | FALSE |
| Jaccard total dissimilarity | 1000. | −0.0631 | [−0.125, 0.004] | −0.161 | [−0.366, 0.044] | TRUE | FALSE |
| Jaccard total dissimilarity | 1000. | −0.0502 | [−0.119, 0.019] | −0.150 | [−0.353, 0.054] | TRUE | FALSE |
| Jaccard total dissimilarity | 1000. | −0.0643 | [−0.126, 0.002] | −0.141 | [−0.344, 0.062] | TRUE | FALSE |
| Jaccard total dissimilarity | 1000. | 0.0322 | [−0.052, 0.112] | −0.0367 | [−0.299, 0.226] | FALSE | FALSE |
| Jaccard total dissimilarity | 1000. | −0.00153 | [−0.270, 0.224] | 0.0444 | [−0.515, 0.604] | FALSE | FALSE |
| Jaccard turnover | 1000. | −0.00227 | [−0.083, 0.078] | 0.0443 | [−0.137, 0.226] | FALSE | FALSE |
| Jaccard turnover | 1000. | 0.0785 | [−0.007, 0.163] | 0.0821 | [−0.096, 0.260] | TRUE | FALSE |
| Jaccard turnover | 1000. | 0.0257 | [−0.060, 0.110] | 0.0941 | [−0.084, 0.273] | TRUE | FALSE |
| Jaccard turnover | 1000. | 0.0921 | [−0.005, 0.192] | 0.236 | [0.008, 0.464] | TRUE | FALSE |
| Jaccard turnover | 1000. | −0.0310 | [−0.284, 0.196] | 0.0186 | [−0.423, 0.460] | FALSE | FALSE |
| Sorensen nestedness-resultant | 1000. | −0.0610 | [−0.118, −0.008] | −0.314 | [−0.560, −0.068] | TRUE | TRUE |

| metric | n_success | median | Empirical 95% interval | main_estimate | Main-model 95% CI | direction_match | empirical_interval_excludes_zero |
|---|---|---|---|---|---|---|---|
| Sorensen nestedness-resultant | 1000. | −0.220 | [−0.296, −0.154] | −0.414 | [−0.659, −0.170] | TRUE | TRUE |
| Sorensen nestedness-resultant | 1000. | −0.0983 | [−0.160, −0.039] | −0.321 | [−0.564, −0.077] | TRUE | TRUE |
| Sorensen nestedness-resultant | 1000. | −0.0482 | [−0.144, 0.040] | −0.288 | [−0.604, 0.028] | TRUE | FALSE |
| Sorensen nestedness-resultant | 1000. | 0.277 | [−0.082, 0.571] | 0.205 | [−0.471, 0.882] | TRUE | FALSE |
| Sorensen total dissimilarity | 1000. | −0.0558 | [−0.120, 0.014] | −0.166 | [−0.368, 0.037] | TRUE | FALSE |
| Sorensen total dissimilarity | 1000. | −0.0498 | [−0.123, 0.021] | −0.152 | [−0.353, 0.049] | TRUE | FALSE |
| Sorensen total dissimilarity | 1000. | −0.0735 | [−0.142, −0.001] | −0.146 | [−0.347, 0.054] | TRUE | TRUE |
| Sorensen total dissimilarity | 1000. | 0.0357 | [−0.051, 0.119] | −0.0371 | [−0.297, 0.223] | FALSE | FALSE |
| Sorensen total dissimilarity | 1000. | 0.00305 | [−0.257, 0.227] | 0.0324 | [−0.521, 0.586] | TRUE | FALSE |
| Sorensen turnover | 1000. | 0.00331 | [−0.084, 0.089] | 0.0350 | [−0.145, 0.215] | TRUE | FALSE |
| Sorensen turnover | 1000. | 0.0798 | [−0.014, 0.171] | 0.0722 | [−0.104, 0.248] | TRUE | FALSE |
| Sorensen turnover | 1000. | 0.0215 | [−0.072, 0.113] | 0.0832 | [−0.093, 0.260] | TRUE | FALSE |
| Sorensen turnover | 1000. | 0.0958 | [−0.009, 0.202] | 0.228 | [0.002, 0.454] | TRUE | FALSE |
| Sorensen turnover | 1000. | −0.0188 | [−0.270, 0.215] | −0.00386 | [−0.444, 0.436] | TRUE | FALSE |

**Table S3b RQ2 Gaussian study cluster bootstrap**

| metric | Observed Gaussian-logit estimate | Bootstrap median | Bootstrap 95% lower | Bootstrap 95% upper | Two-sided bootstrap P | Successful resamples | Predictor |
|---|---|---|---|---|---|---|---|
| Bray-Curtis total dissimilarity | 0.0574 | 0.0670 | −0.377 | 0.613 | 0.768 | 1000. | Secondary vegetation x pair-mean anomaly |
| Bray-Curtis total dissimilarity | −0.166 | −0.166 | −0.480 | 0.127 | 0.204 | 1000. | Low-intensity agriculture x pair-mean anomaly |
| Bray-Curtis total dissimilarity | −0.0282 | 0.000800 | −0.234 | 0.320 | 0.988 | 1000. | High-intensity agriculture x pair-mean anomaly |
| Bray-Curtis total dissimilarity | 0.331 | 0.378 | −0.313 | 4.39 | 0.292 | 1000. | Urban x pair-mean anomaly |
| Bray-Curtis balanced variation | 0.280 | 0.263 | −0.122 | 0.651 | 0.170 | 1000. | Secondary vegetation x pair-mean anomaly |
| Bray-Curtis balanced variation | 0.0370 | 0.0390 | −0.298 | 0.387 | 0.826 | 1000. | Low-intensity agriculture x pair-mean anomaly |
| Bray-Curtis balanced variation | 0.0450 | 0.0504 | −0.207 | 0.413 | 0.700 | 1000. | High-intensity agriculture x pair-mean anomaly |
| Bray-Curtis balanced variation | 0.0202 | 0.112 | −0.614 | 6.46 | 0.784 | 1000. | Urban x pair-mean anomaly |

| metric | Observed Gaussian-logit estimate | Bootstrap median | Bootstrap 95% lower | Bootstrap 95% upper | Two-sided bootstrap P | Successful resamples | Predictor |
|---|---|---|---|---|---|---|---|
| Bray-Curtis abundance gradient | 0.0216 | 0.0408 | −0.405 | 0.529 | 0.826 | 1000. | Secondary vegetation x pair-mean anomaly |
| Bray-Curtis abundance gradient | 0.0675 | 0.0595 | −0.266 | 0.420 | 0.698 | 1000. | Low-intensity agriculture x pair-mean anomaly |
| Bray-Curtis abundance gradient | 0.0756 | 0.0497 | −0.322 | 0.421 | 0.762 | 1000. | High-intensity agriculture x pair-mean anomaly |
| Bray-Curtis abundance gradient | 0.101 | 0.0355 | −7.64 | 0.903 | 0.914 | 1000. | Urban x pair-mean anomaly |
| Jaccard total dissimilarity | 0.0415 | 0.0527 | −0.390 | 0.786 | 0.826 | 1000. | Secondary vegetation x pair-mean anomaly |
| Jaccard total dissimilarity | −0.0794 | −0.0979 | −0.456 | 0.295 | 0.600 | 1000. | Low-intensity agriculture x pair-mean anomaly |
| Jaccard total dissimilarity | −0.00610 | 0.0278 | −0.274 | 0.492 | 0.880 | 1000. | High-intensity agriculture x pair-mean anomaly |
| Jaccard total dissimilarity | 0.303 | 0.373 | −0.391 | 4.82 | 0.386 | 1000. | Urban x pair-mean anomaly |
| Jaccard turnover | 0.219 | 0.194 | −0.218 | 0.609 | 0.286 | 1000. | Secondary vegetation x pair-mean anomaly |
| Jaccard turnover | 0.0777 | 0.0592 | −0.307 | 0.463 | 0.804 | 1000. | Low-intensity agriculture x pair-mean anomaly |
| Jaccard turnover | −0.0725 | −0.0644 | −0.340 | 0.284 | 0.674 | 1000. | High-intensity agriculture x pair-mean anomaly |
| Jaccard turnover | −0.462 | −0.329 | −1.38 | 8.54 | 0.730 | 1000. | Urban x pair-mean anomaly |
| Jaccard nestedness-resultant | 0.0772 | 0.104 | −0.390 | 0.805 | 0.650 | 1000. | Secondary vegetation x pair-mean anomaly |
| Jaccard nestedness-resultant | 0.139 | 0.126 | −0.205 | 0.581 | 0.488 | 1000. | Low-intensity agriculture x pair-mean anomaly |
| Jaccard nestedness-resultant | 0.171 | 0.159 | −0.262 | 0.625 | 0.444 | 1000. | High-intensity agriculture x pair-mean anomaly |
| Jaccard nestedness-resultant | 0.702 | 0.640 | −4.07 | 1.44 | 0.646 | 1000. | Urban x pair-mean anomaly |
| Sorensen total dissimilarity | 0.0286 | 0.0421 | −0.433 | 0.799 | 0.878 | 1000. | Secondary vegetation x pair-mean anomaly |
| Sorensen total dissimilarity | −0.0968 | −0.115 | −0.499 | 0.287 | 0.538 | 1000. | Low-intensity agriculture x pair-mean anomaly |
| Sorensen total dissimilarity | −0.0172 | 0.0192 | −0.296 | 0.486 | 0.914 | 1000. | High-intensity agriculture x pair-mean anomaly |
| Sorensen total dissimilarity | 0.327 | 0.393 | −0.399 | 4.95 | 0.368 | 1000. | Urban x pair-mean anomaly |
| Sorensen turnover | 0.201 | 0.179 | −0.245 | 0.587 | 0.344 | 1000. | Secondary vegetation x pair-mean anomaly |
| Sorensen turnover | 0.0441 | 0.0270 | −0.327 | 0.429 | 0.916 | 1000. | Low-intensity agriculture x pair- |

| metric | Observed Gaussian-logit estimate | Bootstrap median | Bootstrap 95% lower | Bootstrap 95% upper | Two-sided bootstrap P | Successful resamples | Predictor |
|---|---|---|---|---|---|---|---|
| | | | | | | | mean anomaly |
| Sorensen turnover | −0.0883 | −0.0738 | −0.354 | 0.283 | 0.630 | 1000. | High-intensity agriculture x pair-mean anomaly |
| Sorensen turnover | −0.393 | −0.264 | −1.31 | 8.54 | 0.758 | 1000. | Urban x pair-mean anomaly |
| Sorensen nestedness-resultant | 0.104 | 0.134 | −0.378 | 0.833 | 0.578 | 1000. | Secondary vegetation x pair-mean anomaly |
| Sorensen nestedness-resultant | 0.161 | 0.141 | −0.180 | 0.583 | 0.420 | 1000. | Low-intensity agriculture x pair-mean anomaly |
| Sorensen nestedness-resultant | 0.173 | 0.159 | −0.251 | 0.627 | 0.430 | 1000. | High-intensity agriculture x pair-mean anomaly |
| Sorensen nestedness-resultant | 0.692 | 0.642 | −2.95 | 1.35 | 0.654 | 1000. | Urban x pair-mean anomaly |

**Table S3c RQ2 Gaussian bootstrap LOSO comparison**

| metric | Gaussian-logit estimate | Model 95% lower | Model 95% upper | Study-bootstrap 95% lower | Study-bootstrap 95% upper | LOSO minimum | LOSO maximum | Predictor |
|---|---|---|---|---|---|---|---|---|
| Bray-Curtis total dissimilarity | 0.0574 | −0.324 | 0.439 | −0.377 | 0.613 | −0.0879 | 0.169 | Secondary vegetation x pair-mean anomaly |
| Bray-Curtis total dissimilarity | −0.166 | −0.411 | 0.0791 | −0.480 | 0.127 | −0.234 | −0.0852 | Low-intensity agriculture x pair-mean anomaly |
| Bray-Curtis total dissimilarity | −0.0282 | −0.278 | 0.221 | −0.234 | 0.320 | −0.0845 | 0.0525 | High-intensity agriculture x pair-mean anomaly |
| Bray-Curtis total dissimilarity | 0.331 | −0.629 | 1.29 | −0.313 | 4.39 | 0.0242 | 3.09 | Urban x pair-mean anomaly |
| Jaccard total dissimilarity | 0.0415 | −0.394 | 0.477 | −0.390 | 0.786 | −0.0799 | 0.170 | Secondary vegetation x pair-mean anomaly |
| Jaccard total dissimilarity | −0.0794 | −0.362 | 0.204 | −0.456 | 0.295 | −0.195 | −0.00730 | Low-intensity agriculture x pair-mean anomaly |
| Jaccard total dissimilarity | −0.00610 | −0.293 | 0.281 | −0.274 | 0.492 | −0.120 | 0.0899 | High-intensity agriculture x pair-mean anomaly |
| Jaccard total dissimilarity | 0.303 | −0.800 | 1.41 | −0.391 | 4.82 | −0.0731 | 2.89 | Urban x pair-mean anomaly |
| Sorensen total dissimilarity | 0.0286 | −0.420 | 0.477 | −0.433 | 0.799 | −0.103 | 0.164 | Secondary vegetation x pair-mean anomaly |
| Sorensen total dissimilarity | −0.0968 | −0.388 | 0.194 | −0.499 | 0.287 | −0.218 | −0.0194 | Low-intensity agriculture x pair-mean anomaly |
| Sorensen total dissimilarity | −0.0172 | −0.312 | 0.278 | −0.296 | 0.486 | −0.130 | 0.0849 | High-intensity agriculture x pair-mean anomaly |
| Sorensen total dissimilarity | 0.327 | −0.806 | 1.46 | −0.399 | 4.95 | −0.0588 | 3.11 | Urban x pair-mean anomaly |

**Table S4 |** Leave-one-study-out stability of omnibus and coefficient-level RQ1 inference. The complete 518-refit table is supplied in the accompanying source-data workbook.

**Table S4a LOSO summary**

| Model | Attempted | Usable | Omnibus P < 0.05 | Maximum omnibus P | Low-intensity P < 0.05 | High-intensity P < 0.05 | Urban P < 0.05 |
|---|---|---|---|---|---|---|---|
| Total abundance, maximum-temperature anomaly | 121. | 121. | 121. | 0.042 | 121. | 1.00 | 120. |
| Total abundance, mean-temperature anomaly | 121. | 121. | 121. | 0.007 | 121. | 0 | 0 |
| Species richness, maximum-temperature anomaly | 138. | 138. | 138. | 1.6e-04 | 138. | 5.00 | 137. |
| Species richness, mean-temperature anomaly | 138. | 138. | 138. | 5.2e-10 | 138. | 138. | 136. |

**Table S5 |** Harmonised climate-product sensitivity for local biodiversity.

**Table S5a climate product omnibus**

| Response | Climate product | Common-sample n | Studies | Spatial blocks | Likelihood-ratio χ² | d.f. | P | Full-model positive-definite Hessian | Reduced-model positive-definite Hessian |
|---|---|---|---|---|---|---|---|---|---|
| Total abundance | CRU TS | 5750. | 121. | 494. | 20.2 | 4.00 | 4.5e-04 | TRUE | TRUE |
| Total abundance | TerraClimate | 5750. | 121. | 494. | 12.3 | 4.00 | 0.015 | TRUE | TRUE |
| Total abundance | ERA5-Land | 5750. | 121. | 494. | 36.0 | 4.00 | 2.9e-07 | TRUE | TRUE |
| Total abundance | CHELSA v2.1 | 5750. | 121. | 494. | 15.2 | 4.00 | 0.004 | TRUE | TRUE |
| Species richness | CRU TS | 6386. | 138. | 590. | 57.9 | 4.00 | 7.9e-12 | TRUE | TRUE |
| Species richness | TerraClimate | 6386. | 138. | 590. | 41.9 | 4.00 | 1.7e-08 | TRUE | TRUE |
| Species richness | ERA5-Land | 6386. | 138. | 590. | 85.1 | 4.00 | 1.5e-17 | TRUE | TRUE |
| Species richness | CHELSA v2.1 | 6386. | 138. | 590. | 63.8 | 4.00 | 4.6e-13 | TRUE | TRUE |

**Table S5b climate product coefficients**

**CRU TS | Total abundance**

| Predictor | Estimate | s.e. | 95% CI | z | P |
|---|---|---|---|---|---|
| Secondary vegetation x temperature anomaly | 0.161 | 0.118 | [−0.071, 0.393] | 1.36 | 0.173 |
| Low-intensity agriculture x temperature anomaly | 0.326 | 0.0875 | [0.155, 0.498] | 3.73 | 1.9e-04 |
| High-intensity agriculture x temperature anomaly | −0.0105 | 0.0878 | [−0.183, 0.161] | −0.120 | 0.904 |
| Urban x temperature anomaly | 0.235 | 0.288 | [−0.329, 0.800] | 0.817 | 0.414 |

**TerraClimate | Total abundance**

| Predictor | Estimate | s.e. | 95% CI | z | P |
|---|---|---|---|---|---|
| Secondary vegetation x temperature anomaly | 0.0644 | 0.106 | [−0.143, 0.271] | 0.610 | 0.542 |
| Low-intensity agriculture x temperature anomaly | 0.230 | 0.0889 | [0.056, 0.404] | 2.59 | 0.010 |
| High-intensity agriculture x temperature anomaly | −0.129 | 0.100 | [−0.325, 0.068] | −1.29 | 0.198 |
| Urban x temperature anomaly | 0.193 | 0.380 | [−0.553, 0.938] | 0.507 | 0.612 |

**ERA5-Land | Total abundance**

| Predictor | Estimate | s.e. | 95% CI | z | P |
|---|---|---|---|---|---|
| Secondary vegetation x temperature anomaly | 0.470 | 0.120 | [0.235, 0.706] | 3.92 | 8.8e-05 |
| Low-intensity agriculture x temperature anomaly | 0.571 | 0.105 | [0.365, 0.778] | 5.42 | 5.9e-08 |
| High-intensity agriculture x temperature anomaly | 0.361 | 0.117 | [0.132, 0.590] | 3.08 | 0.002 |
| Urban x temperature anomaly | −0.123 | 0.527 | [−1.157, 0.911] | −0.233 | 0.816 |

**CHELSA v2.1 | Total abundance**

| Predictor | Estimate | s.e. | 95% CI | z | P |
|---|---|---|---|---|---|
| Secondary vegetation x temperature anomaly | −0.0593 | 0.146 | [−0.344, 0.226] | −0.407 | 0.684 |
| Low-intensity agriculture x temperature anomaly | 0.0884 | 0.118 | [−0.143, 0.320] | 0.747 | 0.455 |
| High-intensity agriculture x temperature anomaly | −0.538 | 0.153 | [−0.839, −0.238] | −3.51 | 4.5e-04 |
| Urban x temperature anomaly | −0.0480 | 0.533 | [−1.092, 0.997] | −0.0900 | 0.928 |

**CRU TS | Species richness**

| Predictor | Estimate | s.e. | 95% CI | z | P |
|---|---|---|---|---|---|
| Secondary vegetation x temperature anomaly | −0.0157 | 0.0524 | [−0.118, 0.087] | −0.301 | 0.764 |
| Low-intensity agriculture x temperature anomaly | 0.172 | 0.0429 | [0.088, 0.256] | 4.01 | 6.1e-05 |

| Predictor | Estimate | s.e. | 95% CI | z | P |
|---|---|---|---|---|---|
| High-intensity agriculture x temperature anomaly | −0.154 | 0.0402 | [−0.233, −0.075] | −3.83 | 1.3e-04 |
| Urban x temperature anomaly | −0.404 | 0.160 | [−0.718, −0.090] | −2.52 | 0.012 |

**TerraClimate | Species richness**

| Predictor | Estimate | s.e. | 95% CI | z | P |
|---|---|---|---|---|---|
| Secondary vegetation x temperature anomaly | −0.0351 | 0.0508 | [−0.135, 0.065] | −0.691 | 0.490 |
| Low-intensity agriculture x temperature anomaly | 0.0746 | 0.0427 | [−0.009, 0.158] | 1.75 | 0.081 |
| High-intensity agriculture x temperature anomaly | −0.233 | 0.0464 | [−0.324, −0.142] | −5.02 | 5.3e-07 |
| Urban x temperature anomaly | −0.489 | 0.168 | [−0.819, −0.159] | −2.90 | 0.004 |

**ERA5-Land | Species richness**

| Predictor | Estimate | s.e. | 95% CI | z | P |
|---|---|---|---|---|---|
| Secondary vegetation x temperature anomaly | 0.221 | 0.0561 | [0.111, 0.331] | 3.94 | 8.1e-05 |
| Low-intensity agriculture x temperature anomaly | 0.382 | 0.0510 | [0.282, 0.482] | 7.48 | 7.4e-14 |
| High-intensity agriculture x temperature anomaly | −0.0891 | 0.0526 | [−0.192, 0.014] | −1.69 | 0.090 |
| Urban x temperature anomaly | −0.992 | 0.310 | [−1.601, −0.384] | −3.20 | 0.001 |

**CHELSA v2.1 | Species richness**

| Predictor | Estimate | s.e. | 95% CI | z | P |
|---|---|---|---|---|---|
| Secondary vegetation x temperature anomaly | −0.258 | 0.0697 | [−0.395, −0.122] | −3.70 | 2.1e-04 |
| Low-intensity agriculture x temperature anomaly | −0.0712 | 0.0564 | [−0.182, 0.039] | −1.26 | 0.207 |
| High-intensity agriculture x temperature anomaly | −0.527 | 0.0709 | [−0.666, −0.388] | −7.44 | 1.0e-13 |
| Urban x temperature anomaly | −0.781 | 0.263 | [−1.296, −0.266] | −2.97 | 0.003 |

**Table S6 |** Alternative count-family interaction tests and residual diagnostics.

**Table S6a alternative count family LRT**

| Response | Temperature metric | Response family | n | χ² | d.f. | P | Full model Hessian | Reduced model Hessian | Dispersion parameter |
|---|---|---|---|---|---|---|---|---|---|
| richness | mean | COM-Poisson | 6457. | 65.2 | 4.00 | 2.3e-13 | TRUE | TRUE | 1.49 |
| richness | maximum | COM-Poisson | 6457. | 38.2 | 4.00 | 1.0e-07 | TRUE | TRUE | 1.49 |
| richness | mean | generalized Poisson | 6457. | 72.1 | 4.00 | 8.2e-15 | TRUE | TRUE | 1.43 |
| richness | maximum | generalized Poisson | 6457. | 41.1 | 4.00 | 2.6e-08 | TRUE | TRUE | 1.43 |

**Table S6b COMPoisson diagnostics**

| Temperature metric | Response family | Uniformity statistic | Uniformity P | Dispersion statistic | Dispersion P | Outliers | Outlier P | Zero-inflation P | Quantile P |
|---|---|---|---|---|---|---|---|---|---|
| mean | COM-Poisson | 0.241 | 0.0e+00 | 0.0957 | 0 | 0 | 0.480 | 0.084 | 0 |
| maximum | COM-Poisson | 0.227 | 4.4e-290 | 0.0902 | 0 | 0 | 0.580 | 0.148 | 0 |

**Table S6c generalised Poisson diagnostics**

| Temperature metric | Response family | Uniformity statistic | Uniformity P | Dispersion statistic | Dispersion P | Outliers | Outlier P | Zero-inflation P | Quantile P |
|---|---|---|---|---|---|---|---|---|---|
| mean | generalized Poisson | 0.233 | 7.0e-306 | 0.116 | 0 | 0 | 0.460 | 0.092 | 0 |
| maximum | generalized Poisson | 0.225 | 2.3e-284 | 0.110 | 0 | 1.00 | 0.660 | 0.096 | 0 |

**Table S7 |** Within-study permutation tests of residual spatial autocorrelation.

**Table S7 spatial within study permutation**

| Model | Distance (km) | Studies | Pairs | Moran's I | Permutation P | BH-adjusted P |
|---|---|---|---|---|---|---|
| Total abundance, mean-temperature anomaly | 0–50 | 121. | 308834. | −0.00640 | 0.156 | 0.416 |
| Total abundance, mean-temperature anomaly | 50–200 | 121. | 50115. | −0.0139 | 0.179 | 0.416 |
| Total abundance, mean-temperature anomaly | 200–500 | 121. | 3062. | −0.000437 | 0.361 | 0.481 |
| Total abundance, mean-temperature anomaly | 500–2000 | 121. | 0 | | | |
| Total abundance, maximum-temperature anomaly | 0–50 | 121. | 308834. | −0.00639 | 0.134 | 0.416 |
| Total abundance, maximum-temperature anomaly | 50–200 | 121. | 50115. | −0.0140 | 0.152 | 0.416 |
| Total abundance, maximum-temperature anomaly | 200–500 | 121. | 3062. | −0.00142 | 0.435 | 0.522 |
| Total abundance, maximum-temperature anomaly | 500–2000 | 121. | 0 | | | |
| Species richness, mean-temperature anomaly | 0–50 | 138. | 333092. | −0.00837 | 0.295 | 0.477 |
| Species richness, mean-temperature anomaly | 50–200 | 138. | 60762. | −0.00458 | 0.318 | 0.477 |
| Species richness, mean-temperature anomaly | 200–500 | 138. | 3062. | −0.0120 | 0.984 | 0.984 |
| Species richness, mean-temperature anomaly | 500–2000 | 138. | 0 | | | |
| Species richness, maximum-temperature anomaly | 0–50 | 138. | 333092. | −0.00852 | 0.193 | 0.416 |
| Species richness, maximum-temperature anomaly | 50–200 | 138. | 60762. | −0.00396 | 0.208 | 0.416 |
| Species richness, maximum-temperature anomaly | 200–500 | 138. | 3062. | −0.0116 | 0.966 | 0.984 |
| Species richness, maximum-temperature anomaly | 500–2000 | 138. | 0 | | | |

**Table S8 |** Structure of the warming-exposure gradient within and among PREDICTS studies, for each analysis sample and warming metric. Within-study span is the range of the standardised anomaly among sites of the same study. Two medians are given because they answer different questions: the per-study median treats every study equally, whereas the site-weighted median describes the span experienced by a typical site. Variance components are from an intercept-only linear mixed model with a study random intercept. Per-study spans for all 138 studies are given in the source data.

**Table S8 warming exposure support**

| Analysis sample | Warming metric | Sites | Studies | Studies with zero within-study span | Studies with zero span (%) | Median within-study span (per study) | Median within-study span (site-weighted) | 75th percentile of within-study span | Maximum within-study span | Among-study variance | Within-study variance |
|---|---|---|---|---|---|---|---|---|---|---|---|
| Species-richness sample | Mean-temperature anomaly | 6457. | 138. | 58.0 | 42.0 | 0.00610 | 0.0151 | 0.0368 | 0.887 | 0.187 | 0.00320 |
| Species-richness sample | Maximum-temperature anomaly | 6457. | 138. | 62.0 | 44.9 | 0.0442 | 0.165 | 0.366 | 8.30 | 3.12 | 0.184 |
| Total-abundance sample | Mean-temperature anomaly | 5740. | 121. | 52.0 | 43.0 | 0.00420 | 0.0176 | 0.0439 | 0.887 | 0.178 | 0.00360 |
| Total-abundance sample | Maximum-temperature anomaly | 5740. | 121. | 56.0 | 46.3 | 0.0514 | 0.190 | 0.394 | 8.30 | 2.77 | 0.197 |
| All sites with anomalies | Mean-temperature anomaly | 6457. | 138. | 58.0 | 42.0 | 0.00610 | 0.0151 | 0.0368 | 0.887 | 0.187 | 0.00320 |
| All sites with anomalies | Maximum-temperature anomaly | 6457. | 138. | 62.0 | 44.9 | 0.0442 | 0.165 | 0.366 | 8.30 | 3.12 | 0.184 |

**Table S9 |** Reconciliation of the trait-imputation and occurrence-model species denominators.

**Table S9 trait and occurrence species denominators**

| Analysis stage | Quantity | n | Note |
|---|---|---|---|
| 1. Raw ecological-characteristic table | Species compiled | 2946 | All bird species matched to PREDICTS records and to the ecological-characteristic sources |
| 1. Raw ecological-characteristic table | Trait cells (8 per species) | 23568 | 2,946 species x 8 ecological characteristics |
| 2. Missing values | Realised thermal niche breadth missing | 113 | 3.84% of 2,946 species |
| 2. Missing values | Diet breadth missing | 63.0 | 2.14% of 2,946 species |
| 2. Missing values | Trait cells imputed | 176 | 0.75% of 23,568 cells; |
| 2. Missing values | Species with ≥1 imputed value | 114 | 3.87% of 2,946 species; 62 species were missing both ecological characteristics |
| 3. Species retained for clustering | Species retained for clustering after imputation | 2946 | These define the primary eight-characteristic ecological groups |
| 4. Ecological groups | EG1 | 612 | Primary eight-characteristic k-means K = 3 solution |
| 4. Ecological groups | EG2 | 954 | Primary eight-characteristic k-means K = 3 solution |
| 4. Ecological groups | EG3 | 1380 | Primary eight-characteristic k-means K = 3 solution |
| 5. Occurrence model | Species entering the fitted model | 2946 | Equals the sum of EG1-EG3 |
| 5. Occurrence model | Species-site records | 432560. | 6,357 sites x study-specific species pools |

**Table S10 |** Clustering-solution screening, principal-component loadings, explained variance and ecological-group profiles.

**Table S10 cluster solution screening**

| method | K | silhouette | min_prop | mean_cluster_jaccard | min_cluster_jaccard | final_candidate |
|---|---|---|---|---|---|---|
| kmeans | 3.00 | 0.215 | 0.394 | 0.607 | 0.603 | TRUE |
| kmeans | 2.00 | 0.192 | 0.207 | 0.592 | 0.588 | FALSE |
| kmeans | 5.00 | 0.203 | 0.074 | 0.582 | 0.564 | FALSE |
| pam | 5.00 | 0.167 | 0.129 | 0.411 | 0.327 | FALSE |
| ward | 5.00 | 0.147 | 0.089 | 0.305 | 0.264 | FALSE |
| kmeans | 4.00 | 0.199 | 0.154 | 0.549 | 0.529 | FALSE |
| pam | 4.00 | 0.175 | 0.186 | 0.515 | 0.491 | FALSE |
| ward | 4.00 | 0.153 | 0.109 | 0.322 | 0.266 | FALSE |
| pam | 3.00 | 0.207 | 0.250 | 0.528 | 0.499 | FALSE |
| ward | 3.00 | 0.156 | 0.176 | 0.364 | 0.309 | FALSE |
| pam | 2.00 | 0.270 | 0.333 | 0.539 | 0.520 | FALSE |
| ward | 2.00 | 0.166 | 0.359 | 0.418 | 0.407 | FALSE |

**Table S11 |** Sensitivity to adjustment for the site-level 1901–1930 climatological mean annual temperature.

**Table S11a static climate omnibus**

| Response |
|---|
| Total abundance |
| Species richness |

**Table S11b static climate coefficients**

| Response | Predictor |
|---|---|
| Species richness | Intercept (primary vegetation at anomaly = 0) |
| Species richness | Standardised mean-temperature anomaly |
| Species richness | High-intensity agriculture |
| Species richness | High-intensity agriculture × standardised mean-temperature anomaly |
| Species richness | Low-intensity agriculture |
| Species richness | Low-intensity agriculture × standardised mean-temperature anomaly |
| Species richness | Secondary vegetation |
| Species richness | Secondary vegetation × standardised mean-temperature anomaly |
| Species richness | Urban |
| Species richness | Urban × standardised mean-temperature anomaly |
| Species richness | Baseline (1901–1930) mean annual temperature, standardised |
| Total abundance | Intercept (primary vegetation at anomaly = 0) |
| Total abundance | Standardised mean-temperature anomaly |
| Total abundance | High-intensity agriculture |
| Total abundance | High-intensity agriculture × standardised mean-temperature anomaly |
| Total abundance | Low-intensity agriculture |
| Total abundance | Low-intensity agriculture × standardised mean-temperature anomaly |
| Total abundance | Secondary vegetation |
| Total abundance | Secondary vegetation × standardised mean-temperature anomaly |
| Total abundance | Urban |
| Total abundance | Urban × standardised mean-temperature anomaly |
| Total abundance | Baseline (1901–1930) mean annual temperature, standardised |

**Table S12 |** Sensitivity of occurrence-model response-scale decomposition to the spatial-block random intercept.

**Table S12 RQ3 spatial block sensitivity**

| Ecological group | Land use | Without spatial block estimate | Without spatial block 95% CI | With spatial block estimate | With spatial block 95% CI | Classification changed |
|---|---|---|---|---|---|---|
| EG2 | Primary vegetation | 0 | [0.000, 0.000] | 0 | [0.000, 0.000] | FALSE |
| EG2 | Secondary vegetation | 4.85 | [1.307, 8.577] | 5.28 | [1.657, 8.711] | FALSE |
| EG2 | Low-intensity agriculture | 13.6 | [10.431, 17.121] | 14.5 | [11.186, 18.193] | FALSE |
| EG2 | High-intensity agriculture | 6.10 | [2.957, 9.214] | 5.94 | [2.776, 9.098] | FALSE |
| EG2 | Urban | −5.39 | [−13.904, 4.121] | −5.39 | [−14.058, 4.040] | FALSE |
| EG1 | Primary vegetation | 0 | [0.000, 0.000] | 0 | [0.000, 0.000] | FALSE |
| EG1 | Secondary vegetation | 0.225 | [−4.684, 5.242] | 0.883 | [−4.024, 6.129] | FALSE |
| EG1 | Low-intensity agriculture | −5.39 | [−9.703, −1.205] | −4.46 | [−9.008, −0.019] | FALSE |
| EG1 | High-intensity agriculture | −4.61 | [−8.758, −0.616] | −4.84 | [−9.203, −0.615] | FALSE |
| EG1 | Urban | 20.6 | [6.285, 35.932] | 20.4 | [6.738, 35.477] | FALSE |
| EG3 | Primary vegetation | 0 | [0.000, 0.000] | 0 | [0.000, 0.000] | FALSE |
| EG3 | Secondary vegetation | −1.14 | [−3.841, 1.507] | −0.929 | [−3.487, 1.701] | FALSE |
| EG3 | Low-intensity agriculture | 3.93 | [1.325, 6.371] | 4.45 | [1.697, 7.122] | FALSE |
| EG3 | High-intensity agriculture | −3.93 | [−6.704, −1.172] | −4.57 | [−7.606, −1.621] | FALSE |
| EG3 | Urban | 21.5 | [8.286, 35.158] | 20.6 | [6.782, 33.681] | FALSE |

**Table S13 |** Simulation-based residual diagnostics for the four primary local-diversity models.

**Table S13 primary family diagnostics**

| Model | Observations | Simulations | Uniformity statistic | Uniformity P | Dispersion statistic | Dispersion P | Outliers | Outlier P | Zero-inflation P | Quantile P |
|---|---|---|---|---|---|---|---|---|---|---|
| Total abundance, mean-temperature anomaly | 5817. | 500. | 0.180 | 3.5e-164 | 0.933 | 0.620 | 225. | 4.5e-138 | | 0 |
| Total abundance, maximum-temperature anomaly | 5817. | 500. | 0.174 | 5.1e-153 | 0.946 | 0.708 | 211. | 3.1e-124 | | 0 |
| Species richness, mean-temperature anomaly | 6457. | 500. | 0.232 | 3.7e-301 | 0.0874 | 0.0e+00 | 54.0 | 9.1e-07 | 0.028 | 0 |
| Species richness, maximum-temperature anomaly | 6457. | 500. | 0.225 | 9.5e-285 | 0.0834 | 0.0e+00 | 31.0 | 0.278 | 0.052 | 0 |

**Table S14 |** Residual offsets across land-use classes and temperature-anomaly bins.

**Table S14 residual pattern by land use and mean temperature anomaly**

**Land-use strata**

| Model | n | Primary | Secondary | Low-intensity | High-intensity | Urban | Range | Maximum absolute offset |
|---|---|---|---|---|---|---|---|---|
| Total abundance, mean-temperature anomaly | 5817. | 0.434 | 0.414 | 0.450 | 0.480 | 0.362 | 0.118 | 0.138 |
| Species richness, mean-temperature anomaly | 6457. | 0.361 | 0.398 | 0.396 | 0.340 | 0.339 | 0.0590 | 0.161 |
| Total abundance, maximum-temperature anomaly | 5817. | 0.422 | 0.402 | 0.456 | 0.498 | 0.334 | 0.164 | 0.166 |
| Species richness, maximum-temperature anomaly | 6457. | 0.344 | 0.390 | 0.401 | 0.340 | 0.315 | 0.0850 | 0.185 |

**Temperature-anomaly gradient**

| Model | n | Lowest anomaly quintile | Highest anomaly quintile | Spearman rho | Spearman P |
|---|---|---|---|---|---|
| Total abundance, mean-temperature anomaly | 5817. | 0.427 | 0.490 | 0.00500 | 0.702 |
| Species richness, mean-temperature anomaly | 6457. | 0.410 | 0.506 | 0.113 | 9.3e-20 |
| Total abundance, maximum-temperature anomaly | 5817. | 0.428 | 0.508 | 0.0823 | 3.3e-10 |
| Species richness, maximum-temperature anomaly | 6457. | 0.395 | 0.538 | 0.172 | 3.9e-44 |

**Table S15 |** Intraclass correlation coefficients and variance explained for every fitted mixed model. ICC (analytic) divides each random-effect variance by the sum of all random-effect variances and the distribution-specific variance (residual variance for Gaussian models, $\pi^2/3$ for binomial models, and $\log(1 + 1/\mu + 1/\theta)$ for negative-binomial models). ICC (adjusted) is the corresponding value returned by performance::icc. Marginal and conditional $R^2$ follow Nakagawa et al. and are reported once per model. For the negative-binomial richness models the conditional $R^2$ approaches its upper bound because among-study variance is very large relative to the conditional mean, and it should be read alongside the ICC rather than as an independent goodness-of-fit summary.

**Table S15 ICC and variance explained**

| Model | Response family | Random effect | icc_analytic | icc_adjusted | r2_marginal | r2_conditional |
|---|---|---|---|---|---|---|
| Total abundance, mean-temperature anomaly | Gaussian | Study | 0.527 | 0.527 | 0.0192 | 0.595 |
| Total abundance, mean-temperature anomaly | Gaussian | Spatial block within study | 0.0605 | 0.0605 | 0.0192 | 0.595 |
| Total abundance, maximum-temperature anomaly | Gaussian | Study | 0.528 | 0.528 | 0.0237 | 0.601 |
| Total abundance, maximum-temperature anomaly | Gaussian | Spatial block within study | 0.0630 | 0.0630 | 0.0237 | 0.601 |
| Species richness, mean-temperature anomaly | Negative binomial | Study | 0.878 | 0.960 | 0.0103 | 1.00 |
| Species richness, mean-temperature anomaly | Negative binomial | Spatial block within study | 0.0366 | 0.0400 | 0.0103 | 1.00 |
| Species richness, maximum-temperature anomaly | Negative binomial | Study | 0.880 | 0.959 | 0.0138 | 1.00 |
| Species richness, maximum-temperature anomaly | Negative binomial | Spatial block within study | 0.0377 | 0.0410 | 0.0138 | 1.00 |
| Species-level occurrence (with spatial block) | Binomial | Study | 0.327 | 0.327 | 0.0163 | 0.574 |
| Species-level occurrence (with spatial block) | Binomial | Spatial block within study | 0.0133 | 0.0133 | 0.0163 | 0.574 |
| Species-level occurrence (with spatial block) | Binomial | Site within block | 0.0165 | 0.0165 | 0.0163 | 0.574 |
| Species-level occurrence (with spatial block) | Binomial | Species | 0.210 | 0.210 | 0.0163 | 0.574 |
| Bray-Curtis abundance gradient | Ordered beta | Study | | 0.521 | 0.0188 | 0.941 |
| Bray-Curtis abundance gradient | Ordered beta | Site i | | 0.207 | 0.0188 | 0.941 |
| Bray-Curtis abundance gradient | Ordered beta | Site j | | 0.212 | 0.0188 | 0.941 |
| Bray-Curtis total dissimilarity | Ordered beta | Study | | 0.571 | 0.0221 | 0.991 |
| Bray-Curtis total dissimilarity | Ordered beta | Site i | | 0.213 | 0.0221 | 0.991 |
| Bray-Curtis total dissimilarity | Ordered beta | Site j | | 0.208 | 0.0221 | 0.991 |
| Bray-Curtis balanced variation | Ordered beta | Study | | 0.736 | 0.00883 | 0.972 |
| Bray-Curtis balanced variation | Ordered beta | Site i | | 0.120 | 0.00883 | 0.972 |
| Bray-Curtis balanced variation | Ordered beta | Site j | | 0.116 | 0.00883 | 0.972 |
| Jaccard nestedness-resultant | Ordered beta | Study | | 0.500 | 0.0211 | 0.954 |
| Jaccard nestedness-resultant | Ordered beta | Site i | | 0.221 | 0.0211 | 0.954 |
| Jaccard nestedness-resultant | Ordered beta | Site j | | 0.232 | 0.0211 | 0.954 |
| Jaccard total dissimilarity | Ordered beta | Study | | 0.540 | 0.0185 | 0.995 |
| Jaccard total dissimilarity | Ordered beta | Site i | | 0.225 | 0.0185 | 0.995 |
| Jaccard total dissimilarity | Ordered beta | Site j | | 0.230 | 0.0185 | 0.995 |
| Jaccard turnover | Ordered beta | Study | | 0.752 | 0.0128 | 0.983 |
| Jaccard turnover | Ordered beta | Site i | | 0.114 | 0.0128 | 0.983 |
| Jaccard turnover | Ordered beta | Site j | | 0.117 | 0.0128 | 0.983 |
| Sorensen nestedness-resultant | Ordered beta | Study | | 0.505 | 0.0201 | 0.956 |
| Sorensen nestedness-resultant | Ordered beta | Site i | | 0.218 | 0.0201 | 0.956 |
| Sorensen nestedness-resultant | Ordered beta | Site j | | 0.231 | 0.0201 | 0.956 |
| Sorensen total dissimilarity | Ordered beta | Study | | 0.540 | 0.0197 | 0.989 |
| Sorensen total dissimilarity | Ordered beta | Site i | | 0.223 | 0.0197 | 0.989 |
| Sorensen total dissimilarity | Ordered beta | Site j | | 0.226 | 0.0197 | 0.989 |

| Model | Response family | Random effect | icc_analytic | icc_adjusted | r2_marginal | r2_conditional |
|---|---|---|---|---|---|---|
| Sorensen turnover | Ordered beta | Study | | 0.741 | 0.0127 | 0.964 |
| Sorensen turnover | Ordered beta | Site i | | 0.110 | 0.0127 | 0.964 |
| Sorensen turnover | Ordered beta | Site j | | 0.113 | 0.0127 | 0.964 |

**Table S16 |** Crosswalk from principal manuscript claims to supplementary evidence and interpretation limits.

**Table S16 manuscript evidence map**

| manuscript_claim | primary_evidence | interpretation_boundary |
|---|---|---|
| RQ1 mean-temperature land-use interaction | Table S1; Fig. 3 | Omnibus interaction supported; coefficient-level support varies |
| Maximum-temperature sensitivity | Table S1; Extended Data Fig. 2 | Complementary exposure analysis; not a uniform replication of mean-temperature coefficients |
| RQ2 bounded dissimilarity models | Table S2; Fig. 4; Extended Data Fig. 3 | Exact 0 and 1 retained; no global land-use-by-warming interaction among the nine responses |
| RQ2 shared-site dependence | Table S3; Supplementary Figs. S1 and S2 | Disjoint-pair subsampling addresses repeated site use; study resampling addresses study representation |
| RQ3 ecological-group definition | Tables S9 and S10; Extended Data Fig. 4 | Trait pool and occurrence-model species pool are different denominators |
| Climate-product sensitivity | Table S5; Extended Data Figs. 5b and 6; Supplementary Figs. S4 and S11 | Omnibus conclusion stable; some coefficients vary with product, baseline and resolution |
| Static-climate adjustment | Table S11; Supplementary Fig. S7 | Long-term mean annual temperature is an additive confounder adjustment |
| Study influence | Table S4; Extended Data Fig. 5a; Supplementary Fig. S3 | No single study drives the omnibus RQ1 interaction |
| Response-distribution diagnostics | Tables S6, S13 and S14; Extended Data Fig. 5c; Supplementary Figs. S5 and S8-S10 | Primary count model retained because focal inference is stable across families, despite imperfect residual fit |
| Residual spatial autocorrelation | Table S7; Extended Data Fig. 5d | No evidence within studies at distances up to 500 km; no eligible pairs at 500–2000 km |
